# A Non-canonical Role for Hepatocyte MLKL in Promoting Mitochondrial Dysfunction and Senescence in the Aging Liver

**DOI:** 10.1101/2025.05.02.651782

**Authors:** Sabira Mohammed, Chao Jiang, Travis Pennington, Shylesh Bhaskaran, Phoebe Ohene-Marfo, Constantin Georgescu, Kamille Pitts, Albert Tran, Zongkai Peng, Amit Singh, Zhibo Yang, Tiangang Li, Bethany Hannafon, Courtney Houchen, Jonathan D Wren, Veronica Galvan, Nagib Ahsan, Michael Kinter, Tommy L. Lewis, Sathyaseelan S. Deepa

## Abstract

Liver aging is characterized by chronic inflammation and metabolic dysfunction that contributes to the progression of metabolic dysfunction-associated steatotic liver disease (MASLD). Necroptosis, a form of inflammatory cell death, is activated in aging livers, and genetic (*Ripk3^-/-^*or *Mlkl^-/-^* mice) or pharmacological (RIPK1 inhibitor necrostatin-1s) inhibition of necroptosis attenuates liver inflammation and pathology. However, the cell type-specific role of necroptosis in liver aging remains unclear. Given that MLKL is expressed in hepatocytes, and its expression increases with age, we generated hepatocyte-specific MLKL-overexpressing mice (MLKL^HepOE^) to determine its role in liver aging. Unexpectedly, MLKL overexpression in hepatocytes did not induce necroptosis, but instead upregulated markers of cellular senescence (cell cycle arrest genes and SASP factors), increased macrophage infiltration, and elevated M1 macrophage marker expression. Electron microscopy and mitochondrial analyses revealed abnormal mitochondrial morphology, elevated oxidative stress, and disrupted mitochondrial dynamics, while lipidomics demonstrated alterations in hepatic lipid metabolites. In agreement with our observations in MLKL^HepOE^ mice, MLKL overexpression in AML12 hepatocytes impaired mitochondrial respiration, increased proinflammatory extracellular vesicle (EV) release, and induced senescence markers, without triggering cell death. Together, these findings reveal a non-lethal, non-necroptotic role for MLKL in promoting hepatocyte senescence and metabolic dysfunction via mitochondrial impairment and EV-mediated inflammation. Our study highlights MLKL as a novel driver of liver inflammaging and a potential therapeutic target for age-related liver disease.

## 1 Introduction

Aging is characterized by a progressive decline in cellular function, metabolic dysregulation, and a chronic low-grade inflammatory state known as inflammaging (Franceschi et al. 2000). This persistent inflammation is a hallmark of aging and is associated with various age-associated diseases, including neurodegeneration, cardiovascular disease, and metabolic disorders. Several mechanisms drive inflammaging, including cellular and immune senescence, alterations in gut microbiota, coagulation changes, and the release of damage-associated molecular patterns (DAMPs) (Franceschi and Campisi 2014). Among these, DAMPs play a key role in sustaining chronic inflammation in aging (Goldberg and Dixit 2015).

Necroptosis, a regulated form of inflammatory cell death, is a major source of DAMP release. Triggered by signals such as tumor necrosis factor α (TNFα), necroptosis is mediated by receptor-interacting protein kinases RIPK1 and RIPK3 and the terminal effector mixed lineage kinase domain-like pseudokinase (MLKL). Upon TNFα stimulation, RIPK1 autophosphorylates and activates RIPK3, which phosphorylates MLKL. Activated MLKL oligomerizes and translocate to the plasma membrane, disrupting membrane integrity and releasing DAMPs (e.g., HMGB1, ATP, S100 proteins). These DAMPs activate toll-like receptors (TLRs) and NF-κB signaling in innate immune cells, amplifying inflammation and creating a feed-forward loop that further promotes necroptosis (Pasparakis and Vandenabeele 2015). Activation of necroptosis is reported in several age-associated diseases, including neurodegenerative, cardiovascular, chronic kidney, and liver diseases such as MASLD (Yuan et al. 2019; Zhe-Wei et al. 2018; Kolbrink et al. 2023; Hanus et al. 2015; Luedde et al. 2014).

Liver aging is associated with increased inflammatory cytokine levels (Stahl et al. 2020) and chronic liver diseases such as MASLD (Saeed and Jun 2014). Since MASLD prevalence rises with age and increases the risk for cirrhosis and hepatocellular carcinoma (HCC), understanding its underlying mechanisms is critical. DAMPs released from damaged hepatocytes have been implicated in chronic liver inflammation (Koyama and Brenner 2017), and necroptosis has emerged as a key mode of cell death in MASLD/metabolic dysfunction-associated steatohepatitis (MASH) under pathological conditions (Saeed and Jun 2014; Dara et al. 2016). Inhibiting RIPK1, RIPK3, or MLKL reduces hepatic inflammation and fibrosis in mouse models (Majdi et al. 2020; Gautheron et al. 2014; Wu et al. 2020), highlighting necroptosis as a driver of liver pathology. We previously showed that markers of necroptosis (p-MLKL and MLKL oligomer) and inflammation are increased in the livers of *Sod1^-/-^*mice, a model of accelerated aging, and that pharmacological inhibition of RIPK1 with necrostatin-1s is sufficient to reduce liver inflammation and fibrosis in these mice (Mohammed, Nicklas, et al. 2021). Additionally, we found that necroptosis markers in the liver increase with age in wild-type mice and are positively correlated with elevated liver inflammation, steatosis, and fibrosis. Notably, inhibiting necroptosis, either genetically (*Ripk3^-/-^* or *Mlkl^-/-^*mice) or pharmacologically (with necrostatin-1s), significantly reduced liver inflammation, steatosis, and fibrosis (Mohammed, Thadathil, et al. 2021; Mohammed et al. 2025). These findings strongly suggest a role for necroptosis in liver inflammaging and pathology. However, the specific contribution of the liver or individual liver cell types remains unclear, as these studies used whole-body knockout models. Thus, inflammation in other tissues could also influence liver pathology.

To address this question, and considering that hepatocytes make up ∼80% of liver volume (Blouin et al. 1977), we developed a hepatocyte-specific MLKL overexpression (MLKL^HepOE^) mouse model. Our rationale for targeting MLKL was three-fold: (1) it is the terminal effector of necroptosis; (2) overexpression of the N-terminal domain of MLKL is sufficient to induce cell death even without necroptotic stimuli or RIPK3 (Dondelinger et al. 2014); and (3) MLKL protein is significantly upregulated (2–3 fold) in aging livers and hepatocytes, and in pathological conditions, correlating with inflammation and fibrosis (Mohammed, Nicklas, et al. 2021; Mohammed et al. 2025; Mohammed, Thadathil, et al. 2021; Ohene-Marfo et al. 2024). Surprisingly, we found that MLKL overexpression in hepatocytes does not induce necroptosis. Instead, high levels of MLKL caused mitochondrial dysfunction, upregulation of senescence markers, and increased hepatic inflammation, revealing a previously unrecognized, non-necroptotic role for MLKL in driving liver aging and pathology.

## 2 Methods

The detailed methodology is provided in the supplementary section.

### 2.1 Experimental Animals

All animal procedures were approved by the Institutional Animal Care and Use Committee at the University of Oklahoma Health Sciences Center (OUHSC). The mice were group housed in ventilated cages at 20°C ± 2°C, on a 12-hour dark/light cycle. Only male mice were used for the study, and mice were fed with Teklad 7013 NIH-31 Rodent Chow (Repelleted) diet (#C15101i, Research Diets) during the study.

### 2.2 Generation of Rosa26-MLKL and MLKL^HepOE^ mice

Rosa26-MLKL conditional knock in mouse model was generated by ViewSolid Biotech (Oklahoma, USA) by knocking in Mlkl cDNA (Mlkl 202 cDNA, Dharmacon reagents, Horizon Discovery) into the Rosa26 locus of mice in C57BL/6J background through CRISPR/Cas9-mediated homology-directed repair (**Figure S1A**). The conditional expression of the transgene was enabled by a Cre mediated deletion of the lox-P flanked STOP cassette in the tissue/organ of choice by using a tissue/organ specific Cre recombinase expression system. We generated MLKL^HepOE^ mice by injecting AAV8-TBG-Cre (2×10^11 Genomic Copies/mouse, VB1724, Vector Biolabs, PA, USA) virus, and for the control mice AAV8-TBG-Null (2×10^11 Genomic Copies/mouse, 70600-Pre, Vector Biolabs) virus through the tail vein of the Rosa26-MLKL mice (Kiourtis et al. 2021).

### 2.3 Isolation of hepatocytes and non-parenchymal cell (NPC) fraction

Hepatocytes and NPC fraction was isolated as described before (Mohammed, Thadathil, et al. 2021; Liu et al. 2017; Mohar et al. 2015).

### 2.4 Western Blotting

Western blotting was performed as previously described (Mohammed et al. 2025).

### 2.5 Detection of 4-Hydroxynonenal (4-HNE) adducts

4-HNE modified proteins were detected as described previously (Uchida et al. 1995; Mohammed, Nicklas, et al. 2021).

### 2.6 Immunofluorescence (IF) staining

The IF staining was done as previously described (Mohammed, Thadathil, et al. 2021).

### 2.7 Quantitative real-time PCR (qPCR)

RNA was isolated from frozen liver tissues, and qPCR was performed as described previously. (Ohene-Marfo et al. 2024). **Table S1** lists the primers used for the study.

### 2.8 Histological analysis of liver sections

Formalin-fixed and paraffin-embedded liver tissue sections were stained with Hematoxylin & Eosin (H&E) using the standardized protocol at the Stephenson Cancer Center Tissue Pathology core at the OUHSC. Images were taken using an ECHO REVOLVE R4 microscope (Discover Echo Inc). 3 random non-overlapping fields per sample were recorded.

### 2.9 Liver triglyceride quantification

Liver triglyceride levels were quantified using a triglyceride colorimetric assay kit (#10010303, Cayman Chemical Company) following the manufacturer’s instructions, using 50 mg of liver tissue as we have described (Mohammed et al. 2025).

### 2.10 SPiDER-βGal assay

SPiDER-βGal assay for senescence was performed using SPiDER-βGal assay kit (#SG02-10, Dojindo) as per manufacture’s instruction. Images were taken with Leica SP8 confocal microscope at 630 magnification, 10 random non-overlapping fields per sample were recorded.

### 2.11 Transmission electron microscopy

The electron microscopy experiment was carried out at the Oklahoma Medical Research Foundation Imaging Facility using established procedures, as described before (Bhaskaran et al. 2018). For analysis of mitochondria area, five sections/mouse were taken, and the area quantified using ImageJ software.

### 2.12 TUNEL assay

The DeadEnd Fluorometric TUNEL System (#G3250, Promega) was used to detect and quantify cell death at the single-cell level, following manufacturer’s instruction. The sections were mounted using ProLong™ Diamond antifade mountant with DAPI. Images were taken with Leica SP8 confocal microscope at 200x magnification, 4 random non-overlapping fields per sample were analyzed for TUNEL positive cells.

### 2.13 ELISA for Alanine Transaminase (ALT) and Aspartate Aminotransferase (AST)

The plasma ALT and AST levels were quantified using ALT or AST colorimetric activity assay kits (#700260, or #701640, Cayman Chemical Company), following the manufacturer’s instructions.

### 2.14 Human plasma MLKL analysis

Informed consent was obtained from all subjects involved in the study. Patients were included in this study if they had known MASH of any etiology based on laboratory, radiologic or histologic features. The exclusion criteria included age less than 18 years, pregnancy and any other known or suspected cancers. Patient samples were purchased from DX Biosamples, LLC including 20 control patients without known liver disease, and 20 patients with MASH (**Table S2**). The plasma MLKL levels were quantified using human mixed lineage kinase domain-like protein ELISA Kit (#MBS9300811, MyBioSource), following the manufacturer’s instructions.

### 2.15 Label free quantitative proteomic analysis

A total of 100 µg of liver lysate (n=5/group) was used in the analysis as described previously (Mohammed et al. 2025; Ahsan et al. 2023). Proteomics data can be found in the MassIVE database via MSV000097694.

### 2.16 Targeted mitochondrial proteomics

Quantitative targeted proteomics was performed to assess changes in mitochondrial enzyme levels in liver tissue, as previously described (Ohene-Marfo et al. 2024).

### 2.17 Untargeted lipidomics analysis

Untargeted lipidomics was performed by Creative Proteomics (Shirley, NY, USA) using UPLC-MS to profile liver lipid composition.

### 2.18 Overexpression of MLKL in AML12 cells

AML12 cells (#CRL2254, ATCC) were transfected with cDNA (pcDNA3.1+C-DYK, GenScript) or pcDNA-MLKL-Flag (ORF Clone, #OMU41768D, GenScript) using Lipofectamine reagent (#L3000008, Invitrogen). Experiments were performed 24 hour post transfection.

### 2.19 Cell viability assay

Cell viability was measured using Cell Counting kit-8 (CCK-8) (#96992, Sigma-Aldrich) assay according to manufacturer’s instruction, and absorbance was read at 450nm (Bio Tek Synergy H1).

### 2.20 Measurement of mitochondrial respiration

Mitochondrial respiration in intact cells were measured using Agilent Seahorse XF96 Extracellular Flux Analyzer (Agilent, CA, USA) as we have described (Deepa et al. 2016).

### 2.21 Live-cell imaging using photoactivatable GFP and LAMP1/mt-mScarlet reporters

AML12 cells were transfected with DDK tagged MLKL (pCAG MLKL-DDK) and fluorescent reporter (pCAG 2xmt-paGFP p2a 2xmt-mScarlet, pCAG 2xmt-mScarlet or pCAG Lamp1-mEmerald). Following overexpression, live time lapse imaging of the cells was done. For photo-activation experiments, 5µm by 5µm ROIs were selected in each cell imaged. For analysis of mitochondrial dynamics, a square ROI of 5µm by 5µm was used and ROI intensity statistics were quantified across the time course. Data was plotted over the entire time course, with comparisons made with end points and slopes.

### 2.22 Exosome isolation and nanoparticle tracking analysis

AML12 cells were transfected with pcDNA or pcDNA-MLKL-FLAG, HepG2 cells were transfected with either siControl or siMLKL. Exosome isolation was performed using the Total exosome isolation kit (# 4478359, Invitrogen) as per manufacturer’s instruction and analyzed using Nanosight for particle size and concentration analysis.

### 2.23 Bioinformatics

For untargeted proteomics, heatmaps, and volcano plots, were generated by SRplot (https://www.bioinformatics.com.cn/en), a free online platform for data analysis and visualization. All pathway analyses were performed using ShinyGO 0.80 bioinformatics platform (Ge et al. 2020).

### 2.24 Statistical analyses

All data are represented as mean ±SEM. Two-tailed unpaired t-test or One way ANOVA was used to analyze data with GraphPad Prism. P< 0.05 is considered as statistically significant.

## 3 Results

### 3.1 Generation and characterization of MLKL^HepOE^ mouse model

To generate hepatocyte-specific MLKL-overexpressing mice, 1.5-month-old ROSA26-MLKL mice were injected via tail vein with AAV8-TBG-Cre, which drives liver-specific Cre expression with minimal off-target activity (Yan et al. 2012). Control mice received AAV8-TBG-Null virus (**Figure 1A**). Efficient MLKL overexpression was detected two weeks post-injection (**Figure S1B**). Mice were analyzed 4 months later (∼5.5 months old) to capture effects of sustained, rather than transient, MLKL expression. Body weight was similar between groups (**Figure S1C**), but MLKL^HepOE^ mice showed increased absolute liver weight, while liver-to-body weight and other tissue weight ratios were unchanged (**Figures S1D, S1E**). Western blotting across multiple tissues confirmed that MLKL overexpression was confined to the liver (**Figure 1B**). Hepatic *Mlkl* transcript levels increased 9-fold in MLKL^HepOE^ mice, with MLKL-FLAG expression elevated 3.5-fold and endogenous MLKL levels unchanged (**Figure S1F, Figure 1C**). Hepatocyte and non-parenchymal cell (NPC) isolation confirmed that *Mlkl* overexpression was restricted to hepatocytes (**Figure 1D, Figure S1G**).

**FIGURE 1.**
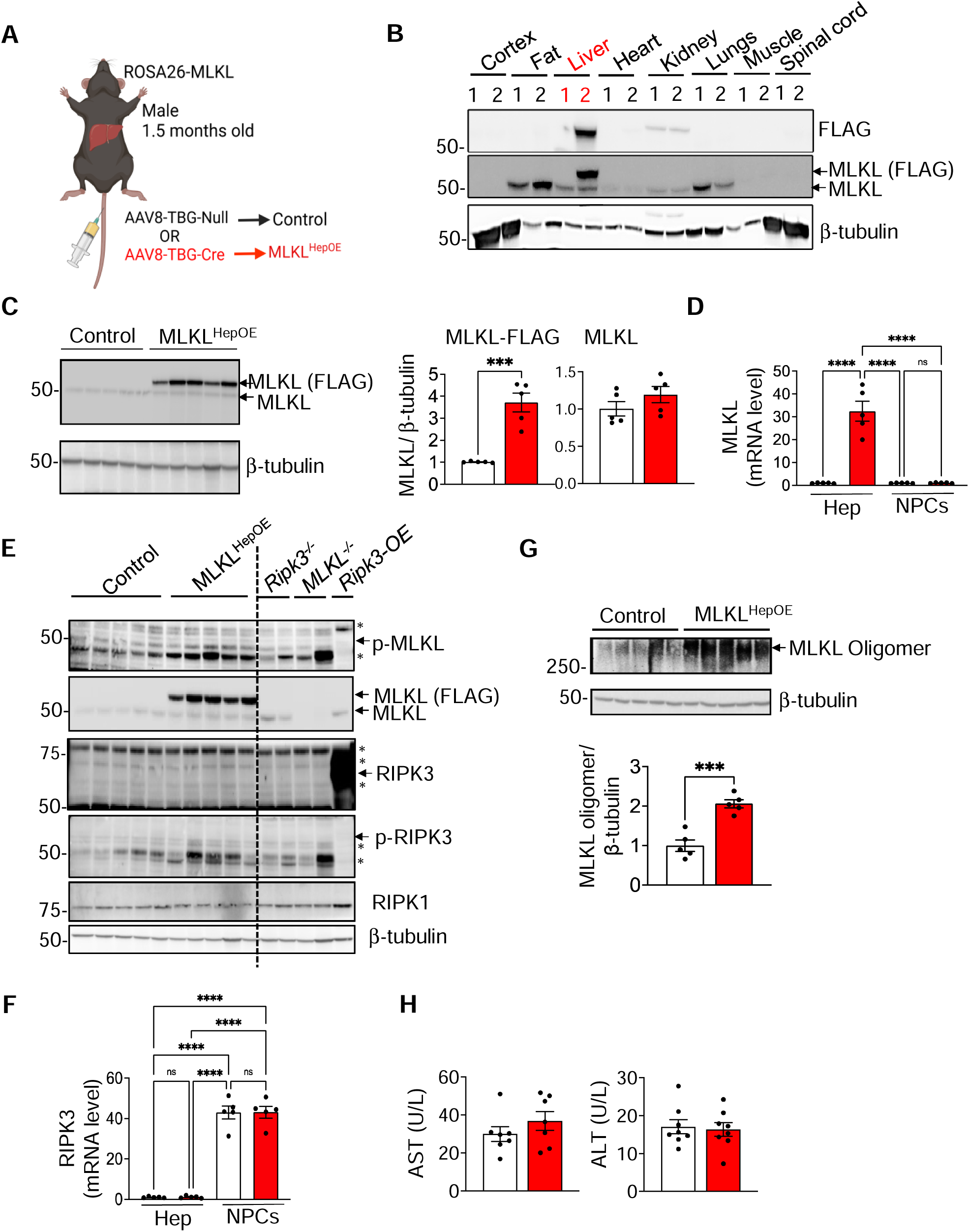
Hepatocyte-specific MLKL overexpression does not induce necroptosis or liver injury. (A) Schematic of the experimental strategy for hepatocyte specific MLKL overexpression. (B) Western blot analysis showing the expression levels of FLAG, MLKL (MLKL-FLAG and endogenous), and β-tubulin (loading control) across various tissues in control (1) and MLKL^HepOE^ (2) mice. (C) *Left:* Western blot analysis of MLKL and β-tubulin (loading control) expression in liver tissue lysates from control and MLKL^HepOE^ mice, 4 months post viral injection. *Right:* Quantification of MLKL-FLAG or endogenous MLKL normalized to β-tubulin and represented as fold change. (D) qPCR analysis of *Mlkl* mRNA levels in isolated hepatocytes (Hep) and liver non-parenchymal cells (NPCs) from control and MLKL^HepOE^ mice. (E) Western blot analysis of liver tissue lysates from control and MLKL^HepOE^ mice for phosphorylated MLKL (p-MLKL), total MLKL, phosphorylated RIPK3 (p-RIPK3), total RIPK3, and RIPK1. β-tubulin was used as a loading control. Liver tissue lysate from *Ripk3*^D*/*D^and *Mlkl*^D*/*D^mice, and cell lysate from HepG2 cells transfected with pcDNA3.1-Ripk3-FLAG served as negative and positive controls. (F) qPCR analysis of *Ripk3* mRNA levels in isolated hepatocytes (Hep) and liver non-parenchymal cells (NPCs) isolated from control and MLKL^HepOE^ mice. (G) *Top*: Western blot analysis showing MLKL oligomers in liver tissue lysates from control and MLKL^HepOE^ mice. *Bottom:* Quantification of MLKL oligomers normalized to β-tubulin. (H) Plasma levels of AST and ALT in control and MLKL^HepOE^ mice. Data represented as mean ± SEM for control (white) and MLKL^HepOE^ (*red*) mice, n=5/group (C-F), n=7-8/group (H). Statistical significance was determined by two-tailed unpaired t-test (C, G, H) or One-way ANOVA (D, F). ****p < 0.0001, ***p < 0.001, ns: non-significant.

### 3.2 Hepatocyte-specific MLKL overexpression does not induce necroptosis, apoptosis, or liver injury

To investigate the effect of hepatocyte MLKL overexpression on necroptosis, we performed western blot analysis of necroptosis pathway proteins in liver tissues from control and MLKL^HepOE^ mice. Phosphorylated MLKL (p-MLKL) was not detected, as similar bands were present in *Mlkl^-/-^*liver tissues (**Figure 1E**). Similarly, phosphorylated RIPK3 (p-RIPK3) showed non-specific bands in both groups, confirmed using *Ripk3^-/-^* liver tissues. RIPK3 protein was undetectable in liver tissue from young mice, validated with RIPK3-FLAG overexpressing HepG2 lysates (**Figure 1E**). Analysis of RIPK3 transcript levels in hepatocyte and non-parenchymal cell (NPC) fractions from control and MLKL^HepOE^ liver tissues showed that RIPK3 expression was negligible in hepatocytes, but detectable in NPCs (**Figure 1F**). RIPK1 levels were similar between groups (**Figure 1E**). Although MLKL overexpression did not alter necroptosis pathway proteins, MLKL oligomer formation increased 2-fold in MLKL^HepOE^ mice (**Figure 1G**). To assess whether MLKL overexpression induces cell death through alternative pathways such as apoptosis, we performed TUNEL staining. The analysis revealed comparable levels of TUNEL-positive cells in liver tissues from both control and MLKL^HepOE^ mice (**Figure S1H**). This was further corroborated by measuring expression of markers of apoptosis, cleaved caspase 3 and cleaved PARP, in the liver tissues of control and MLKL^HepOE^ mice, which showed similar levels of expression (**Figure S1I**). Hepatocyte injury assessment by measuring plasma levels of ALT and AST showed no significant differences between MLKL^HepOE^ and control mice (**Figure 1H**). Thus, hepatocyte-specific MLKL overexpression did not induce necroptosis, apoptosis, or hepatocyte injury, despite increased MLKL oligomer formation, suggesting a non-lethal role for MLKL under basal conditions in young mice.

### 3.3 MLKL overexpression enhances extracellular vesicle (EV) release without inducing cell death

Given that MLKL overexpression did not induce necroptosis despite MLKL oligomer formation, we further investigated its impact in the AML12 mouse hepatocyte cell line, as previous studies suggest a non-necroptotic role for MLKL in EV generation as a potential mechanism to prevent cell death (Yoon et al. 2017). Transfection with pcDNA-MLKL-FLAG resulted in a 10-fold increase in MLKL-FLAG expression compared to control cells (**Figure 2A**). MLKL overexpression did not induce cell death (**Figure 2B**) but enhanced EV release, as measured by nanoparticle tracking analysis (NTA), without altering EV size (**Figure 2C**). Conversely, knocking down MLKL in a human liver cancer cell line HepG2, which exhibits higher MLKL expression compared to the normal human liver cell line THLE2, resulted in a reduction in both the concentration and size of EVs, as determined by NTA (**Figure S2A-C**). Western blotting showed increased EV markers (ALIX, VPS4B, TSG101), exogenous MLKL-FLAG, and the DAMP, HMGB1 in EVs from MLKL-overexpressing AML12 cells compared to controls (**Figure 2D**). Consistent with this, plasma samples from MLKL^HepOE^ mice contained higher levels of circulating MLKL compared to control mice (**Figure 2E**). Importantly, we observed that MLKL levels in the plasma of MASH patients are significantly higher than in healthy individuals (**Figure 2F**). Thus, MLKL-driven EV release could serve as a non-necroptotic, cell-protective mechanism that limits hepatocyte death in the context of MLKL overexpression and oligomerization.

**FIGURE 2.**
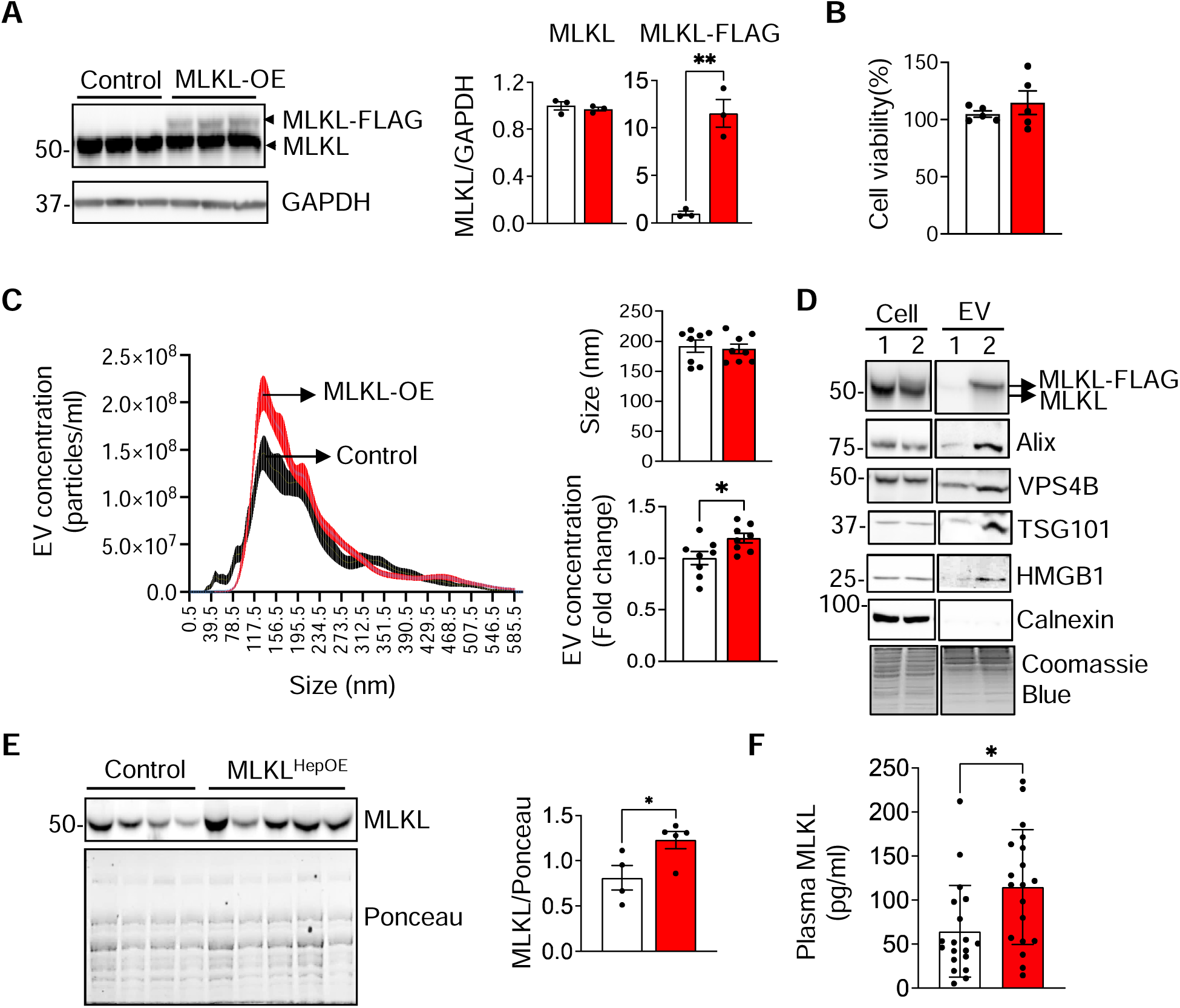
Hepatocyte MLKL overexpression promotes extracellular vesicle release and increases circulating MLKL levels. AML12 mouse hepatocytes were transfected with either empty vector (pcDNA, control) or pcDNA-MLKL-FLAG (MLKL-OE) for 24 hours and the following assays were performed. All cell culture experiments were conducted in triplicate. (A) *Left:* Western blot analysis showing expression of MLKL (MLKL-FLAG and endogenous) and GAPDH (loading control). *Right*: Quantification of MLKL expression normalized to GAPDH and expressed as fold change. (B) Cell viability assessed by CCK-8 assay. (C) *Left*: Nanoparticle Tracking Analysis (NTA) of EVs isolated from control or MLKL overexpressing AML12 cells. *Right:* Quantification of EV size and concentration (*bottom*). (D) Western blot analysis of cell lysates and EVs from control (1) and MLKL-overexpressing (2) AML12 cells for markers of EVs (ALIX, VPS4B, TSG101), HMGB1, MLKL, and calnexin (marker for endoplasmic reticulum). Coomassie blue stained gel is used as loading control. (E) Western blot analysis (*left*) and quantification (*right,* represented as fold change) of MLKL in plasma from control and MLKL^HepOE^ mice at 4 months post overexpression (n=4-5/group). Ponceau stained membrane is used as loading control. (F) MLKL levels in the plasma of normal and MASH patients (n=20/group), as determined by ELISA. Data are presented as mean ± SEM. Statistical significance was determined by two-tailed unpaired t-test. **p<0.01, *p<0.05.

### 3.4 Impact of hepatocyte MLKL overexpression on liver proteome

To understand how MLKL overexpression affects liver physiology, we performed untargeted label-free quantitative proteomics of liver tissues from control and MLKL^HepOE^ mice. Principal component analysis showed clear separation along PC1 (37.8%), with tightly clustered controls and greater variability among MLKL^HepOE^ samples (**Figure 3A**). A heatmap of z-score normalized protein expression revealed distinct proteomic signatures between groups (**Figure 3B**). Differential expression analysis, visualized in a volcano plot (**Figure 3C**), identified upregulated proteins (e.g., CYP4A1, LASP1, SLC16A1, COBLL1, ACOX1, COQ9, MYO1B) linked to fatty acid metabolism, mitochondrial function, and oxidative stress, and downregulated proteins (e.g., SAA2, ABCB11, SEC22B, ENO3, PKM, SEPTIN11) associated with lipid transport, glycolysis, and cytoskeletal organization (**Table S3**). These shifts suggest that MLKL overexpression alters pathways involved in lipid handling, mitochondrial activity, and cellular metabolism.

**FIGURE 3.**
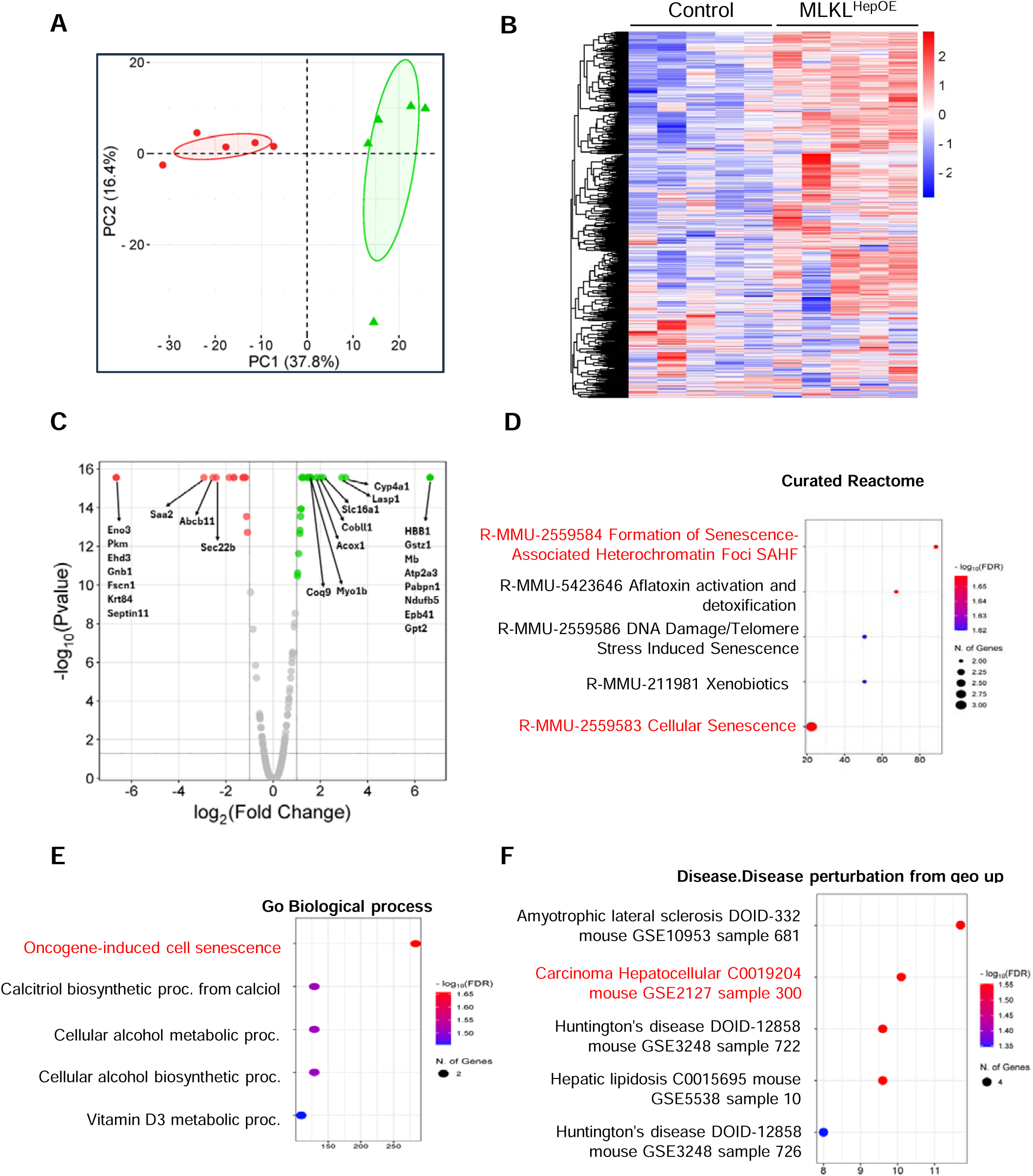
Untargeted label-free quantitative proteomics of liver from control and MLKL^HepOE^ mice. (A) Principal Component Analysis (PCA) plot showing distinct clustering of control (red circles) and MLKL^HepOE^ (green triangles) liver proteomes (n=5/group). (B) Heatmap representing hierarchical clustering of total identified and quantified proteins between control and MLKL^HepOE^ groups, with Z-score normalized expression values. (C) Volcano plot displaying significantly upregulated (green) and downregulated (red) proteins in MLKL^HepOE^ vs control livers (1.5 fold up/down, p-value < 0.05)). Selected significantly altered proteins are labeled. (D) Reactome Pathway enrichment analysis of up-regulated proteins in MLKL^HepOE^ vs control livers. The dot plot shows the top enriched Reactome pathways. (E) Gene Ontology (GO) enrichment analysis for MLKL^HepOE^ vs control livers. The dot plot shows enriched GO biological processes among differentially expressed proteins. (F) Disease association analysis. The dot plot displays disease terms significantly associated with the differentially expressed proteins. In 3D-F, the size of each dot indicates the number of genes associated with the pathways, while dot color represents statistical significance, with red indicating higher and blue indicating lower significance.

To gain functional insights into proteomic alterations in MLKL^HepOE^ livers, we performed reactome pathway enrichment analysis (**Figure 3D**). MLKL^HepOE^ vs. control mice showed significant enrichment of cellular senescence and senescence-associated heterochromatin foci (SAHF) pathways, with cellular senescence having the highest number of associated proteins. Gene Ontology (GO) biological process analysis also identified significant enrichment of pathways, notably oncogene-induced cell senescence (**Figure 3E**), suggesting that MLKL overexpression drives senescence-related mechanisms. Disease-perturbed pathway analysis revealed enrichment of pathways associated with HCC and hepatic lipidosis in MLKL^HepOE^ mice (**Figure 3F**), indicating that hepatocyte MLKL overexpression may influence both cancer-related and metabolic dysfunction pathways in the liver.

### 3.5 Hepatocyte MLKL overexpression upregulated markers of cellular senescence, inflammatory macrophage recruitment, and promotes lipid accumulation with age

To assess the impact of hepatocyte-specific MLKL overexpression on liver senescence, we performed SPiDER-β-gal staining, a widely used method for detecting senescence-associated β-galactosidase (SA-β-Gal) activity, in control and MLKL^HepOE^ mice (Jannone et al. 2020). SA-β-Gal activity was significantly increased in MLKL^HepOE^livers (**Figure 4A**), indicating enhanced senescence. Consistently, cell cycle arrest markers *p16* (3-fold), *p15* (2.7-fold), and *p21* (10-fold) were upregulated, while *p53* mRNA remained unchanged but p53 protein levels increased 2.5-fold (**Figures 4B, 4C**). SASP cytokines and chemokines, including *TNF*α (3.6-fold), *IL1*β (1.7-fold), *IFN*γ (1.5-fold), *CXCL1* (5.5-fold), *CXCL2* (2.3-fold), and *CXCL10* (1.7-fold), were elevated (**Figure 4D**), whereas SASP growth factors and proteases were unchanged (**Figure S3A**). Since the SASP can recruit macrophages and promote a pro-inflammatory M1 phenotype (Behmoaras and Gil 2021), we observed a 2-fold increase in F4/80-positive macrophages (**Figure 4E**) and upregulation of M1 markers (*CD68*: 1.8-fold, *CD11c*: 2-fold, *TLR4*: 1.5-fold) (**Figure 4F**). M2 markers remained unchanged (*Fizz1*, *CD206*) or showed modest increases (*Arg1*: 1.2-fold, *CD163*: 2.2-fold) (**Figure S3B**).

**FIGURE 4.**
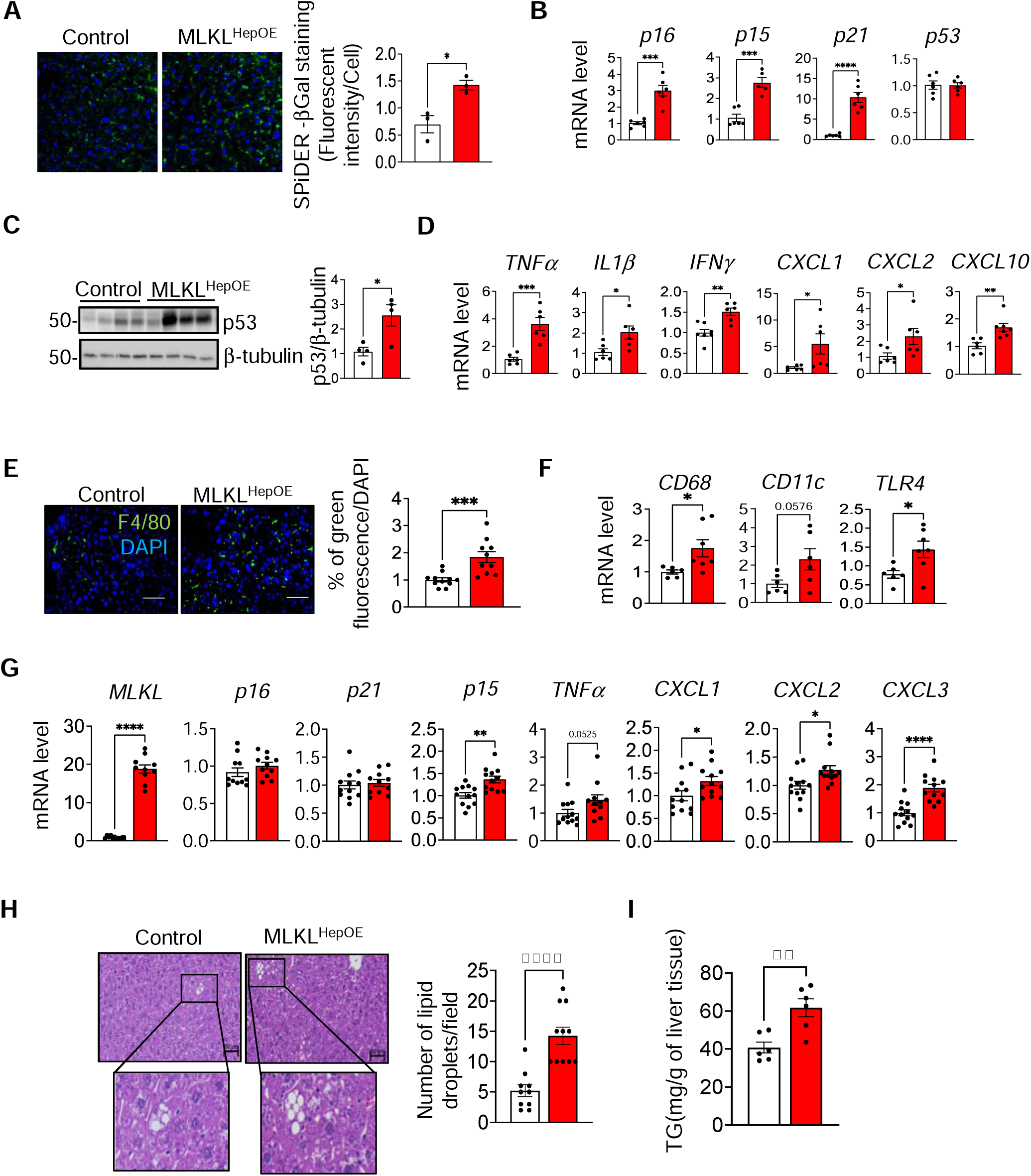
Hepatocyte MLKL overexpression up regulates markers of cellular senescence, inflammatory macrophage recruitment, and promotes lipid accumulation with age. (A) *Left*: Representative images showing SPiDER-βGal staining in liver sections from control and MLKL^HepOE^ mice. SPiDER-βGal–positive cells are shown in green, nuclei counterstained with DAPI (blue). Magnification: 630. *Right:* Quantification of SPiDER-βGal staining represented as green fluorescence intensity per cell from n = 3 mice per group. (B) qPCR analysis for cell cycle arrest markers p16, p15, p21, and p53 in the livers of control and MLKL^HepOE^ mice. (C) *Left:* Western blot analysis showing p53 and β-tubulin (loading control) in control and MLKL^HepOE^ mice livers. *Right:* Quantification of p53 normalized to β-tubulin. (D) qPCR analysis for SASP cytokines and chemokines in the livers of control and MLKL^HepOE^ mice. (E) *Left:* Representative images showing immunofluorescence staining of liver sections from control and MLKL^HepOE^ mice for F4/80 and DAPI (blue). *Right:* Percentage of green fluorescence normalized to DAPI. Scale bar= 50µm, Magnification: 200X. (F) qPCR analysis for M1 macrophage markers in the livers of control and MLKL^HepOE^ mice livers. (G) qPCR analysis for MLKL, cell cycle arrest markers (p16, p15, p21) and SASP (TNFα, CXCL1, CXCL2, CXCL3) in AML12 cells transfected with either pcDNA (control) or pcDNA-MLKL-FLAG for 24 hours. (H) *Left:* H&E staining of liver sections from control and MLKL^HepOE^ mice, 13 months post viral injection. Enlarged insets highlight lipid droplets. *Right*: Quantification of the number of lipid droplets/field. (I) Liver triglyceride levels from control and MLKL^HepOE^ mice, 13 months post viral injection. For all animal studies, n=4-6 mice/group were used and data are represented as fold change. Control (white) and MLKL^HepOE^ (red). Data are presented as mean ± SEM. Statistical significance was determined by two-tailed unpaired t-test. ****p < 0.0001, ***p < 0.001, **p<0.01, *p<0.05.

To define whether MLKL overexpression induces senescence in hepatocytes in a cell-autonomous manner, we transiently overexpressed MLKL in AML12 cells and measured senescence markers 24 hours post-transfection. MLKL overexpression significantly increased p15 expression, while p16 and p21 levels remained unchanged (**Figure 4G**). Among SASP factors, CXCL1, CXCL2, and CXCL3 were significantly upregulated, with TNFα showing a trend toward increase (p = 0.0525). No significant changes were observed in SASP growth factors or proteases (**Figure S3F**). These results suggest that MLKL overexpression induces a partial senescence-like response, marked by selective upregulation of p15 and specific inflammatory mediators in AML12 cells.

Cellular senescence in hepatocytes alters glucose and lipid metabolism, promoting steatosis and triglyceride production, which contribute to MASH pathology (Bonnet et al. 2022). Histological analysis and triglyceride measurements at 5.5 months did not reveal significant differences in lipid accumulation or fibrosis markers between MLKL^HepOE^and control mice (**Figures S3C-E**), indicating that short-term MLKL overexpression does not cause steatosis, fibrosis, or liver injury. However, increased lipid droplets and triglyceride levels were observed at 13 months (**Figures 4H, 4I**). These findings suggest that MLKL overexpression promotes fat accumulation over time, potentially contributing to MASH.

### 3.6 MLKL overexpression induces mitochondrial dysfunction and alters lipid metabolism in the liver

Dysregulated autophagy, NF-κB activation, and mitochondrial dysfunction are known inducers of senescence during aging (Wiley et al. 2016; Li et al. 2023; Salminen et al. 2012). To assess whether these pathways contribute to MLKL-induced senescence, we first examined autophagy and NF-κB activation markers in MLKL^HepOE^ livers. Both pathways were comparable between MLKL^HepOE^ and controls, suggesting they do not drive senescence (**Figure S4A, B**). In contrast, oxidative stress markers 4-Hydroxynonenal (4-HNE) and Nitrotyrosine (NT) were significantly elevated in MLKL^HepOE^ livers (**Figure 5A, 5B**). Given the link between mitochondrial dysfunction, oxidative stress, and senescence (Wiley et al. 2016; Cui et al. 2012), we performed targeted proteomics of mitochondrial proteins to define a potential involvement of mitochondrial dysfunction and oxidative stress in the induction of senescence in MLKL^HepOE^ mouse liver. Six of 25 Krebs cycle proteins and 10 of 31 β-oxidation proteins were upregulated, indicating altered mitochondrial metabolism (**Figure 5C**), while other mitochondrial pathways remained unchanged (**Figure S4C, Table S4**). Lipidomics analysis revealed significant changes, with increased monoacylglycerophosphoglycerol (MGMG), diacylglycerol (DG), and triacylglycerol (TG), and decreased lysodimethylphosphatidylethanolamine (LdMePE) and sphingomyelin (SM) in MLKL^HepOE^ livers (**Figure 5D, Table S5**), suggesting disrupted lipid metabolism. To directly assess mitochondrial function, a mitostress test in MLKL-overexpressing AML12 cells showed significant reductions in basal, maximal, and ATP-coupled respiration, spare respiratory capacity, proton leak, and non-mitochondrial respiration (**Figure 5E**). Together, these findings suggest that MLKL overexpression promotes mitochondrial dysfunction, oxidative stress, and metabolic dysfunction.

**FIGURE 5.**
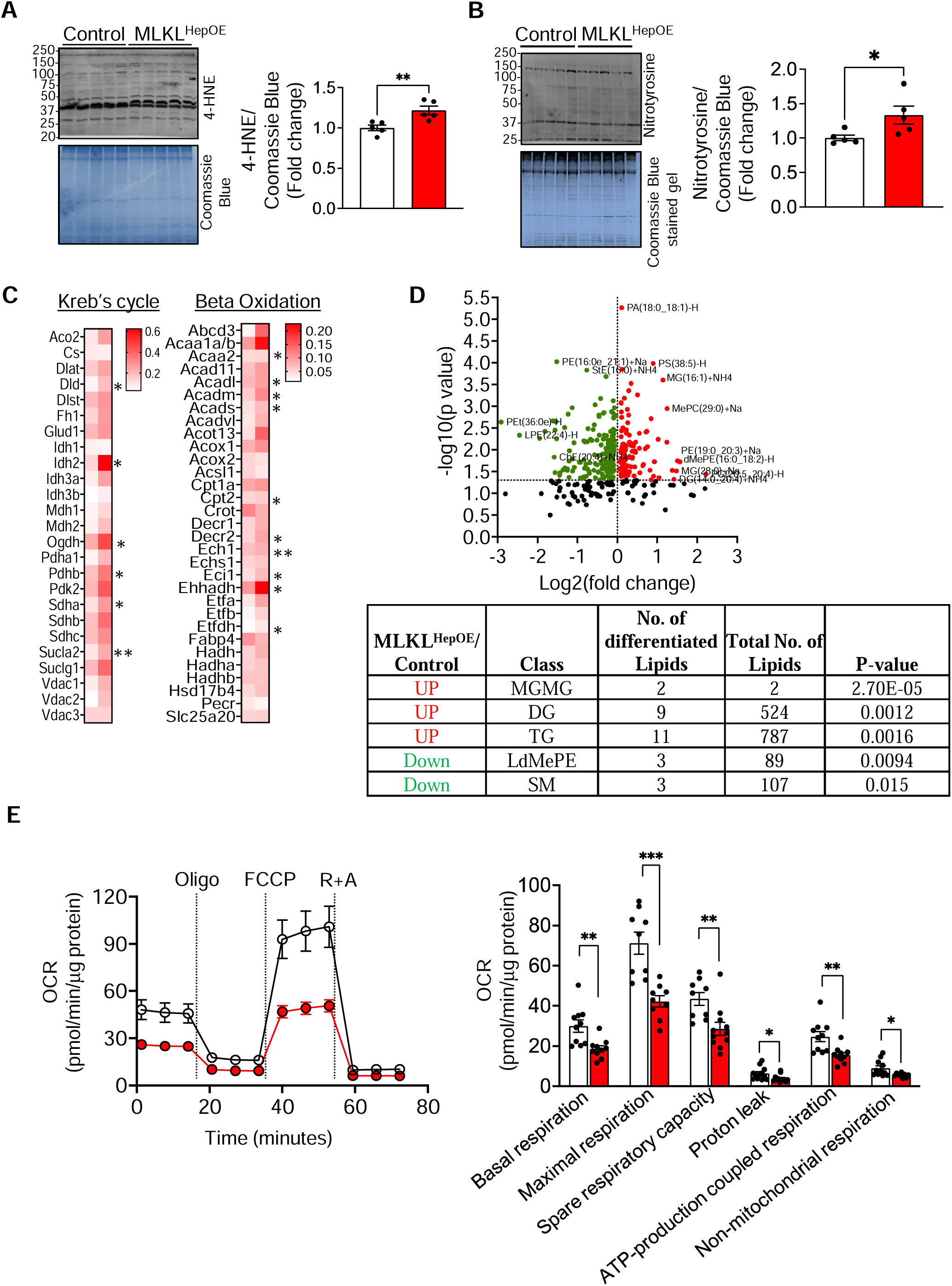
MLKL overexpression induces mitochondrial dysfunction and alters lipid metabolism in the liver. Western blots *(left)* and corresponding densitometric quantification (*right)* of 4-HNE (A) and Nitrotyrosine (B) protein adducts in control and MLKL^HepOE^ mice livers. Coomassie Blue stained gel used as loading control. (C) Heat maps of proteins involved in Kreb’s and TCA cycle obtained by targeted mitochondrial proteomics of liver tissues from MLKL^HepOE^ vs control mice. Red represents up regulation and white denotes no significant change. Asterisks (*) denote statistically significant changes between the two groups. (D) Volcano plot displaying lipidomic changes in MLKL^HepOE^ vs control mice. Each dot represents a lipid species. Red dots and green dots indicate significantly upregulated and down regulated lipids, respectively. Black dots indicate non-significant changes. The table summarizes the total number of differentially accumulated lipid classes. (E) Mitochondrial respiration analysis in control and MLKL overexpressing AML12 mouse hepatocytes. Left: a representative oxygen consumption rate (OCR) curve, with sequential addition of oligomycin, FCCP, and rotenone/antimycin A. Right: bar graph summarizing the quantified respiratory parameters. Control (white) and MLKL^HepOE^ (red). Data are presented as mean ± SEM. Statistical significance was determined by two-tailed unpaired t-test. ***p < 0.001, **p<0.01, *p<0.05.

### 3.7 MLKL overexpression alters mitochondrial dynamics in the liver

Transmission electron microscopy of liver tissues revealed altered mitochondrial morphology in MLKL^HepOE^ mice compared to control mice (**Figure 6A**). Although mitochondrial fragmentation or swelling was not observed, a subset showed abnormal C-shaped morphology without changes in mitochondrial area (**Figure S5A**), suggesting altered mitochondrial dynamics. To further assess this, we examined key fusion proteins (mitofusin 1, Mfn1; Mfn2; optic atrophy 1, OPA1) and fission proteins [Fis1, Drp1, and phospho-Drp1 at Ser637 (inhibitory) and Ser616 (pro-fission)] (Knott et al. 2008). MLKL overexpression significantly increased the ratios of phospho-Drp1 (Ser637 and Ser616) to total Drp1 (**Figures 6B, 6C**), while Mfn1, Mfn2, and Fis1 levels were unchanged. Notably, OPA1 levels were markedly reduced in MLKL^HepOE^ livers. VDAC levels remained similar between groups, indicating mitochondrial mass was unaffected. Together, these results suggest that MLKL overexpression disrupts mitochondrial dynamics.

**FIGURE 6.**
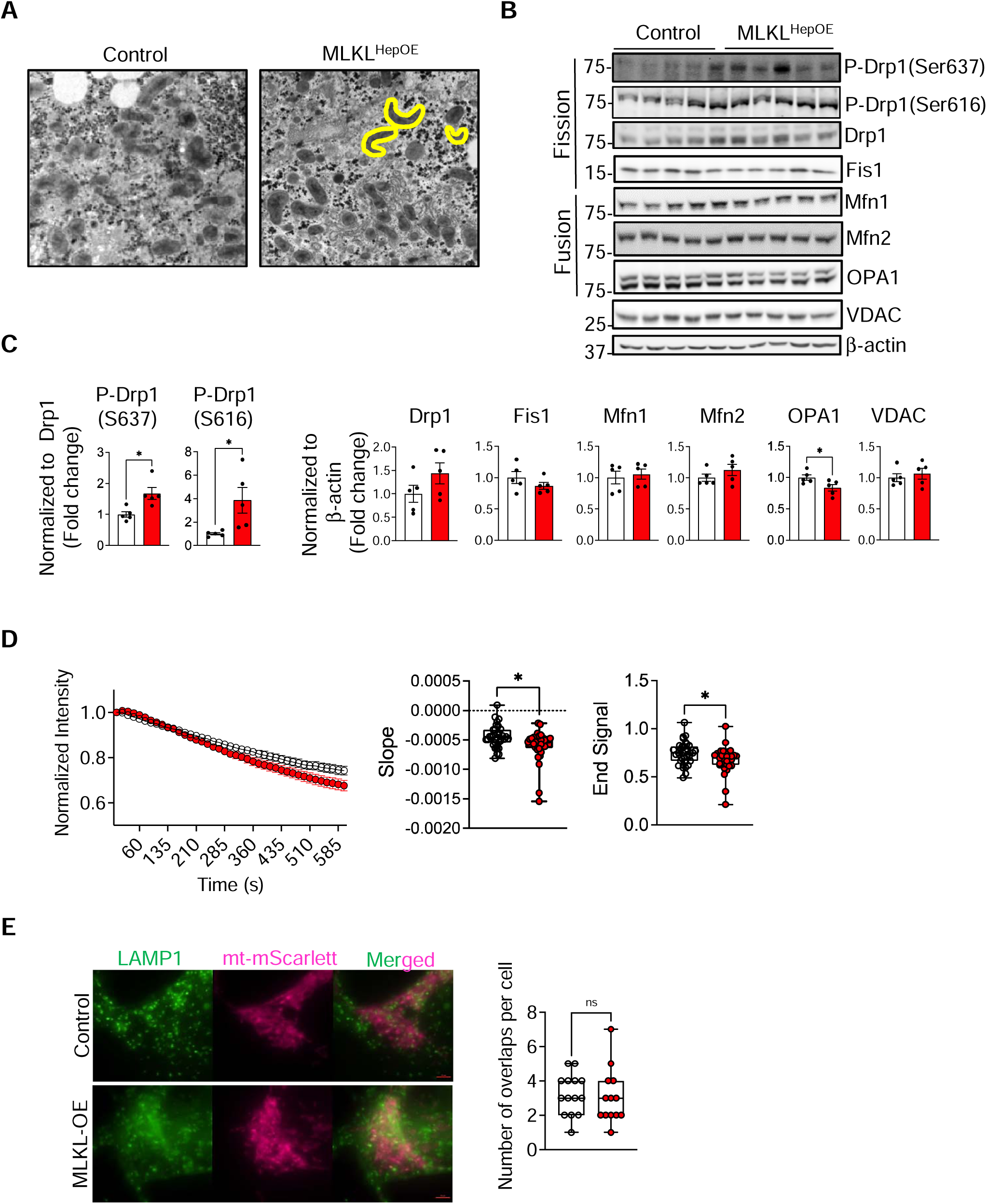
Hepatic MLKL overexpression altered mitochondrial morphology and dynamics. (A) Transmission electron microscopy images of liver tissue from control and MLKL^HepOE^ mice. Magnification: 1000X. (B) Western blot analysis of mitochondrial fusion and fission proteins, VDAC and β-actin (loading control) in control and MLKL^HepOE^ mice livers. (C) Quantification of fission and fusion protein levels normalized to β-actin and represented as fold change. (D) *Left:* Photoactivatable GFP (paGFP) signal intensity over 24 hours in control and MLKL overexpressing AML12 cells. *Right:* Quantification of the paGFP signal, showing slope and end signal. (E) *Left*: Representative images showing colocalization of the lysosomal marker LAMP1 with the mitochondrial marker mt-mScarlett (magenta) in control and MLKL overexpressing AML12 cells. *Right:* Quantification of the number of LAMP1-mt-mScarlett overlaps. Control (white) and MLKL^HepOE^ (red). Data are presented as mean ± SEM. Statistical significance was determined by two-tailed unpaired t-test.*p<0.05, ns: not significant.

To directly test MLKL’s role in mitochondrial dynamics, we measured the spread of matrix-targeted photoactivatable GFP (paGFP) signal 24 hours after transfection in control and MLKL-overexpressing AML12 cells. MLKL-overexpressing cells exhibited a steeper decline in paGFP signal, suggesting enhanced mitochondrial trafficking or dynamics (**Figure 6D**). To assess whether altered dynamics impacted mitophagy, we measured colocalization of the endolysosomal marker LAMP1 with the mitochondrial marker mt-mScarlett. Representative images showed regions of overlap (yellow) in both control and MLKL-overexpressing cells (**Figure 6E**, *left*). Quantification revealed no significant difference in LAMP1/mt-mScarlett interactions per cell (**Figure 6E**, *right*), indicating no change in mitophagy. Consistently, Parkin and PINK1 expression levels were similar in control and MLKL^HepOE^ livers (**Figure S5B**). These findings suggest that MLKL alters mitochondrial morphology independently of mitophagy.

## 4 Discussion

Aging is associated with chronic low-grade inflammation and metabolic dysfunction, both contributing to the progression of liver diseases such as MASLD and HCC. While MLKL is best known as the terminal effector of RIPK3-dependent necroptosis, emerging evidence highlights RIPK3-independent roles for MLKL in endosomal trafficking, EV generation (Yoon et al. 2017), and autophagy (Wu et al. 2020). Here, we show that hepatocyte-specific MLKL overexpression, independent of necroptotic death, induces mitochondrial dysfunction, oxidative stress, and senescence, key processes implicated in aging and disease progression in the liver. We propose that MLKL overexpression disrupts mitochondrial dynamics, impairing mitochondrial function and elevating oxidative stress, which in turn triggers cell cycle arrest, the SASP, macrophage recruitment, and hepatic steatosis.

Upregulation of the necroptosis pathway has been reported in various liver disease conditions and in liver aging (Afonso et al. 2015; Gautheron et al. 2015; Mohammed, Thadathil, et al. 2021). However, in our study, hepatic overexpression of MLKL in young (5.5-month-old) mice did not lead to activation of the canonical necroptosis pathway. This is supported by the lack of detectable RIPK3 expression, absence of its phosphorylated form (p-RIPK3) and p-MLKL, as well as unchanged hepatocyte injury markers such as ALT and AST. Although the ability of hepatocytes to undergo necroptosis remains debated (Dara et al. 2016), recent studies suggest that RIPK3 is epigenetically silenced in hepatocytes, making them resistant to necroptosis (Preston et al. 2022; Dara 2022). Interestingly, despite the absence of RIPK3, which is required for MLKL phosphorylation and oligomerization, our study found a significant increase in MLKL oligomerization in the livers of MLKL^HepOE^ mice. While the mechanism underlying this RIPK3-independent oligomerization is unclear, previous studies suggest several possibilities: TAM kinases (Tyro3, Axl, and Mer) can directly phosphorylate MLKL at Tyr376, promoting oligomerization independently of RIPK3 (Najafov et al. 2019). Thioredoxin-1 (Trx1) deficiency has also been shown to enhance MLKL oligomerization regardless of RIPK3 activity (Reynoso et al. 2017). Moreover, expression of MLKL’s N-terminal domain (residues 1–125 or 1–180) is sufficient to induce oligomerization in the absence of RIPK3 (Hildebrand et al. 2014). The independence of oligomerization from RIPK3 in our study highlights the existence of alternative regulatory mechanisms, such as phosphorylation by TAM kinases that could specifically prime MLKL for these non-necroptotic functions. Importantly, while oligomerization is a prerequisite for necroptosis, it is not sufficient to induce cell death. Additional steps, such as membrane translocation and pore formation, are required. MLKL oligomers can also be ubiquitylated to suppress necroptosis (Liu et al. 2021), and membrane localization is essential for execution of cell death (Samson et al. 2020). Therefore, the observed MLKL oligomerization, although not leading to cell death in this context, could still be an essential conformational switch required for MLKL to engage in its non-canonical roles in mitochondrial regulation and EV biogenesis. However, this possibility remains to be tested in future studies. Thus, our findings suggest that hepatic MLKL overexpression promotes its oligomerization through RIPK3-independent mechanisms without triggering necroptosis, potentially influencing alternative cellular pathways such as EV release to preserve hepatocyte viability.

While MLKL is classically known to promote mitochondrial fragmentation during necroptosis via PGAM5-mediated Drp1 activation (Dan et al. 2023), our results reveal a distinct mechanism. MLKL overexpression increased phosphorylation of Drp1 at both Ser616 (pro-fission) and Ser637 (inhibitory), indicating complex fission regulation in hepatocytes. MLKL also reduced the expression of the inner membrane fusion protein OPA1, without affecting Mfn1 or Mfn2, suggesting a selective shift in mitochondrial dynamics or cristae reorganization (Frezza et al. 2006; Lee et al. 2017; Suga et al. 2023). Despite this imbalance, mitophagy was not activated, as PINK1 and Parkin levels and mitochondria–lysosome colocalization remained unchanged. MLKL-overexpressing livers exhibited C-shaped mitochondria, a morphology linked to excessive fission (Mahajan et al. 2021), along with impaired mitochondrial respiration and elevated oxidative stress (Zong et al. 2024; Srinivasan et al. 2017). Prior studies showed MLKL localizes to mitochondria (Wang et al. 2025) and that its deletion in hepatocytes improves respiration and β-oxidation (Majdi et al. 2020) while reducing oxidative DNA damage (Xu et al. 2023). Together, our data reveal a noncanonical role for MLKL in regulating mitochondrial dynamics and function, independent of necroptosis, with important implications for liver metabolic health.

Mitochondrial dysfunction is a known driver of cellular senescence through mechanisms such as ROS-induced DNA damage (Passos et al. 2010; Correia-Melo et al. 2016), mitochondrial dysfunction-associated senescence (MiDAS) (Wiley et al. 2016), AMPK-mediated energy stress (Correia-Melo and Passos 2015), and mtDNA-driven cGAS–STING activation (Victorelli et al. 2023). In our study, hepatocyte-specific MLKL overexpression impaired mitochondrial function, increased oxidative stress, and upregulated senescence markers (p15, p16, p21, p53) along with pro-inflammatory SASP cytokines and chemokines, but not growth factors or proteases. Similar trends were observed in AML12 cells, though only p15 and inflammatory SASP components were elevated. These findings suggest that independent of RIPK3 and necroptosis, MLKL promotes hepatocyte senescence, likely via ROS-induced DNA damage, supporting a noncanonical role for MLKL in inflammaging and liver disease progression. Consistently, our prior work showed that MLKL deficiency reduces p16, p21, and SASP expression in aged *Mlkl^-/-^*livers (Mohammed et al. 2025). Hepatocyte senescence has been implicated in age-related hepatic steatosis via mitochondrial dysfunction and lipid dysregulation (Ogrodnik et al. 2017). In line with this, short-term MLKL overexpression altered hepatic lipid metabolites, while long-term overexpression exacerbated steatosis, suggesting a link between MLKL-driven senescence and lipid accumulation. SASP factors, particularly chemokines, recruit immune cells and perpetuate chronic inflammation (Coppe et al. 2008). Accordingly, MLKL^HepOE^ livers exhibited increased F4/80 macrophage infiltration, upregulation of M1 markers, and elevated EV release containing HMGB1, a known macrophage activator (Lee et al. 2014). Given that MLKL regulates EV biogenesis independently of RIPK3 (Yoon et al. 2017), these findings further support a non-necroptotic, pro-inflammatory role for MLKL via intercellular communication.

Our comparative proteomic analysis revealed enrichment of pathways associated not only with HCC and hepatic lipidosis, but also with neurodegenerative diseases such as Amyotrophic Lateral Sclerosis (ALS) and Huntington’s disease. Given that inflammation is a key driver of HCC (Yang et al. 2019), the enrichment of HCC-associated pathways supports a role for hepatocyte MLKL in tumorigenesis. The identification of lipidosis-related pathways aligns with the observed steatosis and altered lipid metabolism, suggesting that MLKL drives metabolic dysfunction independent of cell death. Notably, the enrichment of ALS- and Huntington’s disease-related pathways raises the possibility that hepatic MLKL overexpression exerts systemic effects, potentially via the release of inflammatory mediators or EVs influencing distant organs such as the brain. This aligns with emerging evidence linking liver pathology to neuroinflammation, cognitive decline, and Alzheimer’s disease (Yang et al. 2024; Kjaergaard et al. 2024; Garcia-Martinez and Cordoba 2012; Estrada et al. 2019), supporting a role for liver-brain crosstalk in aging and disease. Together, these findings suggest that MLKL-mediated signaling in hepatocytes intersects with oncogenic, metabolic, and neurodegenerative pathways, highlighting the broad pathological impact of MLKL dysregulation.

In summary, we show that MLKL induction, in the absence of canonical necroptotic signaling, leads to significant alterations in mitochondrial dynamics, cellular senescence, and intercellular communication, revealing a novel, non-lethal role for this protein in liver aging and disease. The observation of MLKL oligomerization without accompanying hepatocyte injury (ALT, AST) or necroptotic signaling (p-RIPK3, p-MLKL) further supports distinct functional outcomes of MLKL activation. By enhancing the release of pro-inflammatory EVs, MLKL may drive a feed-forward loop of hepatic inflammation. Together, these findings position MLKL as a regulator of mitochondrial and inflammatory homeostasis, distinct from its classical necroptotic function. A critical next step is to determine whether MLKL exerts similar effects in other liver cell types, such as macrophages, hepatic stellate cells, and endothelial cells, or in extrahepatic tissues involved in metabolic regulation. This will help define the broader physiological relevance of MLKL’s noncanonical functions and identify potential cell type-specific contributions to aging.

### Limitations of the study

While hepatocyte-specific MLKL overexpression induces senescence-associated markers in the liver, the precise liver cell type(s) undergoing senescence remains unclear. Although MLKL overexpression in AML12 cells upregulated p15 and some SASP components, it did not fully recapitulate the *in vivo* phenotype, suggesting that hepatocytes may not be the sole or primary senescent population. It is also possible that non-parenchymal cells undergo secondary senescence in response to MLKL-mediated signals, such as extracellular vesicles or inflammatory cues. Future studies will address this by using *in situ* co-localization of senescence and cell type-specific markers, or single-cell transcriptomics, to identify and characterize senescent populations. Additionally, although MLKL oligomerization was observed in the absence of RIPK3, the mechanism remains undefined. Potential contributors such as TAM kinases or Trx1 deficiency were not directly tested. Further biochemical studies are needed to elucidate how MLKL oligomerizes independently of canonical necroptotic signaling.

## Author Contributions

S.M., C.J., T.P., S.B., and P.O.-M. performed experiments, analyzed data, prepared figures, and contributed to manuscript writing and editing. C.G. and J.D.W. analyzed lipidomics data. K.P. performed ELISA for MLKL in human samples provided by C.H.; Z.P., A.S., Z.Y., and N.A. performed and analyzed label-free quantitative proteomics of liver samples. A.T. assisted with animal studies. M.K. conducted and analyzed targeted mitochondrial proteomics. T.L. provided training with AAV8 viral injections. B.H. provided input on EV isolation and characterization protocols. V.G. provided suggestions for senescence marker analysis. T.L.L. designed mitochondrial reporter assays. S.S.D. designed the experiments, supervised the research, wrote the manuscript. All authors reviewed and edited the manuscript.

## Acknowledgements

This work was supported by NIH grants (R01AG059718 and R03CA262044 to S.S.D), the Harold Hamm Diabetes Center–Stephenson Cancer Center Team Science Grant, and the Oklahoma Center for Adult Stem Cell Research (OCASCR) grant to S.S.D; NIH MIRA grant (R35GM137921) to T.L.L.; and RF1AG068283 to V.G. Salary support for S.M. was provided by Oklahoma Center for the Advancement of Science and Technology (OCAST) Health Research Postdoctoral Fellowship. N.A. gratefully acknowledges initial funding from the Office of the Vice President for Research and Partnerships (VPRP) at the University of Oklahoma for establishing the Proteomics Core Facility. The authors thank the Stephenson Cancer Center Tissue Pathology Core for histology services, the Stephenson Molecular Imaging core for the NanoSight NS300 NTA system and the OMRF Imaging core facility for the electron microscopy services. The Agilent Seahorse XF analyzer system was provided by the MTCRO-COBRE Cell and Molecular Imaging core and Cancer Functional Genomics core which are supported by the grants P30GM154635 (National Institute of General Medical Sciences) and P30CA225520 (National Cancer Institute).

## Data availability statement

Proteomics data can be found in the MassIVE database via MSV000097694. Additional data supporting the study’s findings are provided within the manuscript and its supplementary materials. Correspondence and requests for information should be addressed to S.S.D.

## References

Afonso, M. B., P. M. Rodrigues, T. Carvalho, et al. 2015. “Necroptosis is a key pathogenic event in human and experimental murine models of non-alcoholic steatohepatitis”. Clinical science (London, England) 129, 721–39.10.1042/CS20140732.

Ahsan, N., L. Fornelli, F. Z. Najar, et al. 2023. “Proteomics evaluation of five economical commercial abundant protein depletion kits for enrichment of diseases-specific biomarkers from blood serum”. Proteomics 23, e2300150.10.1002/pmic.202300150.

Behmoaras, J., and J. Gil. 2021. “Similarities and interplay between senescent cells and macrophages”. The Journal of Cell Biology 220.10.1083/jcb.202010162.

Bhaskaran, S., G. Pharaoh, R. Ranjit, et al. 2018. “Loss of mitochondrial protease ClpP protects mice from diet-induced obesity and insulin resistance”. EMBO Reports 19.10.15252/embr.201745009.

Blouin, A., R. P. Bolender, and E. R. Weibel. 1977. “Distribution of organelles and membranes between hepatocytes and nonhepatocytes in the rat liver parenchyma. A stereological study”. The Journal of Cell Biology 72, 441–55.10.1083/jcb.72.2.441.

Bonnet, L., I. Alexandersson, R. K. Baboota, et al. 2022. “Cellular senescence in hepatocytes contributes to metabolic disturbances in NASH”. Frontiers in Endocrinology (Lausanne) 13, 957616.10.3389/fendo.2022.957616.

Coppe, J. P., C. K. Patil, F. Rodier, et al. 2008. “Senescence-associated secretory phenotypes reveal cell-nonautonomous functions of oncogenic RAS and the p53 tumor suppressor”. PLoS Biology 6, 2853–68.10.1371/journal.pbio.0060301.

Correia-Melo, C., F. D. Marques, R. Anderson, et al. 2016. “Mitochondria are required for pro-ageing features of the senescent phenotype”. The EMBO Journal 35, 724–42.10.15252/embj.201592862.

Correia-Melo, C., and J. F. Passos. 2015. “Mitochondria: Are they causal players in cellular senescence?”. Biochimica et Biophysica Acta 1847, 1373–9.10.1016/j.bbabio.2015.05.017.

Cui, H., Y. Kong, and H. Zhang. 2012. “Oxidative stress, mitochondrial dysfunction, and aging”. Journal of Signal Transduction 2012, 646354.10.1155/2012/646354.

Dan, Sang, X. Duan, X. Yu, et al. 2023. “PGAM5 regulates DRP1-mediated mitochondrial fission/mitophagy flux in lipid overload-induced renal tubular epithelial cell necroptosis”. Toxicology Letters 372, 14–24.10.1016/j.toxlet.2022.10.003.

Dara, L. 2022. “No Necroptosis in Hepatocytes: The Final Nail in the Coffin?”. Gastroenterology 163, 1492–95.10.1053/j.gastro.2022.09.025.

Dara, L., Z. X. Liu, and N. Kaplowitz. 2016. “Questions and controversies: the role of necroptosis in liver disease”. Cell Death Discovery 2, 16089.10.1038/cddiscovery.2016.89.

Deepa, S. S., S. Bhaskaran, R. Ranjit, et al. 2016. “Down-regulation of the mitochondrial matrix peptidase ClpP in muscle cells causes mitochondrial dysfunction and decreases cell proliferation”. Free Radical Biology & Medicine 91, 281–92.10.1016/j.freeradbiomed.2015.12.021.

Dondelinger, Y., W. Declercq, S. Montessuit, et al. 2014. “MLKL compromises plasma membrane integrity by binding to phosphatidylinositol phosphates”. Cell Reports 7, 971–81.10.1016/j.celrep.2014.04.026.

Estrada, L. D., P. Ahumada, D. Cabrera, and J. P. Arab. 2019. “Liver Dysfunction as a Novel Player in Alzheimer’s Progression: Looking Outside the Brain”. Frontiers in Aging Neuroscience 11, 174.10.3389/fnagi.2019.00174.

Franceschi, C., M. Bonafe, S. Valensin, et al. 2000. “Inflamm-aging. An evolutionary perspective on immunosenescence”. Annals of the New York Academy of Sciences 908, 244–54.10.1111/j.1749-6632.2000.tb06651.x.

Franceschi, C., and J. Campisi. 2014. “Chronic inflammation (inflammaging) and its potential contribution to age-associated diseases”. The Journals of Gerontology. Series A, Biological Sciences and Medical Sciences 69 Suppl 1, S4–9.10.1093/gerona/glu057.

Frezza, C., S. Cipolat, O. Martins de Brito, et al. 2006. “OPA1 controls apoptotic cristae remodeling independently from mitochondrial fusion”. Cell 126, 177–89.10.1016/j.cell.2006.06.025.

Garcia-Martinez, R., and J. Cordoba. 2012. “Liver-induced inflammation hurts the brain”. Journal of Hepatology 56, 515–7. 10.1016/j.jhep.2011.11.003.

Gautheron, J., M. Vucur, and T. Luedde. 2015. “Necroptosis in Nonalcoholic Steatohepatitis”. Cellular and Molecular Gastroenterology and Hepatology 1, 264–65.10.1016/j.jcmgh.2015.02.001.

Gautheron, J., M. Vucur, F. Reisinger, et al. 2014. “A positive feedback loop between RIP3 and JNK controls non-alcoholic steatohepatitis”. EMBO Molecular Medicine 6, 1062–74.10.15252/emmm.201403856.

Ge, S. X., D. Jung, and R. Yao. 2020. “ShinyGO: a graphical gene-set enrichment tool for animals and plants”. Bioinformatics 36, 2628–29.10.1093/bioinformatics/btz931.

Goldberg, E. L., and V. D. Dixit. 2015. “Drivers of age-related inflammation and strategies for healthspan extension”. Immunological Reviews 265, 63–74. 10.1111/imr.12295.

Hanus, J., C. Anderson, and S. Wang. 2015. “RPE necroptosis in response to oxidative stress and in AMD”. Ageing Research Reviews 24, 286–98.10.1016/j.arr.2015.09.002.

Hildebrand, J. M., M. C. Tanzer, I. S. Lucet, et al. 2014. “Activation of the pseudokinase MLKL unleashes the four-helix bundle domain to induce membrane localization and necroptotic cell death”. Proceedings of the National Academy of Sciences of the United States of America 111, 15072–7.10.1073/pnas.1408987111.

Jannone, G., M. Rozzi, M. Najimi, et al. 2020. “An Optimized Protocol for Histochemical Detection of Senescence-associated Beta-galactosidase Activity in Cryopreserved Liver Tissue”. The Journal of Histochemistry and Cytochemistry: official journal of the Histochemistry Society 68, 269–78.10.1369/0022155420913534.

Kiourtis, C., A. Wilczynska, C. Nixon, et al. 2021. “Specificity and off-target effects of AAV8-TBG viral vectors for the manipulation of hepatocellular gene expression in mice”. Biol Open 10.10.1242/bio.058678.

Kjaergaard, K., A. C. Daugaard Mikkelsen, A. M. Landau, et al. 2024. “Cognitive dysfunction in early experimental metabolic dysfunction-associated steatotic liver disease is associated with systemic inflammation and neuroinflammation”. JHEP Reports: Innovation in Hepatology 6, 100992.10.1016/j.jhepr.2023.100992.

Knott, A. B., G. Perkins, R. Schwarzenbacher, and E. Bossy-Wetzel. 2008. “Mitochondrial fragmentation in neurodegeneration”. Nature Reviews. Neuroscience 9, 505–18.10.1038/nrn2417.

Kolbrink, B., F. A. von Samson-Himmelstjerna, J. M. Murphy, and S. Krautwald. 2023. “Role of necroptosis in kidney health and disease”. Nature Reviews. Nephrology 19, 300–14.10.1038/s41581-022-00658-w.

Koyama, Y., and D. A. Brenner. 2017. “Liver inflammation and fibrosis”. The Journal of Clinical Investigation 127, 55–64.10.1172/JCI88881.

Lee, H., S. B. Smith, and Y. Yoon. 2017. “The short variant of the mitochondrial dynamin OPA1 maintains mitochondrial energetics and cristae structure”. The Journal of Biological Chemistry 292, 7115–30.10.1074/jbc.M116.762567.

Lee, S. A., M. S. Kwak, S. Kim, and J. S. Shin. 2014. “The role of high mobility group box 1 in innate immunity”. Yonsei Medical Journal 55, 1165–76.10.3349/ymj.2014.55.5.1165.

Li, Q., Y. Lin, G. Liang, et al. 2023. “Autophagy and Senescence: The Molecular Mechanisms and Implications in Liver Diseases”. International Journal of Molecular Sciences 24.10.3390/ijms242316880.

Liu, J., X. Huang, M. Werner, et al. 2017. “Advanced Method for Isolation of Mouse Hepatocytes, Liver Sinusoidal Endothelial Cells, and Kupffer Cells”. Methods Mol Biol 1540, 249–58.10.1007/978-1-4939-6700-1_21.

Liu, Z., L. F. Dagley, K. Shield-Artin, et al. 2021. “Oligomerization-driven MLKL ubiquitylation antagonizes necroptosis”. The EMBO Journal 40, e103718.10.15252/embj.2019103718.

Luedde, T., N. Kaplowitz, and R. F. Schwabe. 2014. “Cell death and cell death responses in liver disease: mechanisms and clinical relevance”. Gastroenterology 147, 765–83 e4.10.1053/j.gastro.2014.07.018.

Mahajan, M., N. Bharambe, Y. Shang, et al. 2021. “NMR identification of a conserved Drp1 cardiolipin-binding motif essential for stress-induced mitochondrial fission”. Proceedings of the National Academy of Sciences of the United States of America 118.10.1073/pnas.2023079118.

Majdi, A., L. Aoudjehane, V. Ratziu, et al. 2020. “Inhibition of receptor-interacting protein kinase 1 improves experimental non-alcoholic fatty liver disease”. Journal of Hepatology 72, 627–35.10.1016/j.jhep.2019.11.008.

Mohammed, S., E. H. Nicklas, N. Thadathil, et al. 2021. “Role of necroptosis in chronic hepatic inflammation and fibrosis in a mouse model of increased oxidative stress”. Free Radical Biology & Medicine 164, 315–28.10.1016/j.freeradbiomed.2020.12.449.

Mohammed, S., P. Ohene-Marfo, C. Jiang, et al. 2025. “Impact of Mlkl or Ripk3 deletion on age-associated liver inflammation, metabolic health, and lifespan”. Geroscience.10.1007/s11357-025-01553-5.

Mohammed, S., N. Thadathil, R. Selvarani, et al. 2021. “Necroptosis contributes to chronic inflammation and fibrosis in aging liver”. Aging Cell 20, e13512.10.1111/acel.13512.

Mohar, I., K. J. Brempelis, S. A. Murray, et al. 2015. “Isolation of Non-parenchymal Cells from the Mouse Liver”. Methods Mol Biol 1325, 3–17.10.1007/978-1-4939-2815-6_1.

Najafov, A., A. K. Mookhtiar, H. S. Luu, et al. 2019. “TAM Kinases Promote Necroptosis by Regulating Oligomerization of MLKL”. Molecular cell 75, 457–68 e4.10.1016/j.molcel.2019.05.022.

Ogrodnik, M., S. Miwa, T. Tchkonia, et al. 2017. “Cellular senescence drives age-dependent hepatic steatosis”. Nature Communications 8, 15691.10.1038/ncomms15691.

Ohene-Marfo, P., H. V. M. Nguyen, S. Mohammed, et al. 2024. “Non-Necroptotic Roles of MLKL in Diet-Induced Obesity, Liver Pathology, and Insulin Sensitivity: Insights from a High-Fat, High-Fructose, High-Cholesterol Diet Mouse Model”. International Journal of Molecular Sciences 25.10.3390/ijms25052813.

Pasparakis, M., and P. Vandenabeele. 2015. “Necroptosis and its role in inflammation”. Nature 517, 311–20.10.1038/nature14191.

Passos, J. F., G. Nelson, C. Wang, et al. 2010. “Feedback between p21 and reactive oxygen production is necessary for cell senescence”. Molecular Systems Biology 6, 347.10.1038/msb.2010.5.

Preston, S. P., M. D. Stutz, C. C. Allison, et al. 2022. “Epigenetic Silencing of RIPK3 in Hepatocytes Prevents MLKL-mediated Necroptosis From Contributing to Liver Pathologies”. Gastroenterology 163, 1643–57 e14.10.1053/j.gastro.2022.08.040.

Reynoso, E., H. Liu, L. Li, et al. 2017. “Thioredoxin-1 actively maintains the pseudokinase MLKL in a reduced state to suppress disulfide bond-dependent MLKL polymer formation and necroptosis”. The Journal of Biological Chemistry 292, 17514–24.10.1074/jbc.M117.799353.

Saeed, W. K., and D. W. Jun. 2014. “Necroptosis: an emerging type of cell death in liver diseases”. World Journal of Gastroenterology 20, 12526–32.10.3748/wjg.v20.i35.12526.

Salminen, A., A. Kauppinen, and K. Kaarniranta. 2012. “Emerging role of NF-kappaB signaling in the induction of senescence-associated secretory phenotype (SASP)”. Cellular Signalling 24, 835–45.10.1016/j.cellsig.2011.12.006.

Samson, A. L., Y. Zhang, N. D. Geoghegan, et al. 2020. “MLKL trafficking and accumulation at the plasma membrane control the kinetics and threshold for necroptosis”. Nature Communications 11, 3151.10.1038/s41467-020-16887-1.

Srinivasan, S., M. Guha, A. Kashina, and N. G. Avadhani. 2017. “Mitochondrial dysfunction and mitochondrial dynamics-The cancer connection”. Biochimica et Biophysica Acta. Bioenergetics 1858, 602–14.10.1016/j.bbabio.2017.01.004.

Stahl, E. C., E. R. Delgado, F. Alencastro, et al. 2020. “Inflammation and Ectopic Fat Deposition in the Aging Murine Liver Is Influenced by CCR2”. The American Journal of Pathology 190, 372–87.10.1016/j.ajpath.2019.10.016.

Suga, S., K. Nakamura, Y. Nakanishi, et al. 2023. “An interactive deep learning-based approach reveals mitochondrial cristae topologies”. PLoS Biol 21, e3002246.10.1371/journal.pbio.3002246.

Uchida, K., K. Itakura, S. Kawakishi, et al 1995. “Characterization of epitopes recognized by 4-hydroxy-2-nonenal specific antibodies”. Archives of Biochemistry and Biophysics 324, 241–8.10.1006/abbi.1995.0036.

Victorelli, S., H. Salmonowicz, J. Chapman, et al. 2023. “Apoptotic stress causes mtDNA release during senescence and drives the SASP”. Nature 622, 627–36.10.1038/s41586-023-06621-4.

Wang, Y., W. Wei, Y. Zhang, et al. 2025. “MLKL as an emerging machinery for modulating organelle dynamics: regulatory mechanisms, pathophysiological significance, and targeted therapeutics”. Frontiers in Pharmacology 16, 1512968.10.3389/fphar.2025.1512968.

Wiley, C. D., M. C. Velarde, P. Lecot, et al. 2016. “Mitochondrial Dysfunction Induces Senescence with a Distinct Secretory Phenotype”. Cell Metabolism 23, 303–14.10.1016/j.cmet.2015.11.011.

Wu, X., K. L. Poulsen, C. Sanz-Garcia, et al. 2020. “MLKL-dependent signaling regulates autophagic flux in a murine model of non-alcohol-associated fatty liver and steatohepatitis”. Journal of Hepatology 73, 616–27.10.1016/j.jhep.2020.03.023.

Xu, J., D. Wu, S. Zhou, et al. 2023. “MLKL deficiency attenuated hepatocyte oxidative DNA damage by activating mitophagy to suppress macrophage cGAS-STING signaling during liver ischemia and reperfusion injury”. Cell Death Discovery 9, 58. 10.1038/s41420-023-01357-6.

Yan, Z., H. Yan, and H. Ou. 2012. “Human thyroxine binding globulin (TBG) promoter directs efficient and sustaining transgene expression in liver-specific pattern”. Gene 506, 289–94.10.1016/j.gene.2012.07.009.

Yang, X., K. Qiu, Y. Jiang, et al. 2024. “Metabolic Crosstalk between Liver and Brain: From Diseases to Mechanisms”. International Journal of Molecular Sciences 25.10.3390/ijms25147621.

Yang, Y. M., S. Y. Kim, and E. Seki. 2019. “Inflammation and Liver Cancer: Molecular Mechanisms and Therapeutic Targets”. Seminars in Liver Disease 39, 26–42.10.1055/s-0038-1676806.

Yoon, S., A. Kovalenko, K. Bogdanov, and D. Wallach. 2017. “MLKL, the Protein that Mediates Necroptosis, Also Regulates Endosomal Trafficking and Extracellular Vesicle Generation”. Immunity 47, 51–65 e7.10.1016/j.immuni.2017.06.001.

Yuan, J., P. Amin, and D. Ofengeim. 2019. “Necroptosis and RIPK1-mediated neuroinflammation in CNS diseases”. Nature Reviews. Neuroscience 20, 19–33.10.1038/s41583-018-0093-1.

Zhe-Wei, S., G. Li-Sha, and L. Yue-Chun. 2018. “The Role of Necroptosis in Cardiovascular Disease”. Frontiers in Pharmacology 9, 721.10.3389/fphar.2018.00721.

Zong, Y., H. Li, P. Liao, et al. 2024. “Mitochondrial dysfunction: mechanisms and advances in therapy”. Signal Transduction and Targeted Therapy 9, 124.10.1038/s41392-024-01839-8.

