## Supplementary information for "A Non-canonical Role for Hepatocyte MLKL in Promoting Mitochondrial Dysfunction and Senescence in the Aging Liver"

**Supplementary Methods**

Isolation of hepatocytes and Non-parenchymal cell (NPC) fractions

Mice were euthanized by isoflurane inhalation and *in situ* liver digestion was done. Briefly, liver was perfused through the portal vein with 20ml of perfusion buffer (HBSS, 5 mM HEPES, 0.5 mM ethylenediaminetetraacetic acid [EDTA]) and 20 ml of digestion buffer (HBSS, 5 mM HEPES, 0.5 mM CaCl2, 0.15 mg/ml Liberase (#5401127001, Millipore Sigma). The digested liver was the cut out from the abdominal cavity and transferred to cold wash buffer (Phosphate buffered saline [PBS, #10010023, ThermoFisher Scientific], 4 % fetal bovine serum (FBS, #A5669701, ThermoFisher Scientific), 0.5 mM EDTA) and the cells were released by gently tearing the liver capsule. The suspension was filtered through a 100µM cell strainer (#22-363-549, Fisher Scientific) to remove large clumps. The filtrate was centrifuged at 50g for 3 minutes to obtain the hepatocyte pellet. The supernatant containing NPC was collected and centrifuged at 600g for 10 minutes to obtain the pellet. The NPC pellet was then resuspended in resuspension buffer (PBS, 1 mM EDTA, 2 % FBS) and mixed with equal volume of 40% OptiPrep (#AXS-1114542, Cosmo Bio USA, CA, USA) and centrifuged at 1500g for 30 minutes. The well-defined interphase was collected into 10 ml of cold PBS and centrifuged at 800g, 5 minutes to obtain the enriched NPC pellet.

Western Blotting

In brief, 20 mg liver tissue were homogenized in lysis buffer [50 mM 4-(2-hydroxyethyl)-1-piperazineethanesulfonic acid (HEPES), pH 7.6; 150 mM sodium chloride; 20 mM sodium pyrophosphate; 20 mM β-glycerophosphate; 2 mM EDTA; 1% Nonidet P-40; 10% glycerol; 2 mM phenylmethylsulfonyl fluoride; and protease inhibitor cocktail (GoldBio, St Louis, MO)]. After centrifugation at 14000 x rpm for 20 min, protein concentrations in the supernatant were determined using Bio-Rad protein assay dye reagent (Bio-Rad, Hercules, CA, USA). Western blotting was performed using 30 μg protein/well. Gels after western blotting were stained with Blazin' Blue™ Protein Gel Stain (#P-810-1, Gold Biotechnology), and the blots were stained with Ponceau S solution (#P7170-1L, Sigma-Aldrich ) as per manufacturer’s instruction. Images were taken with the ChemiDoc imager (Bio-Rad) and quantified using ImageJ software (U.S. National Institutes of Health).The following primary antibodies were used: Flag (#F1804, Millipore), MLKL (#MABC604, Millipore), phospho-MLKL (#ab196436, Abcam), phospho-RIPK3 (S232 + T231) (#ab205421, Abcam), RIPK3 (#17563-1-AP, Proteintech), RIPK1 (#NBP1-77077,Novus Biologicals), Alix (#12422-1-AP, Proteintech), VPS4B (#17673-1-AP, Proteintech), TSG101 (#28283-1-AP, Proteintech), HMGB1 (#ab18256, Abcam), Calnexin (#10427-2-AP, Proteintech), LC3 (#14600-1-AP, Proteintech), Nitrotyrosine (#A-21285, Invitrogen), phospho-Drp(Ser637) (#4867, Cell Signaling), phospho-Drp(Ser616) (#PA5-64821, Thermo Scientific), Drp1(#12957-1-AP, Proteintech), Fis1 (#10956-1-AP, Proteintech), Mfn1 (#13798-1-AP, Proteintech), Mfn2 (#12186-1-AP, Proteintech), OPA1 (#ab42364, Abcam), VDAC (#ab15895, Abcam), Cleaved PARP (#13371-1-AP, Proteintech), Cleaved Caspase 3 (#9661, Cell Signaling Technology), Caspase 3 (#19677-1-AP, Proteintech), , phospho-p65 (#3033T, Cell Signaling Technology), p65 (#8242S, Cell Signaling Technology), p53 (#ab26, Abcam), PINK1 (#23274-1-AP, Proteintech), PARKIN (#14060-1-AP, Proteintech), β-actin (#A5441, Sigma-Aldrich), β-tubulin (#T5201, Sigma-Aldrich), GAPDH (#G8795, Sigma-Aldrich). HRP-linked anti-rabbit IgG, HRP-linked anti-mouse IgG and HRP-linked anti-rat IgG secondary antibodies were from Cell Signaling Technology.

Detection of 4-Hydroxynonenal (4-HNE) Adducts

For the detection of 4-HNE modified proteins, equal amounts of protein (40 μg/lane) were separated by SDS–PAGE, transferred to polyvinylidene difluoride membranes and treated with 250 mM sodium borohydride in 100 mM (3-(N-morpholino)propanesulfonic acid, MOPS), pH 8.0 for 15 min. The membrane was washed with water, followed by Tris buffered saline with Tween 20 (TBS-T), and blocked with 5% non-fat milk/TBS-T. The membrane was incubated with a 1:2000 dilution of polyclonal antibody against 4-HNE (gift from Dr. Luke Szweda, Oklahoma Medical Research Foundation). The antibody recognizes cysteine, lysine, and histidine 4-HNE protein adducts and is highly specific to 4-HNE derived protein adducts. This was followed by incubation with anti-rabbit IgG HRP conjugated antibody and the blot was developed using ECL Western Blotting Substrate (ThermoFisher Scientific, Waltham, MA). Images were taken using a ChemiDoc imager (Bio-Rad) and quantified using ImageJ software (U.S. National Institutes of Health, Bethesda, MD, USA).

Immunofluorescence (IF) Staining

Paraffin sections were deparaffinized and rehydrated through xylene, graded ethanol (100%, 95%, 70%, 50%), and deionized water. Heat-induced antigen retrieval was performed, followed by permeabilization with 0.1% Tween-20 in PBS for 10 min. After PBS washing, sections were blocked with 2% BSA at room temperature for 60 min. Primary antibody F4/80 (#NB600-404, Novus Biologicals) was diluted in 1% BSA (1: 400) and incubated at 4°C overnight. After PBS washing, Donkey anti-rat Alexa Fluor 488 (Abcam) secondary antibody was applied for 60 min at room temperature. The sections were washed with PBS and mounted with ProLong™ Diamond antifade mountant with DAPI (#P36962, Thermofisher scientific). Images were taken with Nikon TE2000-E microscope at 200 magnification, 3 random non-overlapping fields per sample were recorded. The percentage area of positive signal (green) per field was quantified using ImageJ software and was normalized to DAPI. The data are presented as fold change after normalization to the control mice.

Quantitative Real-Time PCR (qPCR)

Twenty mg frozen liver tissue was used to isolate RNA using RNeasy Mini Kit (#74106, Qiagen) as per manufacturer’s instruction. First-strand cDNA was synthesized, and the real-time-PCR was performed. The primers used are listed in Table S1. The calculations were conducted by a comparative method (2^−ΔΔCt^). β-microglobulin, β-actin, or hypoxanthine phosphoribosyltransferase 1 (HPRT) was used as controls.

Label Free Quantitative Proteomic Analysis of Liver

A total of 100 µg of liver lysate (n=5/group) was digested in-solution using Trypsin/LysC (#V5071, Promega, WI, USA) following the manufacturer’s instructions. Following digestion, the peptides were desalted with C18 Sep-Pak Plus cartridges (Waters, MA, USA). 100 µL of 0.1% formic acid was used to reconstitute the dried tryptic peptides to a final concentration of 1 µg/µL. Resuspended tryptic peptides (2 μL) were loaded onto a C18 trap column (150 μm × 3 cm, 3 μm resin, Acclaim™ PepMap™ 100 C18 HPLC Column, Thermo Scientific™, USA) using mobile phase A (0.1% formic acid in LC-MS grade water). The flow rate was 3 μl/min for 10 min, and separate peptides on an EASY-Spray™ HPLC analytical column (3 μm x 75 μm × 15 cm, Catalog # ES900 Thermo Scientific™, USA) at the rate of 350 nL/min. The total LC-MS/MS run time was 1 hour. The LC-MS/MS analysis was performed with a Dionex UltiMate® 3000 UHPLC system (Thermo Fisher Scientific, CA, USA) coupled to a Q Exactive HF-X mass spectrometer (Thermo Fisher Scientific, Waltham, MA).

The RAW MS files were searched against the UniProt reviewed mouse (Taxon ID: 10090) protein database, the Sequest algorithm within Proteome Discoverer v 2.4 (Thermo Fisher Scientific, San Jose, CA). Parameters used are listed as follows: trypsin enzyme cleavage specificity, 2 possible missed cleavages, 10 ppm mass tolerance for precursor ions, and 0.02 Da mass tolerance for fragment ions. The above search parameters permit dynamic modification of methionine oxidation (+15.9949 Da) and static modification of carbamidomethylation (+57.0215 Da) on cysteine. After the database search, peptide assignments were filtered down to a 1% FDR (false discovery rate). Label-free quantitation was performed using the Minora algorithm and the adjoining bioinformatics tools available in Proteome Discoverer. Statistically significant is defined as a 1.5-fold increase or decrease in abundance with a p-value <0.05.

Targeted Mitochondrial Proteomics

20 μg of total liver homogenate was separated ~1.5 cm into a 12.5% SDS-PAGE gel (Criterion, Bio-Rad, Berkeley, CA, USA). Gels were fixed and stained using GelCode Blue stain (Pierce, Appleton, WI, USA), and the entire lane was excised into ~1 mm³ sections. Gel pieces were sequentially washed, reduced with dithiothreitol (DTT), alkylated with iodoacetamide, and digested with trypsin. The resulting peptides were extracted using 50% methanol/10% formic acid, dried, reconstituted in 1% acetic acid, and analyzed via selected reaction monitoring (SRM) using a triple quadrupole mass spectrometer (TSQ Quantiva, Thermo Scientific, Waltham, MA, USA) coupled to a splitless capillary HPLC system (Ultimate 3000, Thermo Scientific, Waltham, MA, USA). Peptide quantification was performed using Skyline software, which aligned collision-induced dissociation transitions and quantified chromatographic peak areas. Protein abundance was calculated as the sum of all monitored peptide responses. Relative protein levels were normalized to a BSA internal standard and verified using normalization to housekeeping proteins.

Untargeted Lipidomics Analysis

Approximately 50 mg of liver tissue was homogenized in 1.5 mL chloroform:methanol (2:1, v/v) and 0.5 mL ultrapure water. Samples were ground, vortexed, and sonicated at 4 °C for 30 minutes, followed by centrifugation at 3,000 rpm for 10 minutes. The lower phase was collected, dried under nitrogen, and reconstituted in 200 μL isopropanol:methanol (1:1, v/v) containing LPC (12:0) as an internal standard. After centrifugation at 12,000 rpm for 10 minutes at 4 °C, supernatants were used for UPLC-MS analysis. QC samples were prepared by pooling equal volumes from each sample. Lipid separation was conducted on an ACQUITY UPLC BEH C18 column using a gradient of solvent A (60% acetonitrile, 40% water, 10 mM ammonium formate) and solvent B (90% isopropanol, 10% acetonitrile, 10 mM ammonium formate) at 0.3 mL/min. Analysis was performed using a Q Exactive mass spectrometer (Thermo Scientific) in both ESI+ and ESI– modes with standard parameters. Raw data were processed with LipidSearch for peak alignment and identification. Multivariate analysis was performed using SIMCA-P (v14.1), including PCA, PLS-DA, and OPLS-DA. Significantly altered lipids were identified using VIP > 1.5, fold change > 2, and p < 0.05.

Measurement of Mitochondrial Respiration

To measure mitochondrial respiration, AML12 cells were seeded at a density of 10,000 cells/well kept at 37 °C incubator with 5% CO_2_ for 24 h. The cells were transfected with either pcDNA or pcDNA-MLKL-Flag as described above. Twenty four hours post transfection, cells were changed to assay media [XF base medium (Cat#103575-100, Agilent ) containing 25 mM glucose, 1 mM sodium pyruvate and 1 mM L-glutamine, pH 7.4] and kept in a 37 °C in an incubator without CO_2_ for 60 min. Oxygen consumption rate (OCR) was recorded when cells were metabolically perturbed by the sequential injections of oligomycin (1.5μM), carbonyl cyanide-4-(trifluoromethoxy) phenylhydrazone (FCCP) (2μM) and Rotenone/antimycin A (0.5μM) final concentration). From the obtained OCR values calculations were made as follows: basal respiration (third basal measurement), ATP-linked (difference between basal respiration rate and oligomycin-induced respiration), proton leak (difference between oligomycin-induced and antimycin A-induced respirations), maximal respiration (maximum rate after FCCP injection), reserve capacity (maximal respiration–basal respiration) and non-mitochondrial respiration.

Live-Cell Imaging Using Photoactivatable GFP and LAMP1/Mt-Mscarlet Reporters

AML12 cells were cultured on either 14mm or 20mm coverslips in a 35mm dish (Mattek) that were either tissue culture treated or coated with 1mg/ml poly-d-lysine (PDL; Sigma) at a 1:20 dilution. Cells were transfected using JetPrime (PolyPlus) with 1µg of total DNA and 2µl of transfection reagent in 100µl of buffer per dish/well to be transfected. For MLKL overexpression, 500ng of DDK tagged MLKL (pCAG MLKL-DDK) was transfected alongside 50ng of the fluorescent reporter being used for each experiment (pCAG 2xmt-paGFP p2a 2xmt-mScarlet, pCAG 2xmt-mScarlett or pCAG Lamp1-mEmerald) and the additional 450ng was made up with a pLKO empty vector (ThermoFisher). Following transfection, media was swapped with fresh media to remove transfection reagent after 4-6 hours of incubation at 37 degrees Celsius.

Following overexpression, live time lapse imaging of the cells was done every 15 seconds for 10 minutes on a Nikon Ti2 widefield system equipped with a Hammamatsu ORCA-Fusion CMOS camera, a custom penta-band cube for 378/474/554/635/735 excitation with an Aura III light engine, and 60x (1.4NA) oil objective, and live imaging chamber from OXO (UNO-T-H-CO2) with objective warmer. In addition, a 405nm laser (LUN-F, 50mW) with XY galvo control (Opti-microscan) is connected to perform targeted ROI based stimulation. The whole system is controlled by Nikon Elements. For photo-activation experiments, 5µm by 5µm ROIs were selected in each cell imaged. A before stimulation image was taken, then each box was individually stimulated at 2% laser power for 100µs followed immediately by the time series imaging. For analysis of mitochondrial dynamics, a square ROI of 5µm by 5µm was used and ROI intensity statistics were quantified across the time course. Data was plotted over the entire time course, with comparisons made with end points and slopes.

Exosome Isolation and Nanoparticle Tracking Analysis

AML12 cells were transfected with either control pcDNA or pcDNA-MLKL-Flag as described above. HepG2 cells (#HB-8065, ATCC) were transfected with either siControl (Silencer Select Negative Control No. 1 siRNA, # 4390843, Thermofisher) or siMLKL (Silencer Select siRNA against human MLKL, # 4392420, Thermofisher), and 6-hour post transfection, cell culture medium was changed into exosome free medium 24 hour before harvest. Exosome isolation was performed using the Total exosome isolation kit (# 4478359, Invitrogen) as per manufacturer’s instruction. Briefly, collected cell culture media was centrifuged at 2000 x g for 30min. The supernatant was mixed with the total exosome isolation reagent and incubated overnight at 4 °C. After centrifugation at 10,000 x g for 60min at 4°C, exosomes were contained in the pellet at the bottom of the tube. The pellet was then resuspended in PBS, and analyzed using Nanosight (NTA-NS300, Malvern Panalytical) for particle size and concentration analysis. The quantification was done with GraphPad Prism, and the exosome concentration is presented as fold change to the control group.

Bioinformatics

For lipid metabolites, significantly altered concentrations between experimental conditions were identified using specific functions provided by the Bioconductor package limma (http://www.bioconductor.org/packages/release/bioc/html/limma.html). Moderated t-statistics, relying on empirical Bayes shrinkage of the standard errors toward a common value, were computed using the lmfit function of the package. The P-values corresponding to the moderated t-statistics were adjusted for multiple testing by false discovery rates (FDR) method of Benjamini, Hochberg, and Yekutieli. FDR below 0.05 was used as filtering criteria for significantly altered metabolites. Families of metabolites significantly enriched in altered metabolites were further identified using Fisher test.

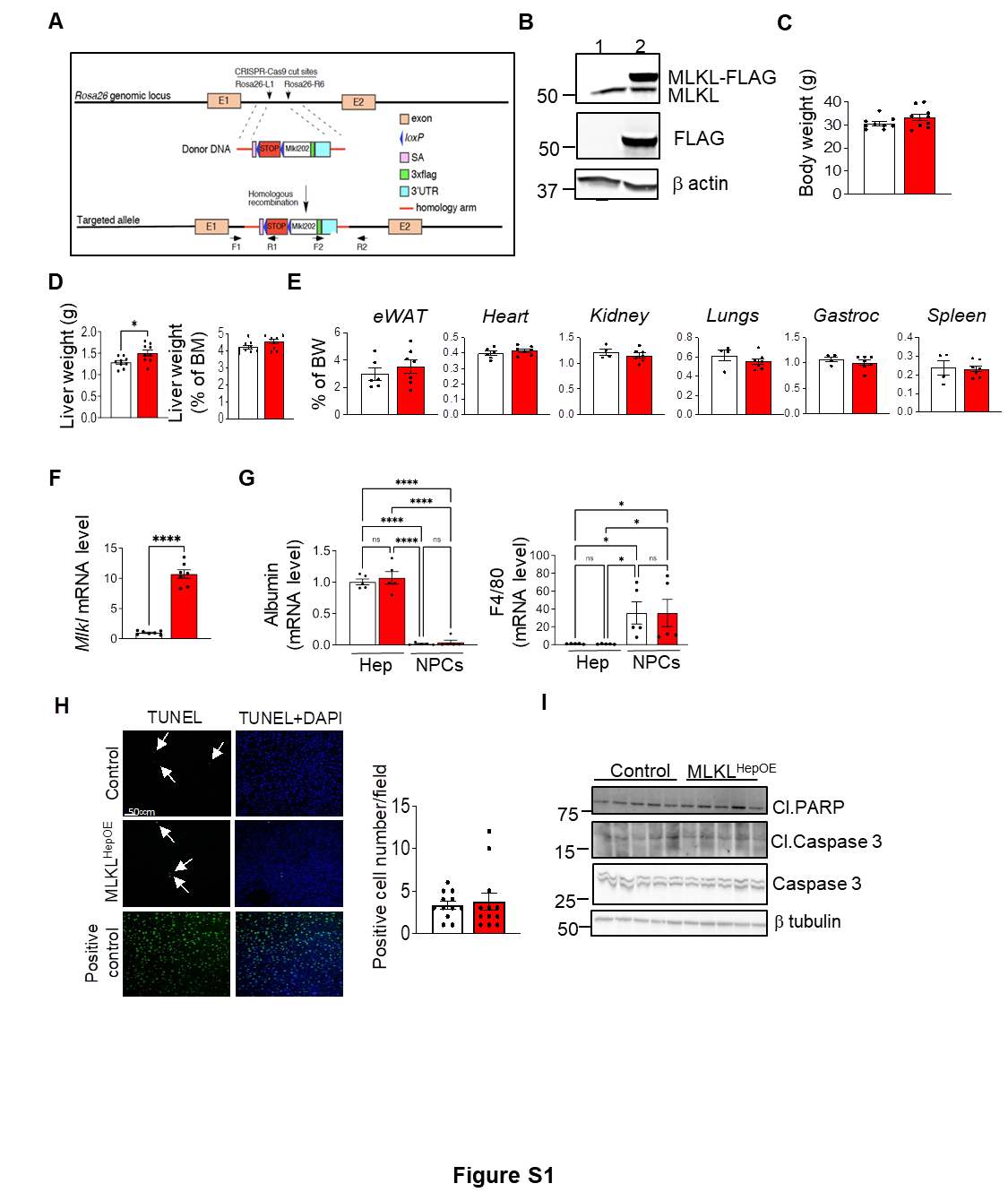

**Figure S1. (A)** Schematic representation of the Mlkl genomic locus and the strategy used to generate MLKL^HepOE^ mice. Exons are depicted as boxes, with loxP sites flanking exon 2 shown as blue triangles. To enable conditional overexpression, a donor DNA construct encoding a stop codon followed by the *Mlkl*-3xFlag sequence was inserted into the *Rosa26* locus via CRISPR-Cas9–mediated homologous recombination. **(B)** Western blot analysis of liver tissue lysates from control and MLKL^HepOE^ mice showing MLKL overexpression, 2 weeks post virus injection. β-actin serves as a loading control. **(C)** Body weight comparison between control and MLKL^HepOE^ mice. **(D)** Absolute liver weight (g) and liver-to-body weight ratio (% of body mass) in control and MLKL^HepOE^ mice **(E)** Weights of epididymal white adipose tissue (eWAT), heart, kidney, lungs, gastrocnemius muscle (Gastro), and spleen, expressed as a percentage of body weight (% BW), in control and MLKL^HepOE^ mice**. (F)** qPCR analysis of Mlkl mRNA levels in the livers of control and MLKL^HepOE^ mice. **(G)** qPCR analysis of albumin and F4/80 levels in isolated hepatocytes (Hep) and non-parenchymal cells (NPCs) isolated from control and MLKL^HepOE^ mice. **(H)** *Left:* Representative images of TUNEL staining (green) with DAPI counterstaining (blue) in liver sections from control, MLKL^HepOE^ mice, and a positive control (DNase treated liver sections). Arrows indicate TUNEL-positive nuclei. ***Right:*** Quantification of TUNEL-positive cells per field (4 fields/section; n = 3 animals per group). **(I)** Western blot analysis of cleaved PARP (Cl. PARP), cleaved caspase 3 (Cl. Caspase 3), and total caspase 3 protein levels in liver lysates of control and MLKL^HepOE^ mice. β-tubulin serves as a loading control. Control (white) and MLKL^HepOE^ (red). Data are presented as mean ± SEM from n=7-9mice/group. Statistical significance was determined by two-tailed unpaired t-test for C-F, H or One-way ANOVA for G . ****p < 0.0001, *p < 0.05, ns: not significant.

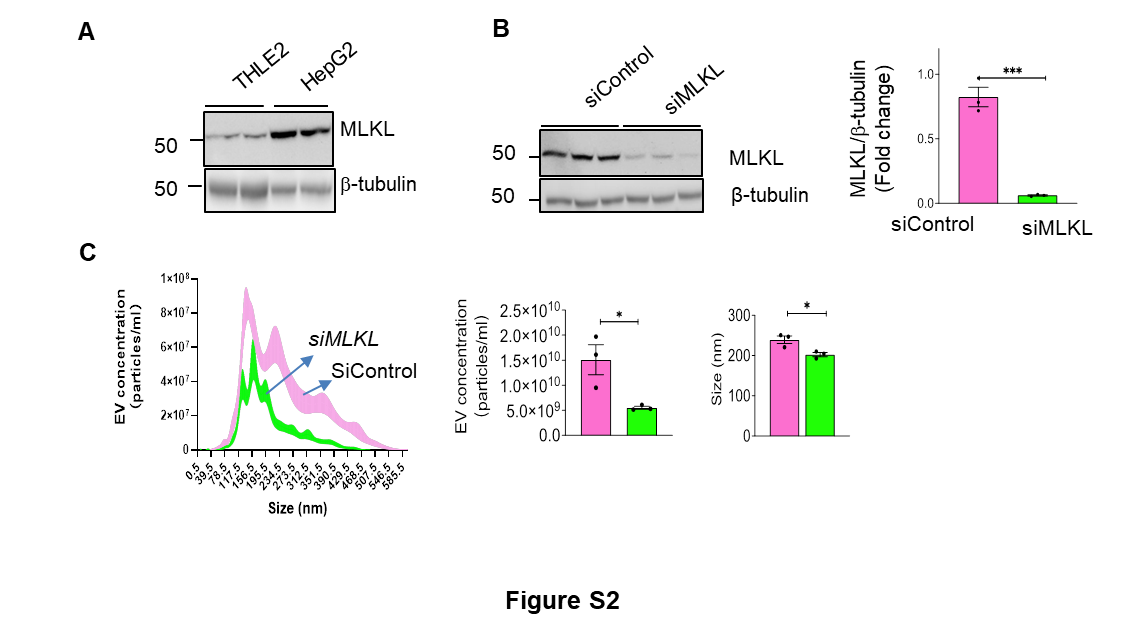

**Figure S2. (A)** Western blot analysis of basal MLKL protein expression in THLE2 (normal) and HepG2 (liver cancer) cell lines. β-tubulin is used as a loading control. **(B)** *Left:* Western blot analysis of MLKL protein expression in HepG2 cells transfected with either siControl or siMLKL. β-tubulin serves as a loading control. *Right:* The bar graph shows quantification of MLKL protein levels normalized to β-tubulin, presented as fold change (n=3 independent experiments). (**C)** Nanoparticle Tracking Analysis (NTA) of EVs isolated from HepG2 cells transfected with siControl (pink) or siMLKL (green). The left panel shows the EV size distribution, the middle panel presents total EV concentration (particles/ml), and the right panel displays average EV diameter (nm) (n = 3 independent experiments). Statistical significance was determined by two-tailed unpaired t-test. ***p < 0.001, *p<0.05.

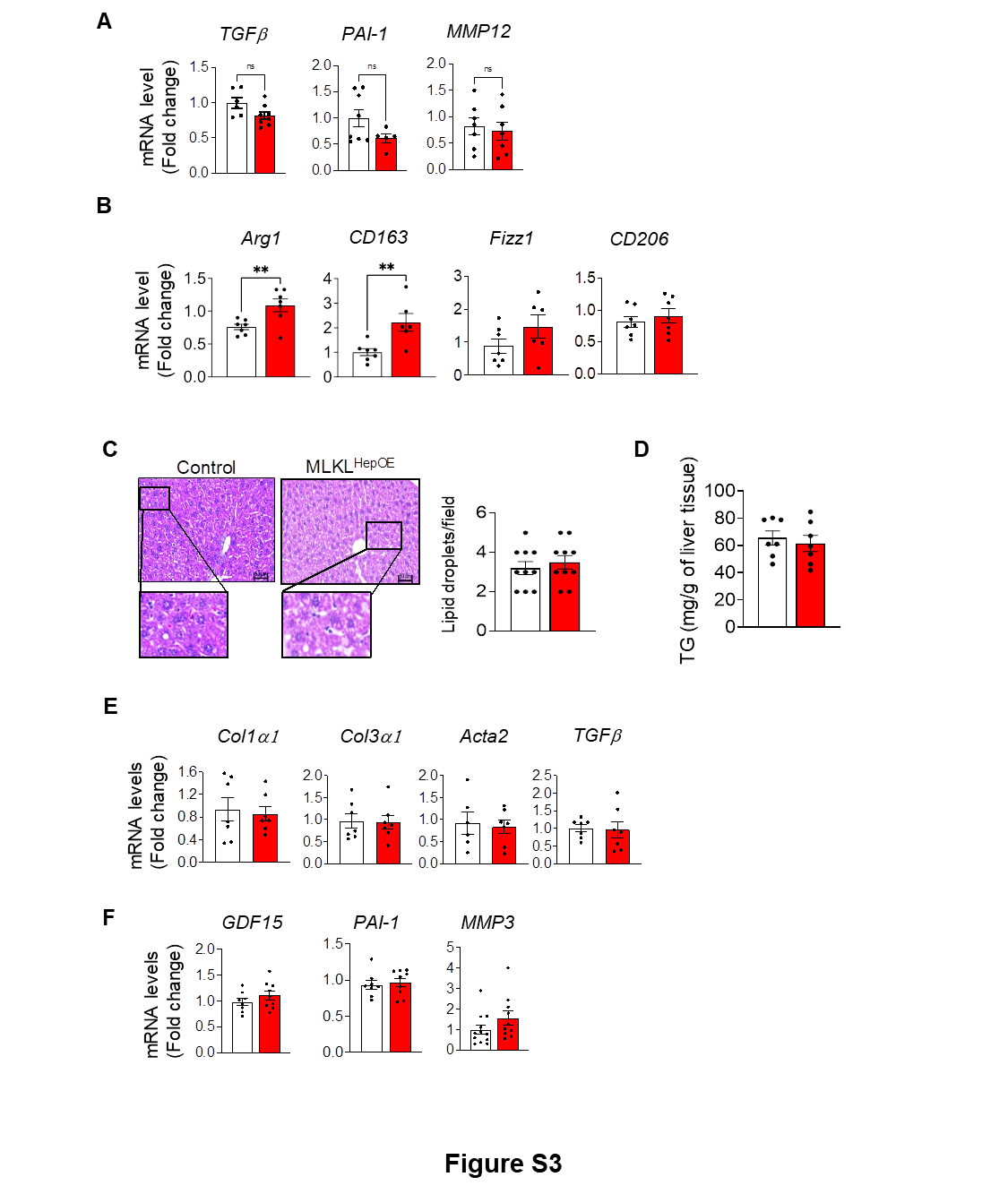

**Figure S3. A-E: Data from control and** MLKL^HepOE^ **mice livers. (A) qPCR analysis of SASP factors. (B) qPCR analysis of M2 macrophage markers (C) Representative images of H&E-stained liver sections. Scale bar=50 μm. Magnification: 100X. The graph on the right shows quantification of lipid droplets per field (n = 3-4 fields per animal, 3 animals per group). (D) Triglyceride (TG) content (mg/g) of liver tissue. (E) qPCR analysis of fibrosis-related genes. (n = 6-8/ group) (F) qPCR analysis of MLKL and SASP factors from control (white) and MLKL overexpressing (red) AML12 cells.** Data from control (white) and MLKL^HepOE^ (red) mice (n = 5/group). Statistical significance was determined by two-tailed unpaired t-test. **p<0.01, ns: not significant.

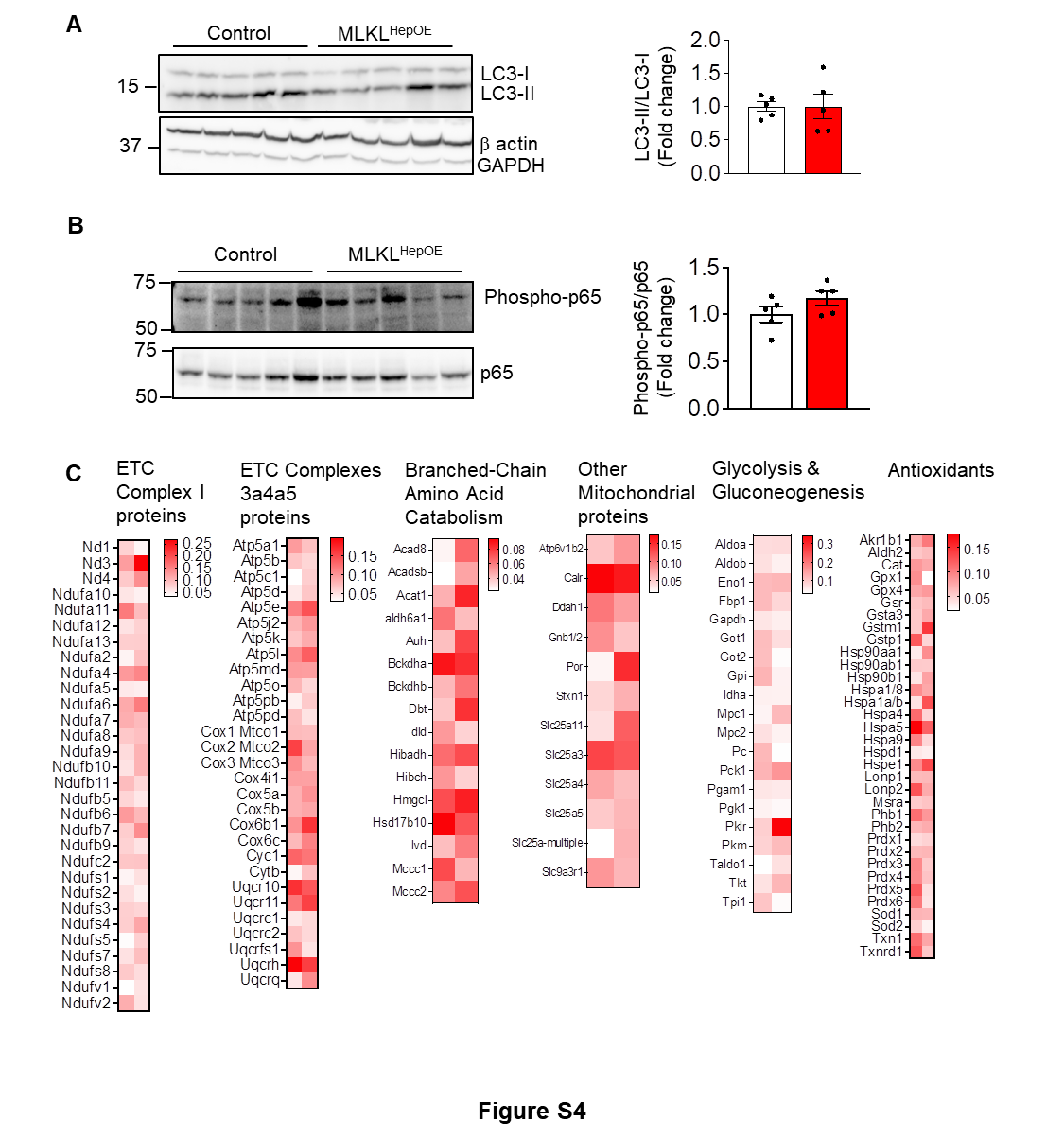

**Figure S4.** Western blot analysis of liver tissue lysates from control (white) and MLKL^HepOE^ (red) mice showing **(A) *Left:*** LC3-I, LC3-II and β-actin (loading control), *Right*: Ratio of LC3-II/LC3-I represented as fold change, **(B)** *Left:* Phosphorylated p65 (Phospho-p65), total p65, *Right:* Phospho-p65/p65 ratio, presented as fold change. (**C)** Heatmap of differentially expressed proteins in the livers of MLKL^HepOE^ mice compared to control mice, as determined by targeted proteomics. Each column represents a biological replicate for the control and MLKL^HepOE^ groups. The color scale indicates the log2 fold change; red indicates upregulation and white indicates no significant change. All data are presented as mean ± SEM from n=5mice/group**.** Statistical significance was determined by two-tailed unpaired t-test.

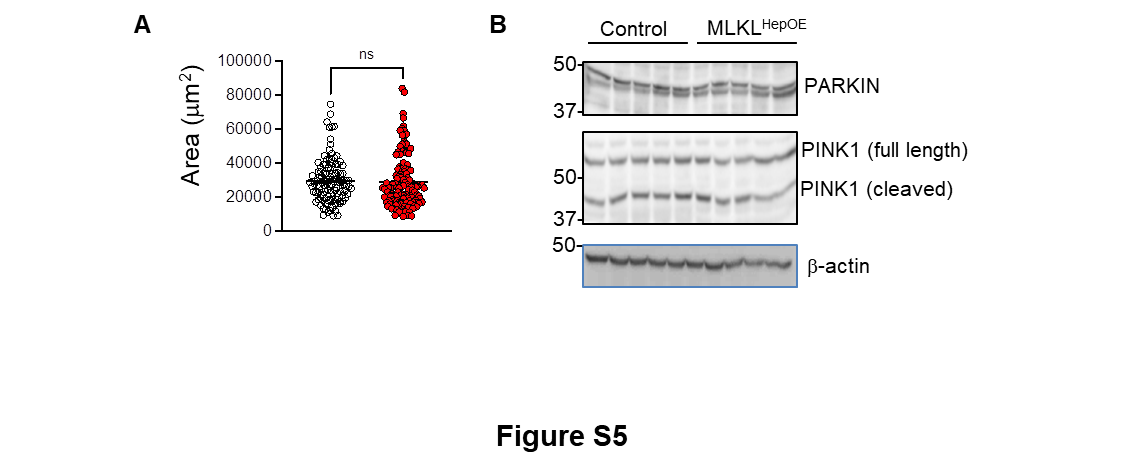

**Figure S5. (A) Mitochondrial area (µm²) in liver sections of control (black circles) and MLKL^HepOE^ (red dots) mice. Each dot represents an individual mitochondrion. (B) Western blot analysis showing protein levels of PARKIN, full-length PINK1, and cleaved PINK1 in liver lysates from control and MLKL^HepOE^ mice. β-Actin** serves as a loading control. All data are presented as mean ± SEM from n=5mice/group**.** Statistical significance was determined by two-tailed unpaired t-test ns: not significant.

**Table S1**: List of primer sequences used for quantitative RT PCR analysis

**Table S2**: Details of patient samples used for human plasma MLKL analysis

**Table S3**: List of significantly altered proteins in untargeted label free quantitative proteomic analysis

**Table S4**: List of proteins identified by targeted mitochondrial proteomics

**Table S5**: List of differentially regulated lipid species in lipidomic analysis
