## Supplementary Table 1 for "A Non-canonical Role for Hepatocyte MLKL in Promoting Mitochondrial Dysfunction and Senescence in the Aging Liver"

| **Primer name** | **Forward primer sequence** | **Reverse primer sequence** |
| --- | --- | --- |
| Mlkl | 5'-CTGAGGGAACTGCTGGATAGAG-3' | 5'-CGAGGAAACTGGAGCTGCTGAT-3' |
| Acta2 | 5'-CTGACAGAGGCACCACTGAA-3' | 5'-CATCTCCAGAGTCCAGCACA-3' |
| Albumin | 5’-GCGCAGATGACAGGGCGGAA-3’ | 5’-GTGCCGTAGCATGCGGGAGG-3’ |
| Arg1 | 5'-CAAGACAGGGCTCCTTTCAG-3' | 5'-AAGCAAGCCAAGGTTAAAGC-3' |
| CD11c | 5’-CTGGATAGCCTTTCTTCTGCTG-3’ | 5’-GCACACTGTGTCCGAACTC-3’ |
| CD163 | 5’- GGCTAGACGAAGTCATCTGCAC-3’ | 5’-CTTCGTTGGTCAGCCTCAGAGA-3’ |
| CD206 | 5’-ACTACACACTCATCCATTACAACCAA-3’ | 5'-GGCACCTATCACAATCAGGAGGA-3' |
| CD68 | 5’-CCACAGGCAGCACAGTGGAC-3’ | 5’-TCCACAGCAGAAGCTTTGGCCC-3’ |
| Col1α1 | 5’-GCTCCTCTTAGGGGCCACT-3’ | 5’-CCACGTCTCACCATTGGGG-3’ |
| Col3α1 | 5’-CTGTAACATGGAAACTGGGGAAA-3’ | 5’- CCATAGCTGAACTGAAAACCACC-3’ |
| CXCL1 | 5’- ACCGAAGTCATAGCCACACTC-3’ | 5’- CTCCGTTACTTGGGGACACC-3’ |
| CXCL2 | 5’-CCTGGTTCAGAAAATCATCCA-3’ | 5’-CTTCCGTTGAGGGACAGC-3’ |
| CXCL3 | 5’- AGTGTGAATGTAAGGTCCCC-3’ | 5’- GGAAGTGTCAATGATACGCTG-3’ |
| CXCL10 | 5’- ATCATCCCTGCGAGCCTATCCT-3’ | 5’- GACCTTTTTTGGCTAAACGCTTTC-3’ |
| F4/80 | 5'-CCCCAGTGTCCTTACAGAGTG-3' | 5'-GTGCCCAGAGTGGATGTCT-3' |
| Fizz1 | 5'-AGGAACTTCTTGCCAATCCA-3' | 5'-ACAAGCACACCCAGTAGCAG-3' |
| GDF15 | 5′-GTTAGCCAAAGACTGCCACTG-3′ | 5′-CCTTGAGCCCATTCCACA-3′ |
| IFN gamma | 5'-CTTCTTCAGCAACAGCAAGGCG-3' | 5'-ATGCTTGGCGCTGGACCTGTG-3' |
| IL1β | 5’-AGGTCAAAGGTTTGGAAGCA-3’ | 5’-TGAAGCAGCTATGGCAACTG-3’ |
| MMp12 | 5’- TGCACTCTGCTGAAAGGAGTCT-3’ | 5’- GTCATTGGAATTCTGTCCTTTCCA-3’ |
| MMP3 | 5’- GTTGGAGAACATGGAGACTTTGT-3’ | 5’- CAAGTTCATGAGCAGCAACCA-3’ |
| P15 | 5'-ATCCCAACGCCCTGAACCGCT-3' | 5'-AGTTGGGTTCTGCTCCGTGGAG-3' |
| p16 | 5’-CCCAACGCCCCGAACT-3’ | 5’-GCAGAAGAGCTGCTACGTGAA-3’ |
| p21 | 5’-GTCAGGCTGGTCTGCCTCCG-3’ | 5’-CGGTCCCGTGGACAGTGAGCAG-3’ |
| p53 | 5’-GTATTTCACCCTCAAGATCC-3’ | 5’-TGGGCATCCTTTAACTCTA-3’ |
| Pai-1 | 5′-GACACCCTCAGCATGTTCATC-3′ | 5′-AGGGTTGCACTAAACATGTCAG-3′ |
| Ripk3 | 5'-GAAGACACGGCACTCCTTGGTA-3' | 5'-CTTGAGGCAGTAGTTCTTGGTGG-3' |
| TGFβ | 5’-ACCATGCCAACTTCTGTCTGGGAC-3’ | 5’-ACAACTGCTCCACCTTGGGCTTG-3’ |
| TLR4 | 5'-ATGGCATGGCTTACACCACC-3' | 5'-GAGGCCAATTTTGTCTCCACA-3' |
| TNFα | 5’-CACAGAAAGCATGATCCGCGACGT-3 | 5’- CGGCAGAGAGGAGGTTGACTTTCT-3’ |

**Table S1**: List of primer sequences used for quantitative real time PCR analysis
