## Supplementary Table 2 for "A Non-canonical Role for Hepatocyte MLKL in Promoting Mitochondrial Dysfunction and Senescence in the Aging Liver"

| **Sample Number** | **DX Sample ID Number** | **Sex** | **Race** | **BMI** | **AGE** |
| --- | --- | --- | --- | --- | --- |
| 1 | NAS5924036 | F | White | 41.8 | 52 |
| 2 | NAS5924037 | M | White | 40.2 | 54 |
| 3 | NAS5924042 | F | White | 40 | 67 |
| 4 | NAS5924046 | F | White | 45.9 | 57 |
| 5 | NAS5924032 | F | African American | 26.5 | 60 |
| 6 | NAS5924056 | F | White | 40.5 | 51 |
| 7 | NAS5924035 | F | White | 53.1 | 45 |
| 8 | NAS5924049 | F | White | 24.74 | 69 |
| 9 | NAS5924015 | M | White | 46.4 | 69 |
| 10 | NAS5924020 | M | White | 45.8 | 30 |
| 11 | NAS5924068 | F | White | 24.1 | 53 |
| 12 | NAS5924061 | F | White | 41 | 49 |
| 13 | NAS5924058 | F | White | 28.2 | 65 |
| 14 | NAS5924005 | M | White | 40.1 | 56 |
| 15 | NAS5924039 | F | White | 40.1 | 48 |
| 16 | NAS5924024 | F | White | 50.3 | 63 |
| 17 | NAS5924057 | F | White | 40.1 | 49 |
| 18 | NAS5924003 | F | African American | 43.4 | 55 |
| 19 | NAS5924014 | F | White | 52.1 | 55 |
| 20 | NAS5924036 | F | White | 40.2 | 52 |
| M – Male;  F - Female |  |  |  |  |  |

**Table S2: Details of patient samples used for human plasma MLKL analysis**
