## Supplementary Table 3 for "A Non-canonical Role for Hepatocyte MLKL in Promoting Mitochondrial Dysfunction and Senescence in the Aging Liver"

| **Description**  **Table S3:** List of significantly altered proteins in untargeted label free quantitative proteomic analysis | **Gene Name** | **Abundance Ratio: (Test) / (Control)** | **Abundance Ratio Adj. P-Value: (Test) / (Control)** |
| --- | --- | --- | --- |
| **Significantly upregulated** | | | |
| Hemoglobin beta chain subunit OS=Mus musculus OX=10090 GN=HBB1 PE=2 SV=1 | HBB1 | 100 | 2.75E-16 |
| Maleylacetoacetate isomerase OS=Mus musculus OX=10090 GN=Gstz1 PE=1 SV=1 | Gstz1 | 100 | 2.75E-16 |
| Myoglobin OS=Mus musculus OX=10090 GN=Mb PE=1 SV=3 | Mb | 100 | 2.75E-16 |
| Calcium-transporting ATPase OS=Mus musculus OX=10090 GN=Atp2a3 PE=1 SV=1 | Atp2a3 | 100 | 2.75E-16 |
| Polyadenylate-binding protein 2 OS=Mus musculus OX=10090 GN=Pabpn1 PE=1 SV=1 | Pabpn1 | 100 | 2.75E-16 |
| Complex I-SGDH (Fragment) OS=Mus musculus OX=10090 GN=Ndufb5 PE=1 SV=1 | Ndufb5 | 100 | 2.75E-16 |
| Alanine aminotransferase 2 OS=Mus musculus OX=10090 GN=Gpt2 PE=1 SV=1 | Gpt2 | 100 | 2.75E-16 |
| Band 4.1 OS=Mus musculus OX=10090 GN=Epb41 PE=1 SV=1 | Epb41 | 100 | 2.75E-16 |
| Cytochrome P450 4A14 OS=Mus musculus OX=10090 GN=Cyp4a14 PE=1 SV=1 | Cyp4a14 | 8.416 | 2.75E-16 |
| LIM and SH3 domain protein 1 (Fragment) OS=Mus musculus OX=10090 GN=Lasp1 PE=1 SV=1 | Lasp1 | 7.511 | 2.75E-16 |
| Monocarboxylate transporter 1 OS=Mus musculus OX=10090 GN=Slc16a1 PE=2 SV=1 | Slc16a1 | 4.325 | 2.75E-16 |
| Cordon-bleu protein-like 1 OS=Mus musculus OX=10090 GN=Cobll1 PE=1 SV=2 | Cobll1 | 3.972 | 2.75E-16 |
| Peroxisomal acyl-coenzyme A oxidase 1 (Fragment) OS=Mus musculus OX=10090 GN=Acox1 PE=1 SV=1 | Acox1 | 3.585 | 2.75E-16 |
| Unconventional myosin-Ib OS=Mus musculus OX=10090 GN=Myo1b PE=1 SV=3 | Myo1b | 3.049 | 2.75E-16 |
| Ubiquinone biosynthesis protein (Fragment) OS=Mus musculus OX=10090 GN=Coq9 PE=1 SV=2 | Coq9 | 3.003 | 2.75E-16 |
| Perilipin-3 OS=Mus musculus OX=10090 GN=Plin3 PE=1 SV=1 | Plin3 | 2.903 | 2.75E-16 |
| Histone-binding protein RBBP4 OS=Mus musculus OX=10090 GN=Rbbp4 PE=1 SV=5 | Rbbp4 | 2.892 | 2.75E-16 |
| CD2-associated protein OS=Mus musculus OX=10090 GN=Cd2ap PE=1 SV=1 | Cd2ap | 2.836 | 2.75E-16 |
| Derlin OS=Mus musculus OX=10090 GN=Derl1 PE=2 SV=1 | Derl1 | 2.792 | 2.75E-16 |
| High mobility group protein HMG-I/HMG-Y OS=Mus musculus OX=10090 GN=Hmga1 PE=1 SV=1 | Hmga1 | 2.705 | 2.75E-16 |
| BolA-like protein 1 OS=Mus musculus OX=10090 GN=Bola1 PE=1 SV=1 | Bola1 | 2.418 | 2.75E-16 |
| Clusterin OS=Mus musculus OX=10090 GN=Clu PE=2 SV=1 | Clu | 2.361 | 2.75E-16 |
| Dynein light intermediate chain (Fragment) OS=Mus musculus OX=10090 GN=Dync1li1 PE=2 SV=1 | Dync1li1 | 2.312 | 2.75E-16 |
| Clathrin interactor 1 OS=Mus musculus OX=10090 GN=Clint1 PE=1 SV=1 | Clint1 | 2.264 | 1.16E-14 |
| Uncharacterized protein OS=Mus musculus OX=10090 GN=F10 PE=1 SV=1 | F10 | 2.263 | 1.16E-14 |
| Cytochrome P450 3A41 OS=Mus musculus OX=10090 GN=Cyp3a41a PE=1 SV=2 | Cyp3a41a | 2.239 | 2.83E-14 |
| AP-2 complex subunit mu (Fragment) OS=Mus musculus OX=10090 GN=Ap2m1 PE=2 SV=1 | Ap2m1 | 2.191 | 1.47E-13 |
| Uncharacterized protein OS=Mus musculus OX=10090 GN=Grn PE=2 SV=1 | Grn | 2.178 | 2.21E-13 |
| Beta-2-globin (Fragment) OS=Mus musculus OX=10090 GN=Hbb-b2 PE=2 SV=1 | Hbb-b2 | 2.112 | 2.35E-12 |
| Exportin-2 (Fragment) OS=Mus musculus OX=10090 GN=Cse1l PE=1 SV=1 | Cse1l | 2.047 | 2.36E-11 |
| MKIAA4115 protein (Fragment) OS=Mus musculus OX=10090 GN=G3bp1 PE=2 SV=1 | G3bp1 | 2.037 | 3.41E-11 |
| **Significantly down regulated proteins** | | | |
| Non-specific serine/threonine protein kinase OS=Mus musculus OX=10090 GN=Slk PE=2 SV=1 | Slk | 0.47 | 1.88E-13 |
| ATP-dependent Clp protease proteolytic subunit (Fragment) OS=Mus musculus OX=10090 GN=Clpp PE=2 SV=1 | Clpp | 0.458 | 2.83E-14 |
| Uncharacterized protein OS=Mus musculus OX=10090 GN=Mavs PE=2 SV=1 | Mavs | 0.443 | 2.75E-16 |
| Malonyl-CoA decarboxylase, mitochondrial OS=Mus musculus OX=10090 GN=Mlycd PE=1 SV=1 | Mlycd | 0.436 | 2.75E-16 |
| Ribosomal protein L19 OS=Mus musculus OX=10090 GN=Rpl19 PE=1 SV=1 | Rpl19 | 0.429 | 2.75E-16 |
| Origin recognition complex subunit 1 OS=Mus musculus OX=10090 GN=Orc1 PE=2 SV=1 | Orc1 | 0.415 | 2.75E-16 |
| Protein-serine/threonine phosphatase OS=Mus musculus OX=10090 GN=Ppm1b PE=1 SV=1 | Ppm1b | 0.412 | 2.75E-16 |
| MKIAA1027 protein (Fragment) OS=Mus musculus OX=10090 GN=Tln1 PE=2 SV=4 | Tln1 | 0.318 | 2.75E-16 |
| DAZ-associated protein 1 OS=Mus musculus OX=10090 GN=Dazap1 PE=1 SV=2 | Dazap1 | 0.312 | 2.75E-16 |
| Tr-type G domain-containing protein OS=Mus musculus OX=10090 GN=Gspt1 PE=2 SV=1 | Gspt1 | 0.275 | 2.75E-16 |
| Uncharacterized protein OS=Mus musculus OX=10090 GN=Sec22b PE=2 SV=1 | Sec22b | 0.188 | 2.75E-16 |
| Bile salt export pump OS=Mus musculus OX=10090 GN=Abcb11 PE=1 SV=1 | Abcb11 | 0.171 | 2.75E-16 |
| Serum amyloid A protein OS=Mus musculus OX=10090 GN=Saa2 PE=2 SV=1 | Saa2 | 0.131 | 2.75E-16 |
| Pyruvate kinase PKM OS=Mus musculus OX=10090 GN=Pkm PE=1 SV=4 | Pkm | 0.01 | 2.75E-16 |
| Keratin, type II cuticular Hb4 OS=Mus musculus OX=10090 GN=Krt84 PE=2 SV=2 | Krt84 | 0.01 | 2.75E-16 |
| Fascin OS=Mus musculus OX=10090 GN=Fscn1 PE=1 SV=1 | Fscn1 | 0.01 | 2.75E-16 |
| Guanine nucleotide-binding protein G(I)/G(S)/G(T) subunit beta-1 (Fragment) OS=Mus musculus OX=10090 GN=Gnb1 PE=1 SV=8 | Gnb1 | 0.01 | 2.75E-16 |
| Septin-11 (Fragment) OS=Mus musculus OX=10090 GN=Septin11 PE=1 SV=1 | Septin11 | 0.01 | 2.75E-16 |
| 2-phospho-D-glycerate hydro-lyase OS=Mus musculus OX=10090 GN=Eno3 PE=2 SV=1 | Eno3 | 0.01 | 2.75E-16 |
| EH domain-containing protein 3 OS=Mus musculus OX=10090 GN=Ehd3 PE=1 SV=2 | Ehd3 | 0.01 | 2.75E-16 |
