## Supplementary Table 4 for "A Non-canonical Role for Hepatocyte MLKL in Promoting Mitochondrial Dysfunction and Senescence in the Aging Liver"

**Table S4**: List of mitochondrial proteins identified by targeted proteomics, organized as per functional pathway

| **Proteins** | **Control** | | | | | **MLKL^HepOE^** | | | | |
| --- | --- | --- | --- | --- | --- | --- | --- | --- | --- | --- |
| Protein Abundance (pmol/100 µg) | | | | | | | | | | |
| **Krebs Cycle** | | | | | | | | | | |
| ACO2 | 0.751 | 0.696 | 0.767 | 0.723 | 0.780 | 0.711 | 0.840 | 0.826 | 1.062 | 0.753 |
| CS | 1.911 | 1.744 | 1.871 | 1.738 | 1.927 | 1.748 | 1.892 | 2.201 | 1.901 | 1.470 |
| DLAT | 0.371 | 0.337 | 0.387 | 0.373 | 0.410 | 0.352 | 0.376 | 0.491 | 0.443 | 0.347 |
| DLD | 0.234 | 0.248 | 0.239 | 0.242 | 0.252 | 0.268 | 0.252 | 0.293 | 0.257 | 0.259 |
| DLST | 1.532 | 1.318 | 1.536 | 1.537 | 1.620 | 1.448 | 1.584 | 1.774 | 1.556 | 1.637 |
| FH1 | 0.181 | 0.180 | 0.191 | 0.225 | 0.212 | 0.155 | 0.238 | 0.216 | 0.272 | 0.216 |
| GLUD1 | 2.624 | 3.250 | 3.301 | 3.265 | 2.655 | 3.354 | 3.430 | 3.252 | 2.480 | 3.451 |
| IDH1 | 3.619 | 3.536 | 3.783 | 3.783 | 3.796 | 3.859 | 3.627 | 4.098 | 3.177 | 3.667 |
| IDH2 | 0.637 | 0.739 | 0.728 | 0.700 | 0.676 | 0.859 | 0.849 | 1.079 | 1.330 | 0.869 |
| IDH3A | 0.223 | 0.217 | 0.211 | 0.199 | 0.201 | 0.201 | 0.222 | 0.217 | 0.276 | 0.276 |
| IDH3B | 0.132 | 0.130 | 0.136 | 0.131 | 0.129 | 0.126 | 0.134 | 0.125 | 0.164 | 0.129 |
| MDH1 | 4.820 | 4.399 | 4.658 | 4.400 | 4.660 | 4.489 | 4.942 | 4.673 | 5.128 | 4.696 |
| MDH2 | 2.441 | 2.498 | 2.588 | 2.480 | 2.370 | 2.451 | 2.582 | 2.781 | 3.110 | 2.524 |
| OGDH | 0.226 | 0.220 | 0.224 | 0.212 | 0.181 | 0.212 | 0.251 | 0.281 | 0.278 | 0.243 |
| PDHA1 | 0.718 | 0.743 | 0.759 | 0.719 | 0.809 | 0.785 | 0.699 | 0.983 | 0.853 |  |
| PDHB | 0.296 | 0.350 | 0.366 | 0.335 | 0.326 | 0.362 | 0.370 | 0.425 | 0.366 | 0.364 |
| PDK2 | 0.042 | 0.039 | 0.045 | 0.035 | 0.040 | 0.036 | 0.041 | 0.055 | 0.072 | 0.039 |
| SDHA | 0.882 | 0.968 | 1.043 | 0.930 | 0.920 | 0.965 | 1.137 | 1.059 | 1.155 | 1.006 |
| SDHB | 0.754 | 0.636 | 0.726 | 0.693 | 0.799 | 0.659 | 0.845 | 0.908 | 0.988 | 0.746 |
| SDHC | 0.135 | 0.113 | 0.106 | 0.115 | 0.134 | 0.094 | 0.131 | 0.178 | 0.150 | 0.111 |
| SUCLA2 | 0.718 | 0.761 | 0.803 | 0.802 | 0.748 | 0.794 | 0.840 | 0.880 | 0.883 | 0.810 |
| SUCLG1 | 0.647 | 0.615 | 0.672 | 0.558 | 0.702 | 0.672 | 0.731 | 0.791 | 0.813 | 0.636 |
| VDAC1 | 0.358 | 0.384 | 0.356 | 0.412 | 0.372 | 0.349 | 0.423 | 0.444 | 0.445 | 0.401 |
| VDAC2 | 0.249 | 0.272 | 0.253 | 0.254 | 0.251 | 0.259 | 0.282 | 0.286 | 0.321 | 0.266 |
| VDAC3 | 0.354 | 0.302 | 0.299 | 0.342 | 0.348 | 0.288 | 0.332 | 0.343 | 0.350 | 0.330 |
| **Beta oxidation** | | | | | | | | | | |
| ABCD3 | 0.133 | 0.123 | 0.128 | 0.122 | 0.124 | 0.118 | 0.147 | 0.206 | 0.159 | 0.129 |
| ACAA1A/B | 0.744 | 0.691 | 0.780 | 0.588 | 0.889 | 0.817 | 0.858 | 1.569 | 1.298 | 0.716 |
| ACAA2 | 6.982 | 6.915 | 8.076 | 7.245 | 6.738 | 7.790 | 8.056 | 9.348 | 7.952 | 8.060 |
| ACAD11 | 0.267 | 0.309 | 0.322 | 0.291 | 0.349 | 0.326 | 0.334 | 0.394 | 0.422 | 0.294 |
| ACADL | 0.385 | 0.333 | 0.451 | 0.397 | 0.419 | 0.419 | 0.450 | 0.543 | 0.524 | 0.454 |
| ACADM | 0.565 | 0.544 | 0.585 | 0.549 | 0.641 | 0.615 | 0.712 | 0.813 | 0.912 | 0.619 |
| ACADS | 0.591 | 0.574 | 0.657 | 0.578 | 0.637 | 0.638 | 0.764 | 0.813 | 0.654 | 0.756 |
| ACADVL | 0.644 | 0.628 | 0.688 | 0.646 | 0.712 | 0.687 | 0.676 | 0.824 | 0.853 | 0.667 |
| ACOT13 | 0.338 | 0.331 | 0.289 | 0.316 | 0.324 | 0.289 | 0.307 | 0.386 | 0.488 | 0.295 |
| ACOX1 | 0.879 | 0.768 | 0.731 | 0.878 | 0.998 | 0.863 | 1.167 | 1.041 | 1.030 | 0.777 |
| ACOX2 | 0.120 | 0.111 | 0.131 | 0.135 | 0.125 | 0.123 | 0.132 | 0.139 | 0.142 | 0.140 |
| ACSL1 | 2.178 | 2.310 | 2.657 | 2.219 | 2.338 | 2.566 | 2.510 | 2.702 | 2.360 | 2.497 |
| CPT1A | 0.145 | 0.110 | 0.152 | 0.123 | 0.143 | 0.119 | 0.163 | 0.153 | 0.174 | 0.134 |
| CPT2 | 0.335 | 0.360 | 0.411 | 0.356 | 0.360 | 0.390 | 0.386 | 0.444 | 0.461 | 0.384 |
| CROT | 0.050 | 0.049 | 0.063 | 0.037 | 0.067 | 0.050 | 0.066 | 0.066 | 0.065 | 0.065 |
| DECR1 | 1.587 | 1.672 | 1.838 | 1.639 | 1.883 | 1.832 | 1.878 | 2.293 | 1.908 | 1.644 |
| DECR2 | 0.262 | 0.277 | 0.266 | 0.275 | 0.229 | 0.275 | 0.305 | 0.393 | 0.327 | 0.297 |
| ECH1 | 0.334 | 0.387 | 0.416 | 0.349 | 0.421 | 0.451 | 0.485 | 0.553 | 0.590 | 0.450 |
| ECHS1 | 0.508 | 0.446 | 0.477 | 0.513 | 0.550 | 0.486 | 0.554 | 0.545 | 0.613 | 0.499 |
| ECI1 | 0.439 | 0.426 | 0.497 | 0.424 | 0.452 | 0.467 | 0.511 | 0.587 | 0.553 | 0.455 |
| EHHADH | 0.534 | 0.442 | 0.523 | 0.427 | 0.626 | 0.536 | 0.702 | 0.883 | 1.209 | 0.531 |
| ETFA | 1.383 | 1.352 | 1.468 | 1.315 | 1.476 | 1.323 | 1.597 | 1.793 | 1.833 | 1.457 |
| ETFB | 1.944 | 2.027 | 2.023 | 2.121 | 2.046 | 1.989 | 2.117 | 2.249 | 2.414 | 2.007 |
| ETFDH | 0.706 | 0.719 | 0.832 | 0.688 | 0.702 | 0.791 | 0.874 | 0.836 | 0.818 | 0.767 |
| FABP4 | 0.113 | 0.144 | 0.097 | 0.136 | 0.115 | 0.106 | 0.135 | 0.124 | 0.685 | 0.119 |
| HADH | 4.251 | 3.820 | 4.207 | 3.825 | 4.594 | 4.033 | 4.703 | 5.263 | 5.257 | 4.229 |
| HADHA | 0.820 | 0.860 | 0.992 | 0.835 | 0.941 | 0.917 | 0.989 | 1.126 | 1.072 | 0.907 |
| HADHB | 1.902 | 2.140 | 2.303 | 2.193 | 2.480 | 2.370 | 2.469 | 2.827 | 2.609 | 2.189 |
| HSD17B4 | 0.790 | 0.872 | 0.900 | 0.890 | 0.950 | 0.885 | 1.058 | 1.106 | 0.970 | 0.895 |
| PECR | 1.553 | 1.532 | 1.534 | 1.686 | 1.409 | 1.622 | 1.632 | 1.683 | 1.750 | 1.624 |
| SLC25A20 | 0.431 | 0.464 | 0.513 | 0.478 | 0.518 | 0.502 | 0.548 | 0.602 | 0.497 | 0.503 |
| **Electron Transport Chain Complex I** | | | | | | | | | | |
| ND1 | 0.241 | 0.215 | 0.229 | 0.295 | 0.249 | 0.241 | 0.275 | 0.278 | 0.271 | 0.245 |
| ND3 | 0.058 | 0.035 | 0.042 | 0.039 | 0.045 | 0.028 | 0.059 | 0.079 | 0.061 | 0.040 |
| ND4 | 0.058 | 0.041 | 0.051 | 0.053 | 0.055 | 0.040 | 0.047 | 0.062 | 0.057 | 0.038 |
| NDUFA10 | 0.271 | 0.293 | 0.256 | 0.324 | 0.274 | 0.272 | 0.311 | 0.324 | 0.316 | 0.285 |
| NDUFA11 | 0.052 |  | 0.037 | 0.060 | 0.056 | 0.041 | 0.058 | 0.056 | 0.049 | 0.044 |
| NDUFA12 | 0.134 | 0.134 | 0.122 | 0.157 | 0.128 | 0.143 | 0.128 | 0.154 | 0.160 | 0.121 |
| NDUFA13 | 0.253 | 0.266 | 0.204 | 0.281 | 0.261 | 0.255 | 0.277 | 0.297 | 0.323 | 0.236 |
| NDUFA2 | 0.121 | 0.111 | 0.100 | 0.115 | 0.113 | 0.095 | 0.118 | 0.119 | 0.138 | 0.103 |
| NDUFA4 | 0.204 | 0.157 | 0.134 | 0.214 | 0.156 | 0.146 | 0.161 | 0.188 | 0.254 | 0.151 |
| NDUFA5 | 0.109 | 0.103 | 0.090 | 0.111 | 0.106 | 0.101 | 0.110 | 0.115 | 0.123 | 0.103 |
| NDUFA6 | 0.156 | 0.150 | 0.136 | 0.213 | 0.144 | 0.138 | 0.153 | 0.187 | 0.234 | 0.132 |
| NDUFA7 | 0.051 | 0.045 | 0.039 | 0.058 | 0.039 | 0.041 | 0.040 | 0.048 | 0.057 | 0.041 |
| NDUFA8 | 0.114 | 0.123 | 0.090 | 0.121 | 0.104 | 0.104 | 0.132 | 0.136 | 0.151 | 0.117 |
| NDUFA9 | 0.144 | 0.125 | 0.123 | 0.127 | 0.156 | 0.123 | 0.160 | 0.170 | 0.160 | 0.121 |
| NDUFB10 | 0.254 | 0.211 | 0.234 | 0.217 | 0.262 | 0.209 | 0.230 | 0.288 | 0.299 | 0.221 |
| NDUFB11 | 0.129 | 0.125 | 0.121 | 0.165 | 0.111 | 0.133 | 0.139 | 0.130 | 0.159 | 0.118 |
| NDUFB5 | 0.106 | 0.098 | 0.104 | 0.131 | 0.100 | 0.091 | 0.097 | 0.103 | 0.114 | 0.108 |
| NDUFB6 | 0.084 | 0.068 | 0.068 | 0.108 | 0.089 | 0.079 | 0.085 | 0.107 | 0.109 | 0.081 |
| NDUFB7 | 0.111 | 0.113 | 0.095 | 0.134 | 0.117 | 0.100 | 0.145 | 0.169 | 0.159 | 0.121 |
| NDUFB9 | 0.118 | 0.121 | 0.101 | 0.111 | 0.140 | 0.122 | 0.120 | 0.138 | 0.142 | 0.115 |
| NDUFC2 | 0.222 | 0.178 | 0.180 | 0.237 | 0.224 | 0.193 | 0.242 | 0.242 | 0.272 | 0.208 |
| NDUFS1 | 0.492 | 0.468 | 0.461 | 0.531 | 0.538 | 0.532 | 0.549 | 0.582 | 0.642 | 0.495 |
| NDUFS2 | 0.252 | 0.197 | 0.232 | 0.250 | 0.229 | 0.236 | 0.283 | 0.271 | 0.254 | 0.249 |
| NDUFS3 | 0.336 | 0.277 | 0.267 | 0.330 | 0.348 | 0.295 | 0.354 | 0.348 | 0.375 | 0.292 |
| NDUFS4 | 0.189 | 0.150 | 0.156 | 0.197 | 0.185 | 0.163 | 0.198 | 0.248 | 0.240 | 0.182 |
| NDUFS5 | 0.249 | 0.223 | 0.213 | 0.226 | 0.232 | 0.218 | 0.238 | 0.225 | 0.287 | 0.213 |
| NDUFS7 | 0.147 | 0.114 | 0.120 | 0.127 | 0.129 | 0.123 | 0.134 | 0.142 | 0.162 | 0.110 |
| NDUFS8 | 0.273 | 0.221 | 0.211 | 0.262 | 0.280 | 0.265 | 0.268 | 0.281 | 0.333 | 0.257 |
| NDUFV1 | 0.435 | 0.396 | 0.410 | 0.444 | 0.448 | 0.416 | 0.451 | 0.474 | 0.526 | 0.426 |
| NDUFV2 | 0.207 | 0.170 | 0.173 | 0.247 | 0.232 | 0.197 | 0.209 | 0.225 | 0.248 | 0.199 |
| **Electron Transport Chain Complex 3a4a5** | | | | | | | | | | |
|  | 5c 3 | 5c 4 | 5c 5 | 5c 1 | 5c 2 | 5T 10 | 5T 9 | 5T 6 | 5T 7 | 5T 8 |
| ACTIN | 4.954 | 4.722 | 4.559 | 4.891 | 5.361 | 4.690 | 5.254 | 5.884 | 5.464 | 4.156 |
| ATP5A1 | 3.505 | 2.634 | 3.425 | 2.859 | 3.710 | 3.118 | 3.664 | 3.767 | 3.874 | 3.102 |
| ATP5B | 3.896 | 3.106 | 3.546 | 4.020 | 3.826 | 3.298 | 4.008 | 3.942 | 3.712 | 3.424 |
| ATP5C1 | 0.982 | 1.020 | 1.100 | 1.051 | 1.011 | 1.042 | 1.069 | 1.150 | 1.082 | 0.896 |
| ATP5D | 0.731 | 0.647 | 0.646 | 0.734 | 0.664 | 0.604 | 0.698 | 0.751 | 0.746 | 0.621 |
| ATP5E | 0.244 | 0.266 | 0.212 | 0.302 | 0.195 | 0.213 | 0.297 | 0.341 | 0.349 | 0.233 |
| ATP5J2 | 1.422 | 1.305 | 1.328 | 1.756 | 1.392 | 1.246 | 1.472 | 1.699 | 1.784 | 1.367 |
| ATP5K | 0.093 | 0.091 | 0.074 | 0.095 | 0.084 | 0.074 | 0.098 | 0.087 | 0.073 | 0.075 |
| ATP5L | 0.159 | 0.164 | 0.130 | 0.185 | 0.126 | 0.113 | 0.195 | 0.166 | 0.174 | 0.143 |
| ATP5MD | 0.194 | 0.190 | 0.135 | 0.165 | 0.175 | 0.139 | 0.185 | 0.187 | 0.186 | 0.149 |
| ATP5O | 1.206 | 1.184 | 1.182 | 1.554 | 1.357 | 1.238 | 1.309 | 1.440 | 1.419 | 1.181 |
| ATP5PB | 1.221 | 1.083 | 1.138 | 1.126 | 1.249 | 1.094 | 1.145 | 1.281 | 1.262 | 1.016 |
| ATP5PD | 0.510 | 0.493 | 0.477 | 0.630 | 0.526 | 0.483 | 0.483 | 0.557 | 0.523 | 0.468 |
| COX1 MTCO1 | 0.141 | 0.111 | 0.120 | 0.123 | 0.145 | 0.115 | 0.151 | 0.148 | 0.134 | 0.132 |
| COX2 MTCO2 | 3.108 | 5.069 | 2.988 | 2.750 | 3.372 | 2.825 | 3.385 | 3.484 | 4.019 | 3.056 |
| COX3 MTCO3 | 0.083 | 0.062 | 0.086 | 0.091 | 0.090 | 0.081 | 0.090 | 0.091 | 0.098 | 0.073 |
| COX4I1 | 1.210 | 0.960 | 0.881 | 1.163 | 1.167 | 0.931 | 1.210 | 1.165 | 1.338 | 1.042 |
| COX5A | 1.399 | 1.344 | 1.113 | 1.538 | 1.224 | 1.138 | 1.261 | 1.366 | 1.711 | 1.241 |
| COX5B | 0.178 | 0.147 | 0.146 | 0.171 | 0.194 | 0.160 | 0.214 | 0.198 | 0.157 | 0.176 |
| COX6B1 | 0.176 | 0.136 | 0.114 | 0.136 | 0.144 | 0.117 | 0.172 | 0.198 | 0.212 | 0.135 |
| COX6C | 0.506 | 0.515 | 0.431 | 0.564 | 0.437 | 0.377 | 0.513 | 0.481 | 0.603 | 0.456 |
| CYC1 | 0.205 | 0.171 | 0.178 | 0.129 | 0.219 | 0.139 | 0.202 | 0.214 | 0.207 | 0.164 |
| CYTB | 0.115 |  | 0.104 | 0.114 | 0.112 | 0.101 | 0.125 | 0.132 | 0.118 | 0.109 |
| UQCR10 | 0.335 | 0.385 | 0.265 | 0.519 | 0.302 | 0.270 | 0.354 | 0.437 | 0.450 | 0.316 |
| UQCR11 | 0.070 | 0.056 | 0.051 | 0.081 | 0.063 | 0.051 | 0.068 | 0.079 | 0.094 | 0.058 |
| UQCRC1 | 1.044 | 0.995 | 1.169 | 1.047 | 1.142 | 1.151 | 1.133 | 1.236 | 1.304 | 1.054 |
| UQCRC2 | 0.536 | 0.433 | 0.444 | 0.426 | 0.514 | 0.474 | 0.501 | 0.489 | 0.552 | 0.440 |
| **Branched Chain Amino Acid catabolism** | | | | | | | | | | |
| ACAD8 | 0.262 | 0.248 | 0.257 | 0.277 | 0.257 | 0.277 | 0.249 | 0.300 | 0.278 | 0.232 |
| ACADSB | 0.282 | 0.269 | 0.284 | 0.278 | 0.297 | 0.268 | 0.306 | 0.324 | 0.313 | 0.288 |
| ACAT1 | 1.438 | 1.352 | 1.600 | 1.506 | 1.369 | 1.556 | 1.444 | 1.760 | 1.636 | 1.252 |
| ALDH6A1 | 1.431 | 1.261 | 1.419 | 1.653 | 1.433 | 1.505 | 1.479 | 1.520 | 1.606 | 1.346 |
| AUH | 0.156 | 0.145 | 0.149 | 0.169 | 0.164 | 0.141 | 0.156 | 0.175 | 0.175 | 0.136 |
| BCKDHA | 0.115 | 0.086 | 0.104 | 0.161 | 0.106 | 0.104 | 0.120 | 0.115 | 0.137 | 0.100 |
| BCKDHB | 0.070 | 0.064 | 0.067 | 0.076 | 0.065 | 0.070 | 0.086 | 0.075 | 0.083 | 0.070 |
| DBT | 0.633 | 0.567 | 0.627 | 0.569 | 0.559 | 0.647 | 0.706 | 0.667 | 0.848 | 0.614 |
| DLD | 0.230 | 0.209 | 0.212 | 0.251 | 0.209 | 0.230 | 0.227 | 0.230 | 0.222 | 0.201 |
| HIBADH | 0.186 | 0.158 | 0.185 | 0.207 | 0.192 | 0.208 | 0.202 | 0.202 | 0.248 | 0.182 |
| HIBCH | 0.412 | 0.355 | 0.405 | 0.429 | 0.366 | 0.434 | 0.404 | 0.418 | 0.412 | 0.373 |
| HMGCL | 0.350 | 0.304 | 0.322 | 0.405 | 0.367 | 0.313 | 0.338 | 0.438 | 0.340 | 0.323 |
| HSD17B10 | 1.416 | 1.237 | 1.086 | 1.598 | 1.343 | 1.168 | 1.443 | 1.323 | 1.560 | 1.302 |
| IVD | 0.839 | 0.774 | 0.852 | 0.751 | 0.870 | 0.931 | 0.916 | 0.966 | 1.098 | 0.857 |
| MCCC1 | 0.108 | 0.089 | 0.106 | 0.109 | 0.122 | 0.112 | 0.101 | 0.105 | 0.114 | 0.097 |
| MCCC2 | 0.185 | 0.162 | 0.189 | 0.207 | 0.186 | 0.193 | 0.187 | 0.190 | 0.232 | 0.171 |
| **Glycolysis and Gluconeogenesis** | | | | | | | | | | |
| ALDOA | 0.196 | 0.183 | 0.195 | 0.156 | 0.210 | 0.175 | 0.207 | 0.221 | 0.455 | 0.187 |
| ALDOB | 20.710 | 18.196 | 17.655 | 17.099 | 21.841 | 17.244 | 19.001 | 20.336 | 17.624 | 16.989 |
| ENO1 | 2.926 | 2.266 | 2.056 | 2.931 | 3.036 | 1.967 | 2.394 | 3.115 | 2.077 | 2.072 |
| FBP1 | 1.744 | 1.222 | 1.298 | 1.686 | 1.775 | 1.321 | 1.571 | 1.744 | 1.693 | 1.374 |
| GAPDH | 7.940 | 7.111 | 6.455 | 7.975 | 8.040 | 6.552 | 7.927 | 7.766 | 7.096 | 6.973 |
| GOT1 | 0.615 | 0.458 | 0.417 | 0.611 | 0.527 | 0.495 | 0.624 | 0.529 | 1.444 | 0.526 |
| GOT2 | 2.644 | 2.172 | 2.303 | 3.388 | 2.619 | 2.423 | 2.413 | 2.501 | 2.478 | 2.190 |
| GPI | 0.136 | 0.106 | 0.093 | 0.121 | 0.149 | 0.106 | 0.102 | 0.114 | 0.093 | 0.102 |
| LDHA | 9.685 | 9.098 | 7.954 | 8.612 | 9.261 | 8.798 | 9.730 | 8.759 | 7.916 | 8.812 |
| MPC1 | 0.438 | 0.431 | 0.441 | 0.405 | 0.375 | 0.439 | 0.440 | 0.517 | 0.627 | 0.398 |
| MPC2 | 0.338 | 0.328 | 0.372 | 0.383 | 0.295 | 0.386 | 0.378 | 0.406 | 0.413 | 0.344 |
| PC | 1.456 | 1.287 | 1.339 | 2.044 | 1.549 | 1.477 | 1.404 | 1.547 | 1.421 | 1.366 |
| PCK1 | 0.298 | 0.429 | 0.338 | 0.466 | 0.429 | 0.345 | 0.363 | 0.224 | 0.443 | 0.330 |
| PGAM1 | 0.503 | 0.443 | 0.448 | 0.498 | 0.558 | 0.427 | 0.472 | 0.535 | 0.490 | 0.445 |
| PGK1 | 0.974 | 0.860 | 0.957 | 1.049 | 0.956 | 0.883 | 0.985 | 1.023 | 0.999 | 0.935 |
| PKLR | 2.392 | 2.112 | 1.775 | 1.858 | 2.386 | 1.553 | 1.640 | 2.472 | 0.611 | 1.530 |
| PKM | 0.478 | 0.361 | 0.390 | 0.399 | 0.489 | 0.346 | 0.412 | 0.465 | 0.549 | 0.387 |
| TALDO1 | 0.430 | 0.418 | 0.398 | 0.446 | 0.444 | 0.401 | 0.457 | 0.504 | 0.438 | 0.387 |
| TKT | 2.000 | 2.214 | 2.032 | 1.715 | 2.193 | 1.916 | 2.014 | 2.568 | 1.550 | 1.765 |
| TPI1 | 0.568 | 0.552 | 0.453 | 0.699 | 0.610 | 0.542 | 0.536 | 0.582 | 0.564 | 0.504 |
| **Antioxidants** | | | | | | | | | | |
| AKR1B1 | 0.118 | 0.098 | 0.090 | 0.091 | 0.095 | 0.104 | 0.103 | 0.091 | 0.127 | 0.084 |
| ALDH2 | 2.262 | 1.961 | 2.204 | 2.364 | 2.453 | 2.302 | 2.149 | 2.609 | 2.352 | 2.029 |
| CAT | 1.916 | 1.449 | 1.662 | 1.569 | 1.961 | 1.700 | 1.861 | 2.080 | 1.868 | 1.536 |
| GPX1 | 1.189 | 0.944 | 1.058 | 1.364 | 1.090 | 1.051 | 1.111 | 1.019 | 1.037 | 1.049 |
| GPX4 | 0.181 | 0.204 | 0.164 | 0.219 | 0.176 | 0.182 | 0.207 | 0.249 | 0.220 | 0.186 |
| GSR | 0.154 | 0.128 | 0.126 | 0.146 | 0.145 | 0.146 | 0.136 | 0.172 | 0.146 | 0.136 |
| GSTA3 | 16.287 | 15.762 | 13.191 | 15.340 | 15.747 | 14.224 | 15.969 | 18.740 | 16.091 | 14.262 |
| GSTM1 | 17.972 | 21.046 | 20.140 | 18.455 | 21.800 | 22.733 | 19.547 | 30.933 | 26.542 | 17.893 |
| GSTP1 | 9.596 | 7.506 | 8.397 | 12.338 | 9.939 | 8.667 | 7.695 | 8.903 | 3.518 | 8.541 |
| HSP90AA1 | 0.309 | 0.286 | 0.315 | 0.317 | 0.328 | 0.280 | 0.327 | 0.303 | 0.226 | 0.294 |
| HSP90AB1 | 2.578 | 2.346 | 2.503 | 2.904 | 2.730 | 2.521 | 2.692 | 2.502 | 2.184 | 2.323 |
| HSP90B1 | 1.249 | 1.155 | 1.187 | 1.344 | 1.239 | 1.202 | 1.317 | 1.067 | 0.960 | 1.145 |
| HSPA1/8 | 2.706 | 2.347 | 2.444 | 3.328 | 2.893 | 2.552 | 2.725 | 2.542 | 2.016 | 2.460 |
| HSPA1A/B | 0.079 | 0.088 | 0.085 | 0.092 | 0.089 | 0.071 | 0.111 | 0.077 | 0.073 | 0.097 |
| HSPA4 | 0.119 | 0.105 | 0.120 | 0.168 | 0.128 | 0.118 | 0.105 | 0.115 | 0.098 | 0.107 |
| HSPA5 | 0.817 | 0.629 | 0.630 | 1.214 | 0.863 | 0.676 | 0.720 | 0.561 | 0.472 | 0.711 |
| HSPA9 | 0.870 | 0.779 | 0.852 | 1.151 | 0.948 | 0.892 | 0.880 | 0.911 | 0.992 | 0.814 |
| HSPD1 | 1.906 | 1.697 | 1.865 | 1.882 | 1.962 | 1.945 | 1.980 | 1.938 | 1.882 | 1.751 |
| HSPE1 | 0.705 | 0.627 | 0.487 | 0.646 | 0.569 | 0.467 | 0.629 | 0.655 | 0.790 | 0.529 |
| LONP1 | 0.251 | 0.246 | 0.248 | 0.311 | 0.261 | 0.261 | 0.247 | 0.287 | 0.276 | 0.223 |
| LONP2 | 0.081 | 0.064 | 0.078 | 0.110 | 0.086 | 0.069 | 0.080 | 0.083 | 0.064 | 0.079 |
| MSRA | 0.946 | 0.840 | 0.790 | 0.887 | 0.956 | 0.897 | 1.041 | 1.039 | 0.853 | 0.935 |
| PHB1 | 1.115 | 0.866 | 1.061 | 1.383 | 1.139 | 0.874 | 1.148 | 1.202 | 1.022 | 0.957 |
| PHB2 | 0.676 | 0.574 | 0.551 | 0.633 | 0.695 | 0.568 | 0.651 | 0.686 | 0.670 | 0.534 |
| PRDX1 | 8.503 | 7.579 | 7.225 | 7.632 | 8.256 | 7.793 | 7.950 | 9.072 | 8.002 | 7.415 |
| PRDX2 | 1.020 | 0.852 | 0.861 | 0.822 | 0.996 | 0.889 | 0.922 | 1.009 | 1.112 | 0.870 |
| PRDX3 | 0.610 | 0.455 | 0.479 | 0.520 | 0.628 | 0.502 | 0.553 | 0.606 | 0.572 | 0.503 |
| PRDX4 | 0.165 | 0.154 | 0.122 | 0.172 | 0.137 | 0.156 | 0.164 | 0.158 | 0.133 | 0.142 |
| PRDX5 | 1.627 | 1.287 | 1.303 | 2.029 | 1.515 | 1.316 | 1.502 | 1.477 | 1.324 | 1.440 |
| PRDX6 | 1.750 | 1.437 | 1.399 | 2.057 | 1.714 | 1.521 | 1.654 | 1.717 | 1.513 | 1.528 |
| SOD1 | 3.422 | 2.781 | 3.084 | 2.670 | 3.219 | 3.277 | 3.075 | 3.739 | 2.968 | 2.931 |
| SOD2 | 1.430 | 1.389 | 1.358 | 1.656 | 1.462 | 1.439 | 1.373 | 1.495 | 1.386 | 1.332 |
| TXN1 | 2.221 | 1.842 | 1.701 | 2.619 | 1.949 | 1.765 | 1.856 | 2.268 | 1.695 | 1.762 |
| TXNRD1 | 0.199 | 0.151 | 0.166 | 0.248 | 0.203 | 0.160 | 0.187 | 0.187 | 0.156 | 0.176 |
| **Other Mitochondrial Proteins** | | | | | | | | | | |
| ATP6V1B2 | 0.120 | 0.116 | 0.128 | 0.142 | 0.126 | 0.129 | 0.140 | 0.134 | 0.165 | 0.118 |
| CALR | 1.883 | 1.586 | 1.578 | 2.762 | 2.616 | 1.060 | 1.930 | 1.736 | 1.457 | 1.880 |
| DDAH1 | 0.191 | 0.259 | 0.268 | 0.218 | 0.272 | 0.256 | 0.284 | 0.311 | 0.230 | 0.248 |
| GNB1/2 | 0.377 | 0.521 | 0.422 | 0.499 | 0.430 | 0.459 | 0.444 | 0.486 | 0.485 | 0.401 |
| POR | 0.933 | 0.889 | 0.970 | 0.942 | 0.955 | 1.201 | 1.164 | 1.160 | 1.782 | 0.955 |
| SFXN1 | 0.194 | 0.168 | 0.189 | 0.190 | 0.197 | 0.173 | 0.201 | 0.185 | 0.225 | 0.189 |
| SLC25A11 | 0.389 | 0.399 | 0.361 | 0.361 | 0.403 | 0.379 | 0.415 | 0.464 | 0.498 | 0.315 |
| SLC25A3 | 1.067 | 1.410 | 1.186 | 0.946 | 1.567 | 1.131 | 1.169 | 1.232 | 1.394 | 0.841 |
| SLC25A4 | 2.707 | 2.246 | 2.552 | 2.934 | 2.324 | 2.666 | 2.961 | 3.114 | 3.143 | 2.606 |
| SLC25A5 | 3.235 | 2.986 | 2.977 | 3.470 | 3.459 | 3.136 | 3.500 | 3.770 | 3.655 | 3.056 |
| SLC25A-Multiple | 8.931 | 8.733 | 9.081 | 9.304 | 9.021 | 8.483 | 9.058 | 9.961 | 10.497 | 8.433 |
| SLC9A3R1 | 0.228 | 0.180 | 0.194 | 0.240 | 0.230 | 0.194 | 0.236 | 0.223 | 0.189 | 0.207 |
