## Supplementary Table 5 for "A Non-canonical Role for Hepatocyte MLKL in Promoting Mitochondrial Dysfunction and Senescence in the Aging Liver"

| **Downregulated lipid species**  **Table S5:** List of differentially regulated lipid species in lipidomic analysis | | | | |
| --- | --- | --- | --- | --- |
| **Lipid ID** | **Lipid Class** | **FattyAcid** | **log2(FC)** | **log10 (p value)** |
| PEt_1128n | PEt | (36:0e) | -2.91253716 | 2.634644645 |
| DG_262p | DG | (16:1e) | -2.80735492 | 0.988940517 |
| LPE_344n | LPE | (22:4) | -2.45285896 | 2.333376926 |
| PE_753n | PE | (16:0_18:2) | -1.94753258 | 2.259636761 |
| PE_3095p | PE | (16:0e) | -1.9068906 | 0.993061628 |
| PC_2388p | PC | (8:1e_10:1) | -1.81557543 | 2.420110134 |
| LdMePE_379n | LdMePE | (16:0) | -1.68314289 | 0.501251711 |
| MePC_2233p | MePC | (37:4e) | -1.65711229 | 1.287339851 |
| PE_710n | PE | (20:2e) | -1.61325155 | 1.248581419 |
| LPC_1000p | LPC | (18:2) | -1.61143471 | 2.665049656 |
| ChE_195p | ChE | (20:4) | -1.5849625 | 1.830768427 |
| MePC_2272p | MePC | (39:4) | -1.56187889 | 2.450627692 |
| PC_2554p | PC | (20:1_14:0) | -1.56187889 | 1.477133465 |
| DG_675p | DG | (19:0_18:2) | -1.53533173 | 2.644996111 |
| Cer_21n | Cer | (d40:1) | -1.53373718 | 1.221083757 |
| PE_3621p | PE | (20:4_22:6) | -1.52356196 | 2.254267406 |
| PE_3407p | PE | (16:0e_21:1) | -1.5187288 | 4.020422757 |
| MG_1730p | MG | (29:1) | -1.51727569 | 0.915690563 |
| PC_2744p | PC | (36:2) | -1.45943162 | 1.280222073 |
| PA_506n | PA | (8:0e_10:0) | -1.45490196 | 1.059100768 |
| PE_950n | PE | (18:0e_20:4) | -1.44745898 | 1.412670233 |
| PE_939n | PE | (18:0_20:4) | -1.43538614 | 1.474650601 |
| DG_475p | DG | (17:1_16:1) | -1.43295941 | 1.129588993 |
| PE_685n | PE | (18:1e) | -1.35693454 | 1.541742036 |
| PE_830n | PE | (36:2) | -1.3360492 | 0.929928487 |
| PC_2384p | PC | (8:1e_10:1) | -1.32192809 | 2.510672886 |
| MePC_1996p | MePC | (33:1) | -1.31577587 | 1.579708951 |
| PC_2569p | PC | (20:0_14:1) | -1.25457283 | 1.526480587 |
| LdMePE_377n | LdMePE | (16:0) | -1.24792751 | 1.561257113 |
| LdMePE_446n | LdMePE | (20:4) | -1.23562825 | 0.763388212 |
| PC_3049p | PC | (18:4_22:4) | -1.23266076 | 2.816607788 |
| MePC_2259p | MePC | (38:9) | -1.22616911 | 1.201686538 |
| PC_2686p | PC | (34:2e) | -1.2107671 | 1.363757171 |
| TG_4712p | TG | (20:3_20:5_22:6) | -1.20903404 | 1.739802403 |
| PC_573n | PC | (18:0_16:0) | -1.20163386 | 1.976754087 |
| PE_3463p | PE | (18:0_20:4) | -1.19714647 | 0.743373361 |
| SM_3814p | SM | (d36:1) | -1.13404797 | 2.965365937 |
| PC_2935p | PC | (18:3_20:3) | -1.13358315 | 1.019090011 |
| LPE_340n | LPE | (20:4) | -1.12151782 | 1.510455906 |
| Cer_113p | Cer | (d18:1_24:1) | -1.11973924 | 1.789807522 |
| PE_3599p | PE | (40:7) | -1.10825289 | 0.943522679 |
| TG_4225p | TG | (15:0_14:0_16:1) | -1.08615664 | 2.159630319 |
| PE_3282p | PE | (22:0_12:2) | -1.07885542 | 1.594513786 |
| LPE_280n | LPE | (17:0) | -1.06413034 | 1.19567151 |
| MG_1733p | MG | (29:1) | -1.06342538 | 0.915584651 |
| PE_3286p | PE | (16:1e_18:1) | -1.00580562 | 1.208568828 |
| DG_311p | DG | (19:2e) | -0.98859524 | 0.822596719 |
| Cer_65p | Cer | (d18:0_16:0) | -0.95269429 | 1.714795646 |
| MePC_2243p | MePC | (37:6) | -0.9248125 | 1.351300172 |
| dMePE_1897n | dMePE | (16:0_22:6) | -0.92386893 | 1.714337725 |
| PC_3045p | PC | (40:8) | -0.88613204 | 0.935463527 |
| PE_3100p | PE | (16:1e) | -0.85913746 | 1.349066581 |
| PE_1067n | PE | (18:1_22:6) | -0.84582907 | 1.378988373 |
| SM_1389n | SM | (d34:1) | -0.83927969 | 2.206309382 |
| LdMePE_453n | LdMePE | (22:4) | -0.83561318 | 1.771463169 |
| DG_748p | DG | (54:4) | -0.83471539 | 1.17083938 |
| dMePE_1590n | dMePE | (16:0_18:1) | -0.8326056 | 0.841170297 |
| LdMePE_443n | LdMePE | (20:4) | -0.82086685 | 1.576108326 |
| dMePE_1978n | dMePE | (20:4_22:6) | -0.80851886 | 2.182697755 |
| LPA_222n | LPA | (20:3) | -0.80145432 | 1.092645305 |
| dMePE_1698n | dMePE | (17:1_18:2) | -0.78622071 | 1.668503963 |
| PE_764n | PE | (16:1_18:2) | -0.78297076 | 1.387065998 |
| MG_1776p | MG | (29:1) | -0.78068697 | 1.171795264 |
| DG_233p | DG | (16:0e) | -0.77536911 | 0.735875293 |
| MG_1199p | MG | (16:0) | -0.77536911 | 0.735875293 |
| PE_3372p | PE | (16:0p_20:4) | -0.77230797 | 1.188977913 |
| StE_3920p | StE | (16:0) | -0.76733924 | 3.827204131 |
| LPE_254n | LPE | (16:0) | -0.7635598 | 1.088000897 |
| dMePE_1778n | dMePE | (18:1_18:3) | -0.75032398 | 2.185257877 |
| PC_2299p | PC | (15:0) | -0.74908188 | 1.559025885 |
| PE_3487p | PE | (38:6) | -0.73887518 | 1.256189349 |
| PIP_1328n | PIP | (43:3e) | -0.73696559 | 1.176891536 |
| PE_3614p | PE | (41:5e) | -0.73426644 | 2.482140097 |
| dMePE_1974n | dMePE | (18:2_22:6) | -0.71848617 | 1.065486063 |
| TG_4044p | TG | (10:0_12:2_12:2) | -0.71271805 | 1.511137368 |
| LPC_1081p | LPC | (20:4) | -0.70744651 | 1.277613988 |
| LPC_1095p | LPC | (20:5) | -0.70209513 | 1.149506601 |
| PC_2819p | PC | (22:0_14:4) | -0.70191865 | 1.459273549 |
| DLCL_142n | DLCL | (40:8) | -0.6918777 | 1.143717329 |
| TG_4105p | TG | (12:1e_6:0_18:3) | -0.68846708 | 1.701295748 |
| LPE_1142p | LPE | (16:0) | -0.67442413 | 2.239102174 |
| dMePE_1775n | dMePE | (16:0_20:4) | -0.65298092 | 0.95707941 |
| PC_3044p | PC | (40:8) | -0.63911827 | 1.002321928 |
| PE_768n | PE | (16:0_18:3) | -0.63584367 | 2.215473296 |
| PI_1275n | PI | (18:3_18:2) | -0.62583478 | 0.997603458 |
| PE_3603p | PE | (40:8) | -0.60741735 | 2.464270378 |
| PA_562n | PA | (20:4_22:6) | -0.58862882 | 1.52989753 |
| TG_4256p | TG | (46:2) | -0.57994443 | 2.57447416 |
| LPC_1078p | LPC | (20:4) | -0.56910855 | 1.885260294 |
| LPC_1128p | LPC | (22:6) | -0.55908387 | 3.033171985 |
| TG_4705p | TG | (20:1_18:1_22:6) | -0.55581616 | 1.685019348 |
| LPC_244n | LPC | (20:3) | -0.54949782 | 1.356721741 |
| LPC_246n | LPC | (20:4) | -0.54167094 | 0.991632342 |
| LPC_1132p | LPC | (22:6) | -0.51164582 | 2.218687934 |
| cPA_1438n | cPA | (16:0) | -0.50951027 | 1.656362486 |
| SPHP_1430n | SPHP | (m22:1) | -0.50753753 | 0.966835573 |
| CerP_101n | CerP | (d44:3) | -0.50182127 | 0.972452958 |
| PE_3181p | PE | (10:1e_10:2) | -0.50165985 | 0.938928709 |
| MePC_1906p | MePC | (21:3e) | -0.48351724 | 1.484784815 |
| PIP_1327n | PIP | (37:4e) | -0.46496272 | 1.49745207 |
| PC_616n | PC | (17:0_20:4) | -0.46406307 | 2.796222525 |
| PC_2486p | PC | (10:1e_14:4) | -0.45943162 | 1.227866285 |
| Hex2Cer_856p | Hex2Cer | (d45:0+O) | -0.45779807 | 2.765715928 |
| Hex2Cer_865p | Hex2Cer | (t45:0) | -0.45779807 | 2.765715928 |
| LPC_1127p | LPC | (22:6) | -0.45655677 | 1.868572247 |
| MePC_2273p | MePC | (39:4e) | -0.45391039 | 2.337212301 |
| PC_2882p | PC | (28:0_10:4) | -0.44332654 | 1.302724218 |
| PC_3034p | PC | (40:7) | -0.44332654 | 1.302724218 |
| dMePE_1841n | dMePE | (18:0_20:2) | -0.43785821 | 0.933544845 |
| PE_3481p | PE | (18:0_20:5) | -0.41932486 | 3.176935332 |
| CmE_206p | CmE | (12:0) | -0.40690962 | 2.111682505 |
| PC_659n | PC | (18:0_22:6) | -0.40647549 | 1.729221003 |
| PA_508n | PA | (10:1e_8:0) | -0.39707095 | 1.681770067 |
| PC_2534p | PC | (20:4_13:0) | -0.39067295 | 1.234928192 |
| PE_3359p | PE | (36:4) | -0.39067295 | 1.234928192 |
| FA_176n | FA | (22:6) | -0.38023307 | 1.577315259 |
| dMePE_1968n | dMePE | (20:3_20:4) | -0.37791804 | 1.14494882 |
| TG_4691p | TG | (29:0_11:4_18:3) | -0.37533738 | 1.438695786 |
| LPA_217n | LPA | (18:1) | -0.36878191 | 1.56096808 |
| dMePE_1484n | dMePE | (22:2) | -0.36257008 | 1.358508889 |
| PE_3475p | PE | (24:2_14:3) | -0.357552 | 1.903118089 |
| SM_1409n | SM | (d42:2) | -0.35443073 | 2.24771756 |
| PC_3082p | PC | (22:6_22:6) | -0.35363695 | 1.602660936 |
| PE_3561p | PE | (39:4e) | -0.35314683 | 2.838507516 |
| PE_1109n | PE | (22:6_22:6) | -0.3479233 | 1.623460655 |
| LPA_223n | LPA | (20:4) | -0.34178852 | 1.639192225 |
| PA_512n | PA | (8:0e_12:4) | -0.34178852 | 1.639192225 |
| PE_868n | PE | (16:1_20:4) | -0.34158505 | 1.353311433 |
| dMePE_1689n | dMePE | (14:0_20:5) | -0.34158505 | 1.353311433 |
| ChE_197p | ChE | (20:5) | -0.3337374 | 2.852263363 |
| PE_3261p | PE | (22:0_12:1) | -0.33238245 | 1.626634806 |
| PC_3056p | PC | (20:5_22:5) | -0.3301486 | 2.336792911 |
| ChE_203p | ChE | (22:6) | -0.32698132 | 1.777567366 |
| PE_1079n | PE | (20:4_22:6) | -0.32098176 | 1.594401744 |
| LdMePE_380n | LdMePE | (16:0) | -0.30869779 | 1.298855162 |
| LPE_338n | LPE | (20:4) | -0.28506892 | 2.030704714 |
| MePC_2234p | MePC | (37:4e) | -0.28488111 | 3.678263668 |
| PC_3040p | PC | (20:4e_20:3) | -0.28488111 | 3.678263668 |
| PE_3613p | PE | (41:4e) | -0.28488111 | 3.678263668 |
| SM_3811p | SM | (d18:1_17:0) | -0.27710959 | 1.698354424 |
| LPC_251n | LPC | (22:6) | -0.27548615 | 1.72901627 |
| MePC_2178p | MePC | (36:5) | -0.27443917 | 1.909112275 |
| dMePE_1494n | dMePE | (16:0_16:0) | -0.27267056 | 2.324417478 |
| PG_1246n | PG | (20:3_22:6) | -0.27156879 | 1.578030582 |
| cPA_1454n | cPA | (18:2) | -0.27008916 | 1.229003393 |
| SM_3867p | SM | (t40:5) | -0.26905819 | 1.886135393 |
| ChE_194p | ChE | (20:4) | -0.26860369 | 1.953326501 |
| ZyE_4882p | ZyE | (20:3) | -0.26808763 | 1.96390328 |
| PE_968n | PE | (18:1e_20:4) | -0.26670472 | 1.51218852 |
| dMePE_1523n | dMePE | (32:2) | -0.25342959 | 1.402378376 |
| PE_774n | PE | (18:0_18:0) | -0.24773527 | 1.245377562 |
| MG_1832p | MG | (32:0) | -0.23769156 | 1.202401971 |
| PC_2998p | PC | (39:6) | -0.2356957 | 2.224368338 |
| MGDG_477n | MGDG | (49:5) | -0.23055374 | 1.8471757 |
| WE_4840p | WE | (24:4_14:1) | -0.22881869 | 1.692538383 |
| WE_4838p | WE | (20:1_18:3) | -0.22862438 | 2.137411399 |
| PA_510n | PA | (8:1e_10:1) | -0.22404027 | 1.577275168 |
| LPA_219n | LPA | (18:2) | -0.22404027 | 1.577275168 |
| LPE_292n | LPE | (18:0) | -0.22092998 | 0.986834987 |
| Cer_116p | Cer | (d18:2_24:2) | -0.21996568 | 2.043220928 |
| PE_3630p | PE | (20:0p_22:6) | -0.21907512 | 1.989045903 |
| Cer_72p | Cer | (d16:0_18:2) | -0.21705898 | 1.892817292 |
| LPC_245n | LPC | (20:4) | -0.21675413 | 1.371252601 |
| DG_728p | DG | (40:6e) | -0.21667877 | 1.501134783 |
| LdMePE_442n | LdMePE | (20:4) | -0.20958735 | 1.498226603 |
| CmE_205p | CmE | (10:0) | -0.20884589 | 1.661475982 |
| LPA_226n | LPA | (22:6) | -0.20712566 | 1.366507297 |
| MePC_2292p | MePC | (41:7) | -0.20567503 | 1.726770593 |
| PC_3079p | PC | (22:5_22:5) | -0.20567503 | 1.716611313 |
| PC_2922p | PC | (38:5e) | -0.20086871 | 1.948970365 |
| PE_3615p | PE | (41:5e) | -0.20086871 | 1.948970365 |
| LdMePE_439n | LdMePE | (20:3) | -0.20007784 | 1.881227671 |
| LPC_250n | LPC | (22:6) | -0.1906174 | 1.540592838 |
| LdMePE_458n | LdMePE | (22:6) | -0.19052191 | 1.357119581 |
| SM_3849p | SM | (d18:1_24:1) | -0.18788013 | 2.215188403 |
| PE_3581p | PE | (16:2e_24:2) | -0.18696293 | 1.745417189 |
| WE_4824p | WE | (20:2_16:1) | -0.18674262 | 2.040892224 |
| PC_672n | PC | (22:6_22:6) | -0.18607242 | 1.195247381 |
| PE_3598p | PE | (18:1_22:6) | -0.18555565 | 1.969764087 |
| PE_3622p | PE | (20:4_22:6) | -0.18555565 | 1.969764087 |
| TG_4050p | TG | (35:0e) | -0.18421772 | 1.721296551 |
| PS_1349n | PS | (18:0_18:2) | -0.18202396 | 1.616489951 |
| PE_3490p | PE | (16:0_22:6) | -0.18171092 | 1.306011571 |
| MePC_2054p | MePC | (34:6) | -0.18171092 | 1.304499081 |
| PC_2526p | PC | (19:1_14:1) | -0.18120307 | 1.574228033 |
| MePC_1946p | MePC | (32:5) | -0.17578321 | 1.494012281 |
| dMePE_1677n | dMePE | (18:4_16:0) | -0.17479708 | 1.355982743 |
| FA_144n | FA | (17:0) | -0.17300439 | 1.355029023 |
| MePC_2037p | MePC | (34:3) | -0.169925 | 0.850765677 |
| PE_3443p | PE | (20:1_18:2) | -0.169925 | 0.850765677 |
| FA_145n | FA | (17:0) | -0.168337 | 2.419728763 |
| PC_666n | PC | (20:4_22:6) | -0.16821504 | 1.353766007 |
| PG_1151n | PG | (20:0) | -0.16061622 | 1.475895053 |
| dMePE_1990n | dMePE | (22:6_22:6) | -0.15728079 | 1.255705412 |
| LPC_1054p | LPC | (20:3) | -0.15200309 | 1.684576634 |
| PE_3229p | PE | (12:1e_11:2) | -0.15200309 | 1.684576634 |
| PE_3296p | PE | (17:0_18:1) | -0.15100779 | 1.732220969 |
| PC_3083p | PC | (22:6_22:6) | -0.15067891 | 2.105410241 |
| PC_3084p | PC | (44:12) | -0.15067891 | 2.105410241 |
| DG_571p | DG | (18:0_17:0) | -0.14248693 | 2.875928202 |
| MLCL_488n | MLCL | (18:2_18:2_18:2) | -0.14076429 | 2.046174629 |
| PC_2524p | PC | (25:0_8:0) | -0.14004796 | 1.860048202 |
| LPC_1067p | LPC | (20:4) | -0.13806792 | 1.729789345 |
| MePC_1883p | MePC | (17:1e) | -0.13806792 | 1.729789345 |
| CmE_207p | CmE | (14:0) | -0.13750352 | 1.707163007 |
| Cer_138p | Cer | (m19:1_23:3) | -0.1299303 | 1.767614507 |
| Hex2Cer_784p | Hex2Cer | (d29:1) | -0.12980735 | 0.988902281 |
| PC_2717p | PC | (36:0) | -0.12928302 | 2.308404445 |
| LdMePE_441n | LdMePE | (20:4) | -0.12553088 | 1.520436234 |
| dMePE_1977n | dMePE | (20:4_22:6) | -0.12517356 | 1.319404124 |
| SM_3855p | SM | (d42:3) | -0.12401919 | 1.996386496 |
| LdMePE_416n | LdMePE | (18:2) | -0.1186445 | 1.491612361 |
| FA_172n | FA | (22:6) | -0.11560404 | 1.533143407 |
| MePC_2279p | MePC | (39:7) | -0.11414754 | 2.412951335 |
| dMePE_1987n | dMePE | (20:3_22:6) | -0.11321061 | 1.194593133 |
| PC_3057p | PC | (20:5_22:5) | -0.10983455 | 2.281920376 |
| Hex2Cer_844p | Hex2Cer | (d13:0_21:5) | -0.10983455 | 2.26439924 |
| LPC_1068p | LPC | (20:4) | -0.10942283 | 1.371684377 |
| PE_3479p | PE | (24:1_14:4) | -0.10930972 | 1.677767882 |
| PE_3605p | PE | (20:4_20:4) | -0.10930972 | 1.677767882 |
| FA_152n | FA | (20:4) | -0.10877635 | 1.756803251 |
| PC_664n | PC | (18:2_22:6) | -0.10638934 | 2.363500438 |
| MePC_2160p | MePC | (35:5) | -0.10516093 | 1.615204958 |
| PS_1378n | PS | (18:0_22:6) | -0.10475774 | 3.125165133 |
| dMePE_1882n | dMePE | (18:1_20:4) | -0.10350458 | 1.553142063 |
| Hex2Cer_813p | Hex2Cer | (d31:1) | -0.10142773 | 2.470100232 |
| dMePE_1532n | dMePE | (16:0_17:0) | -0.10122947 | 2.202799432 |
| PE_1009n | PE | (17:0_22:6) | -0.09740406 | 2.463400003 |
| PE_1073n | PE | (18:2_22:6) | -0.09725067 | 2.832643524 |
| MePC_2169p | MePC | (36:3) | -0.0961237 | 2.110811279 |
| PC_2999p | PC | (39:6) | -0.0961237 | 2.110811279 |
| PE_3572p | PE | (40:3) | -0.0961237 | 2.110811279 |
| PE_879n | PE | (17:0_20:4) | -0.09505525 | 1.88274259 |
| MePC_2223p | MePC | (37:4) | -0.09463687 | 2.729838966 |
| PC_2436p | PC | (10:0e_10:4) | -0.09399206 | 2.038977911 |
| MePC_2275p | MePC | (39:6) | -0.09214471 | 2.35603048 |
| PC_3076p | PC | (22:5_20:4) | -0.09214471 | 2.35603048 |
| FA_153n | FA | (20:4) | -0.09096363 | 1.038132333 |
| LPC_1065p | LPC | (20:4) | -0.09067932 | 1.955854258 |
| dMePE_1835n | dMePE | (17:0_20:4) | -0.08892026 | 1.647739165 |
| MePC_2031p | MePC | (34:1) | -0.08888128 | 1.519801554 |
| Hex2Cer_841p | Hex2Cer | (d34:4) | -0.08387256 | 1.97868661 |
| MePC_1882p | MePC | (17:1e) | -0.08101469 | 1.534878165 |
| dMePE_1973n | dMePE | (18:2_22:6) | -0.07733713 | 2.444902865 |
| PC_3042p | PC | (18:3_22:5) | -0.0768936 | 2.771504765 |
| PE_1070n | PE | (18:1e_22:6) | -0.07434734 | 2.961973004 |
| MePC_2258p | MePC | (38:9) | -0.07352904 | 2.210149265 |
| dMePE_1894n | dMePE | (18:2_20:4) | -0.06900777 | 2.092748101 |
| MePC_2236p | MePC | (37:5) | -0.06473626 | 2.782345533 |
| PC_2913p | PC | (38:5) | -0.06473626 | 2.782345533 |
| MePC_2108p | MePC | (35:3) | -0.05277703 | 2.093101908 |
| PI_1288n | PI | (18:0_20:4) | -0.03851524 | 2.78139426 |
| PI_1289n | PI | (18:0_20:4) | -0.03851524 | 2.78139426 |
| **Up regulated lipid species** | | | | |
| **Lipid ID** | **Lipid Class** | **FattyAcid** | **log2(FC)** | **log10 (p value)** |

| PG_1241n | PG | (20:5_20:4) | 2.216656052 | 1.437818705 |
| --- | --- | --- | --- | --- |
| SPH_3893p | SPH | (t16:0) | 2.195015982 | 1.110056307 |
| PG_1226n | PG | (16:1_22:6) | 1.906890596 | 0.949217275 |
| MG_1222p | MG | (16:0) | 1.85010457 | 0.963120401 |
| TG_3939p | TG | (4:0_11:1_12:0) | 1.678071905 | 1.219525748 |
| dMePE_1612n | dMePE | (16:0_18:2) | 1.567684509 | 1.723586297 |
| PE_3528p | PE | (19:0_20:3) | 1.509674373 | 1.744664418 |
| MG_1514p | MG | (28:0) | 1.479992941 | 1.512772826 |
| DG_568p | DG | (14:0_20:4) | 1.419225296 | 1.322594663 |
| MG_1424p | MG | (19:1) | 1.40275917 | 1.186241852 |
| TG_4046p | TG | (14:0e_8:0_12:4) | 1.377304852 | 1.52835393 |
| PC_2723p | PC | (27:1_9:0) | 1.333126301 | 1.18901787 |
| MePC_1907p | MePC | (29:0) | 1.245476034 | 2.945853593 |
| TG_4111p | TG | (12:0e_12:3_12:3) | 1.244307198 | 2.173072841 |
| MePC_2102p | MePC | (35:3) | 1.203283598 | 0.620760036 |
| MG_1261p | MG | (16:1) | 1.143835773 | 3.599204384 |
| DG_465p | DG | (16:1_16:1) | 1.043327432 | 1.357168184 |
| MG_1259p | MG | (16:0e) | 1.022367813 | 2.236830728 |
| MG_1747p | MG | (29:1) | 0.961931959 | 1.2735316 |
| DG_667p | DG | (14:0_22:6) | 0.928446739 | 1.727416254 |
| LPC_955p | LPC | (18:0) | 0.916806064 | 1.010992859 |
| TG_4113p | TG | (36:6e) | 0.909989834 | 1.395419294 |
| PC_2837p | PC | (16:1_20:4) | 0.907333752 | 1.185612461 |
| PS_1362n | PS | (38:5) | 0.897901812 | 3.978599702 |
| LPC_973p | LPC | (18:1) | 0.884522783 | 1.194874615 |
| DG_334p | DG | (22:1) | 0.854149134 | 1.312106041 |
| MG_1348p | MG | (18:0) | 0.846087317 | 2.714977726 |
| PA_533n | PA | (16:0_20:5) | 0.819427754 | 1.530638917 |
| MG_1411p | MG | (18:3e) | 0.757143476 | 2.070908717 |
| TG_3946p | TG | (4:0_10:2_16:0) | 0.739610315 | 2.422184954 |
| PE_725n | PE | (14:0_18:2) | 0.716207034 | 2.40720643 |
| DG_522p | DG | (16:0_18:1) | 0.70268446 | 0.967606844 |
| MG_1445p | MG | (20:1) | 0.699694717 | 2.024213711 |
| TG_3930p | TG | (4:0_8:0_9:0) | 0.679113938 | 1.184082563 |
| PG_1175n | PG | (18:1_18:1) | 0.675236604 | 0.9339243 |
| PE_967n | PE | (16:0e_22:5) | 0.654671473 | 1.663797263 |
| TG_4522p | TG | (16:0_16:0_20:5) | 0.638204522 | 1.63313954 |
| WE_4807p | WE | (3:0_20:2) | 0.627341103 | 0.769370181 |
| TG_3954p | TG | (31:2) | 0.563429339 | 1.471293413 |
| TG_3986p | TG | (16:1_6:0_11:4) | 0.563429339 | 1.471293413 |
| LPC_888p | LPC | (16:0) | 0.554588852 | 2.033396631 |
| TG_3958p | TG | (32:0e) | 0.552541023 | 2.378239828 |
| DG_446p | DG | (16:0_16:0) | 0.507155701 | 3.259663212 |
| DG_283p | DG | (19:1) | 0.492280498 | 1.292060361 |
| TG_3929p | TG | (19:0) | 0.475579041 | 1.29689645 |
| DG_344p | DG | (12:0_10:3) | 0.456176194 | 2.173713396 |
| DG_561p | DG | (16:1_18:2) | 0.454417032 | 1.939849987 |
| MG_1263p | MG | (16:1) | 0.447786082 | 1.221014104 |
| DG_665p | DG | (16:1_20:4) | 0.447112965 | 1.969407709 |
| PC_610n | PC | (16:1_20:4) | 0.43673257 | 1.607674989 |
| PS_1366n | PS | (16:0_22:6) | 0.430239906 | 1.21747347 |
| DG_712p | DG | (39:2) | 0.423101774 | 1.282558813 |
| PG_1160n | PG | (16:1_18:1) | 0.419903254 | 1.400518848 |
| PI_1257n | PI | (16:2e_16:0) | 0.401616984 | 2.081591811 |
| TG_3966p | TG | (32:1e) | 0.393011193 | 0.842485021 |
| DG_458p | DG | (18:1_14:0) | 0.381178695 | 1.826695915 |
| AcCa_13p | AcCa | (20:3) | 0.377367081 | 2.132275586 |
| DG_575p | DG | (17:1_18:1) | 0.376148486 | 1.166268063 |
| TG_4052p | TG | (12:1e_6:0_17:1) | 0.376148486 | 1.166268063 |
| PIP_1324n | PIP | (18:0e_15:0) | 0.373617572 | 1.963647549 |
| TG_3962p | TG | (12:1e_6:0_14:0) | 0.371297184 | 1.804808608 |
| PG_3663p | PG | (16:0_18:1) | 0.345644164 | 2.098302238 |
| DG_468p | DG | (18:2_14:1) | 0.344264425 | 3.522296482 |
| TG_3970p | TG | (12:1e_10:1_10:1) | 0.344264425 | 3.522296482 |
| Cer_35n | Cer | (d18:1_24:0) | 0.337581359 | 1.60881081 |
| cPA_1453n | cPA | (18:1) | 0.305413008 | 2.173747055 |
| BisMePA_36p | BisMePA | (28:0e) | 0.303010882 | 1.496344765 |
| DG_466p | DG | (8:1e_24:1) | 0.300704824 | 1.843700314 |
| MGDG_478n | MGDG | (58:9) | 0.300013318 | 2.33628585 |
| PE_722n | PE | (16:0_16:1) | 0.289029614 | 1.828183223 |
| dMePE_1489n | dMePE | (16:1_14:0) | 0.289029614 | 1.829798731 |
| TG_4161p | TG | (14:1e_11:1_14:4) | 0.286304185 | 3.35615151 |
| LPC_1031p | LPC | (18:4) | 0.285560975 | 1.497675263 |
| MGMG_480n | MGMG | (37:1) | 0.282530581 | 2.796252381 |
| PI_1269n | PI | (16:0_20:3) | 0.272525855 | 1.010004108 |
| PE_874n | PE | (16:1_20:5) | 0.266992419 | 1.412122323 |
| MePC_2015p | MePC | (33:2) | 0.26607486 | 1.442750723 |
| DG_430p | DG | (30:0e) | 0.265928479 | 1.429623272 |
| MG_1801p | MG | (30:0) | 0.265928479 | 1.429623272 |
| Cer_23n | Cer | (d18:2_22:0) | 0.261066199 | 1.012070011 |
| Cer_20n | Cer | (d18:1_22:0) | 0.257981178 | 2.620241551 |
| DG_562p | DG | (16:1_18:2) | 0.257157839 | 1.634619074 |
| TG_4042p | TG | (14:0e_10:1_10:2) | 0.257157839 | 1.634619074 |
| TG_4221p | TG | (4:0_20:4_20:4) | 0.250406798 | 2.924943016 |
| PS_1384n | PS | (20:1_22:6) | 0.23878686 | 1.268383075 |
| SPH_3888p | SPH | (d18:1) | 0.230883882 | 1.296483412 |
| DG_567p | DG | (16:1_18:3) | 0.229587923 | 2.379692751 |
| PE_728n | PE | (18:0_16:0) | 0.226571121 | 1.468762689 |
| TG_4045p | TG | (14:0e_9:0_11:4) | 0.225985686 | 2.454378706 |
| TG_3968p | TG | (12:1e_6:0_14:1) | 0.21818017 | 2.0937105 |
| TG_3964p | TG | (12:1e_6:0_14:0) | 0.21818017 | 1.387512954 |
| DG_464p | DG | (16:1_16:1) | 0.215646903 | 2.0893996 |
| DG_657p | DG | (18:2_18:2) | 0.211855682 | 2.098719735 |
| TG_4103p | TG | (12:1e_6:0_18:3) | 0.211855682 | 2.098719735 |
| DG_576p | DG | (17:1_18:2) | 0.207804885 | 2.049386437 |
| TG_4054p | TG | (18:1e_6:0_11:2) | 0.207804885 | 2.049386437 |
| PC_569n | PC | (16:0_16:1) | 0.20292512 | 1.561542507 |
| CerP_67n | CerP | (d36:2) | 0.20270179 | 1.991931794 |
| PE_723n | PE | (16:0_16:1) | 0.20270179 | 1.991931794 |
| DG_431p | DG | (16:1_14:0) | 0.201234278 | 1.409910489 |
| TG_3944p | TG | (14:1e_8:0_8:0) | 0.201234278 | 1.409910489 |
| TG_3963p | TG | (12:1e_6:0_14:0) | 0.194397694 | 1.760907006 |
| DG_531p | DG | (16:1_18:1) | 0.182636053 | 1.810853676 |
| TG_4023p | TG | (12:1e_6:0_16:1) | 0.182636053 | 1.810853676 |
| MGMG_479n | MGMG | (22:0) | 0.175771259 | 3.184306511 |
| DG_457p | DG | (18:1_14:0) | 0.166608166 | 1.675400441 |
| DG_700p | DG | (18:2_20:4) | 0.165825089 | 1.94245089 |
| TG_4148p | TG | (18:0e_10:3_10:3) | 0.165825089 | 1.94245089 |
| DG_502p | DG | (16:0_18:1) | 0.160936218 | 1.464371016 |
| MePC_2053p | MePC | (34:6) | 0.160788207 | 1.908475707 |
| Hex2Cer_767p | Hex2Cer | (d13:0_13:0) | 0.16073583 | 1.519995575 |
| Cer_30n | Cer | (d18:0_24:0) | 0.158781002 | 2.130426702 |
| DG_560p | DG | (16:1_18:2) | 0.157771219 | 1.939910687 |
| PI_1258n | PI | (18:0_16:1) | 0.156342029 | 1.274301153 |
| Hex1Cer_752p | Hex1Cer | (d37:5) | 0.155582984 | 2.108630739 |
| CerP_98n | CerP | (d44:2) | 0.14385622 | 2.087343954 |
| PE_1011n | PE | (18:1_22:0) | 0.14385622 | 2.087343954 |
| dMePE_1839n | dMePE | (18:0_20:1) | 0.14385622 | 2.087343954 |
| PI_1300n | PI | (18:0_20:5) | 0.14139394 | 2.502161912 |
| MePC_1930p | MePC | (32:1) | 0.138130919 | 2.126226758 |
| MG_1830p | MG | (32:0) | 0.134460321 | 1.227792314 |
| DG_569p | DG | (34:4e) | 0.130907258 | 1.616336378 |
| MG_1842p | MG | (32:1) | 0.130907258 | 1.616336378 |
| DG_574p | DG | (17:1_18:1) | 0.125763857 | 1.543818409 |
| TG_4053p | TG | (12:1e_6:0_17:1) | 0.125763857 | 1.543818409 |
| PC_2839p | PC | (22:1_14:4) | 0.123701985 | 1.401483604 |
| PA_514n | PA | (16:0_18:1) | 0.120294234 | 3.84399967 |
| PE_749n | PE | (16:0_18:2) | 0.118644496 | 1.494067987 |
| MePC_1909p | MePC | (30:2) | 0.117100351 | 2.05937531 |
| PA_520n | PA | (18:0_18:1) | 0.108252891 | 5.260517161 |
| MePC_2195p | MePC | (36:8) | 0.106915204 | 1.785084391 |
| dMePE_1830n | dMePE | (19:0_18:1) | 0.106199404 | 2.438017358 |
| DG_474p | DG | (15:0_18:2) | 0.105154746 | 2.743325458 |
| TG_3976p | TG | (16:0e_6:0_11:2) | 0.100694 | 2.867449978 |
| TG_4002p | TG | (12:1e_8:0_14:0) | 0.099419369 | 1.393179968 |
| DG_507p | DG | (16:0_18:1) | 0.099187575 | 1.427556677 |
| DG_501p | DG | (16:0_18:1) | 0.097562003 | 1.388656079 |
| dMePE_1517n | dMePE | (16:0_16:1) | 0.096753233 | 1.387829357 |
| PC_2508p | PC | (18:0_14:1) | 0.096676019 | 2.300198755 |
| PC_574n | PC | (16:0_18:1) | 0.096278289 | 3.084356573 |
| PC_575n | PC | (16:0_18:1) | 0.096278289 | 3.06997366 |
| DG_656p | DG | (18:2_18:2) | 0.094434802 | 1.795071615 |
| TG_4104p | TG | (12:1e_6:0_18:3) | 0.094434802 | 1.795071615 |
| MG_1267p | MG | (16:2e) | 0.094297297 | 1.456629665 |
| PI_1272n | PI | (16:0_20:4) | 0.090892581 | 1.938469954 |
| DG_530p | DG | (16:1_18:1) | 0.090146556 | 1.984090785 |
| TG_4022p | TG | (12:1e_6:0_16:1) | 0.090146556 | 1.984090785 |
| TG_3975p | TG | (18:1e_6:0_9:0) | 0.086952544 | 1.706897677 |
| Cer_34n | Cer | (d18:1_24:0) | 0.070808168 | 1.734566947 |
| PC_594n | PC | (18:1_18:1) | 0.063725598 | 1.55094607 |
| PC_2722p | PC | (20:0_16:1) | 0.055763272 | 1.56802515 |
| PI_1288n | PI | (18:0_20:4) | -0.03851524 | 2.78139426 |
| PI_1289n | PI | (18:0_20:4) | -0.03851524 | 2.78139426 |
| MePC_2108p | MePC | (35:3) | -0.05277703 | 2.093101908 |
| MePC_2236p | MePC | (37:5) | -0.06473626 | 2.782345533 |
| PC_2913p | PC | (38:5) | -0.06473626 | 2.782345533 |
| dMePE_1894n | dMePE | (18:2_20:4) | -0.06900777 | 2.092748101 |
| MePC_2258p | MePC | (38:9) | -0.07352904 | 2.210149265 |
| PE_1070n | PE | (18:1e_22:6) | -0.07434734 | 2.961973004 |
| PC_3042p | PC | (18:3_22:5) | -0.0768936 | 2.771504765 |
| dMePE_1973n | dMePE | (18:2_22:6) | -0.07733713 | 2.444902865 |
| MePC_1882p | MePC | (17:1e) | -0.08101469 | 1.534878165 |
| Hex2Cer_841p | Hex2Cer | (d34:4) | -0.08387256 | 1.97868661 |
| MePC_2031p | MePC | (34:1) | -0.08888128 | 1.519801554 |
| dMePE_1835n | dMePE | (17:0_20:4) | -0.08892026 | 1.647739165 |
| LPC_1065p | LPC | (20:4) | -0.09067932 | 1.955854258 |
| FA_153n | FA | (20:4) | -0.09096363 | 1.038132333 |
| MePC_2275p | MePC | (39:6) | -0.09214471 | 2.35603048 |
| PC_3076p | PC | (22:5_20:4) | -0.09214471 | 2.35603048 |
| PC_2436p | PC | (10:0e_10:4) | -0.09399206 | 2.038977911 |
| MePC_2223p | MePC | (37:4) | -0.09463687 | 2.729838966 |
| PE_879n | PE | (17:0_20:4) | -0.09505525 | 1.88274259 |
| MePC_2169p | MePC | (36:3) | -0.0961237 | 2.110811279 |
| PC_2999p | PC | (39:6) | -0.0961237 | 2.110811279 |
| PE_3572p | PE | (40:3) | -0.0961237 | 2.110811279 |
| PE_1073n | PE | (18:2_22:6) | -0.09725067 | 2.832643524 |
| PE_1009n | PE | (17:0_22:6) | -0.09740406 | 2.463400003 |
| dMePE_1532n | dMePE | (16:0_17:0) | -0.10122947 | 2.202799432 |
| Hex2Cer_813p | Hex2Cer | (d31:1) | -0.10142773 | 2.470100232 |
| dMePE_1882n | dMePE | (18:1_20:4) | -0.10350458 | 1.553142063 |
| PS_1378n | PS | (18:0_22:6) | -0.10475774 | 3.125165133 |
| MePC_2160p | MePC | (35:5) | -0.10516093 | 1.615204958 |
| PC_664n | PC | (18:2_22:6) | -0.10638934 | 2.363500438 |
| FA_152n | FA | (20:4) | -0.10877635 | 1.756803251 |
| PE_3479p | PE | (24:1_14:4) | -0.10930972 | 1.677767882 |
| PE_3605p | PE | (20:4_20:4) | -0.10930972 | 1.677767882 |
| LPC_1068p | LPC | (20:4) | -0.10942283 | 1.371684377 |
| PC_3057p | PC | (20:5_22:5) | -0.10983455 | 2.281920376 |
| Hex2Cer_844p | Hex2Cer | (d13:0_21:5) | -0.10983455 | 2.26439924 |
| dMePE_1987n | dMePE | (20:3_22:6) | -0.11321061 | 1.194593133 |
| MePC_2279p | MePC | (39:7) | -0.11414754 | 2.412951335 |
| FA_172n | FA | (22:6) | -0.11560404 | 1.533143407 |
| LdMePE_416n | LdMePE | (18:2) | -0.1186445 | 1.491612361 |
| SM_3855p | SM | (d42:3) | -0.12401919 | 1.996386496 |
| dMePE_1977n | dMePE | (20:4_22:6) | -0.12517356 | 1.319404124 |
| LdMePE_441n | LdMePE | (20:4) | -0.12553088 | 1.520436234 |
| PC_2717p | PC | (36:0) | -0.12928302 | 2.308404445 |
| Hex2Cer_784p | Hex2Cer | (d29:1) | -0.12980735 | 0.988902281 |
| Cer_138p | Cer | (m19:1_23:3) | -0.1299303 | 1.767614507 |
| CmE_207p | CmE | (14:0) | -0.13750352 | 1.707163007 |
| LPC_1067p | LPC | (20:4) | -0.13806792 | 1.729789345 |
| MePC_1883p | MePC | (17:1e) | -0.13806792 | 1.729789345 |
| PC_2524p | PC | (25:0_8:0) | -0.14004796 | 1.860048202 |
| MLCL_488n | MLCL | (18:2_18:2_18:2) | -0.14076429 | 2.046174629 |
| DG_571p | DG | (18:0_17:0) | -0.14248693 | 2.875928202 |
| PC_3083p | PC | (22:6_22:6) | -0.15067891 | 2.105410241 |
| PC_3084p | PC | (44:12) | -0.15067891 | 2.105410241 |
| PE_3296p | PE | (17:0_18:1) | -0.15100779 | 1.732220969 |
| LPC_1054p | LPC | (20:3) | -0.15200309 | 1.684576634 |
| PE_3229p | PE | (12:1e_11:2) | -0.15200309 | 1.684576634 |
| dMePE_1990n | dMePE | (22:6_22:6) | -0.15728079 | 1.255705412 |
| PG_1151n | PG | (20:0) | -0.16061622 | 1.475895053 |
| PC_666n | PC | (20:4_22:6) | -0.16821504 | 1.353766007 |
| FA_145n | FA | (17:0) | -0.168337 | 2.419728763 |
| MePC_2037p | MePC | (34:3) | -0.169925 | 0.850765677 |
| PE_3443p | PE | (20:1_18:2) | -0.169925 | 0.850765677 |
| FA_144n | FA | (17:0) | -0.17300439 | 1.355029023 |
| dMePE_1677n | dMePE | (18:4_16:0) | -0.17479708 | 1.355982743 |
| MePC_1946p | MePC | (32:5) | -0.17578321 | 1.494012281 |
| PC_2526p | PC | (19:1_14:1) | -0.18120307 | 1.574228033 |
| PE_3490p | PE | (16:0_22:6) | -0.18171092 | 1.306011571 |
| MePC_2054p | MePC | (34:6) | -0.18171092 | 1.304499081 |
| PS_1349n | PS | (18:0_18:2) | -0.18202396 | 1.616489951 |
| TG_4050p | TG | (35:0e) | -0.18421772 | 1.721296551 |
| PE_3598p | PE | (18:1_22:6) | -0.18555565 | 1.969764087 |
| PE_3622p | PE | (20:4_22:6) | -0.18555565 | 1.969764087 |
| PC_672n | PC | (22:6_22:6) | -0.18607242 | 1.195247381 |
| WE_4824p | WE | (20:2_16:1) | -0.18674262 | 2.040892224 |
| PE_3581p | PE | (16:2e_24:2) | -0.18696293 | 1.745417189 |
| SM_3849p | SM | (d18:1_24:1) | -0.18788013 | 2.215188403 |
| LdMePE_458n | LdMePE | (22:6) | -0.19052191 | 1.357119581 |
| LPC_250n | LPC | (22:6) | -0.1906174 | 1.540592838 |
| LdMePE_439n | LdMePE | (20:3) | -0.20007784 | 1.881227671 |
| PC_2922p | PC | (38:5e) | -0.20086871 | 1.948970365 |
| PE_3615p | PE | (41:5e) | -0.20086871 | 1.948970365 |
| MePC_2292p | MePC | (41:7) | -0.20567503 | 1.726770593 |
| PC_3079p | PC | (22:5_22:5) | -0.20567503 | 1.716611313 |
| LPA_226n | LPA | (22:6) | -0.20712566 | 1.366507297 |
| CmE_205p | CmE | (10:0) | -0.20884589 | 1.661475982 |
| LdMePE_442n | LdMePE | (20:4) | -0.20958735 | 1.498226603 |
| DG_728p | DG | (40:6e) | -0.21667877 | 1.501134783 |
| LPC_245n | LPC | (20:4) | -0.21675413 | 1.371252601 |
| Cer_72p | Cer | (d16:0_18:2) | -0.21705898 | 1.892817292 |
| PE_3630p | PE | (20:0p_22:6) | -0.21907512 | 1.989045903 |
| Cer_116p | Cer | (d18:2_24:2) | -0.21996568 | 2.043220928 |
| LPE_292n | LPE | (18:0) | -0.22092998 | 0.986834987 |
| PA_510n | PA | (8:1e_10:1) | -0.22404027 | 1.577275168 |
| LPA_219n | LPA | (18:2) | -0.22404027 | 1.577275168 |
| WE_4838p | WE | (20:1_18:3) | -0.22862438 | 2.137411399 |
| WE_4840p | WE | (24:4_14:1) | -0.22881869 | 1.692538383 |
| MGDG_477n | MGDG | (49:5) | -0.23055374 | 1.8471757 |
| PC_2998p | PC | (39:6) | -0.2356957 | 2.224368338 |
| MG_1832p | MG | (32:0) | -0.23769156 | 1.202401971 |
| PE_774n | PE | (18:0_18:0) | -0.24773527 | 1.245377562 |
| dMePE_1523n | dMePE | (32:2) | -0.25342959 | 1.402378376 |
| PE_968n | PE | (18:1e_20:4) | -0.26670472 | 1.51218852 |
| ZyE_4882p | ZyE | (20:3) | -0.26808763 | 1.96390328 |
| ChE_194p | ChE | (20:4) | -0.26860369 | 1.953326501 |
| SM_3867p | SM | (t40:5) | -0.26905819 | 1.886135393 |
| cPA_1454n | cPA | (18:2) | -0.27008916 | 1.229003393 |
| PG_1246n | PG | (20:3_22:6) | -0.27156879 | 1.578030582 |
| dMePE_1494n | dMePE | (16:0_16:0) | -0.27267056 | 2.324417478 |
| MePC_2178p | MePC | (36:5) | -0.27443917 | 1.909112275 |
| LPC_251n | LPC | (22:6) | -0.27548615 | 1.72901627 |
| SM_3811p | SM | (d18:1_17:0) | -0.27710959 | 1.698354424 |
| MePC_2234p | MePC | (37:4e) | -0.28488111 | 3.678263668 |
| PC_3040p | PC | (20:4e_20:3) | -0.28488111 | 3.678263668 |
| PE_3613p | PE | (41:4e) | -0.28488111 | 3.678263668 |
| LPE_338n | LPE | (20:4) | -0.28506892 | 2.030704714 |
| LdMePE_380n | LdMePE | (16:0) | -0.30869779 | 1.298855162 |
| PE_1079n | PE | (20:4_22:6) | -0.32098176 | 1.594401744 |
| ChE_203p | ChE | (22:6) | -0.32698132 | 1.777567366 |
| PC_3056p | PC | (20:5_22:5) | -0.3301486 | 2.336792911 |
| PE_3261p | PE | (22:0_12:1) | -0.33238245 | 1.626634806 |
| ChE_197p | ChE | (20:5) | -0.3337374 | 2.852263363 |
| PE_868n | PE | (16:1_20:4) | -0.34158505 | 1.353311433 |
| dMePE_1689n | dMePE | (14:0_20:5) | -0.34158505 | 1.353311433 |
| LPA_223n | LPA | (20:4) | -0.34178852 | 1.639192225 |
| PA_512n | PA | (8:0e_12:4) | -0.34178852 | 1.639192225 |
| PE_1109n | PE | (22:6_22:6) | -0.3479233 | 1.623460655 |
| PE_3561p | PE | (39:4e) | -0.35314683 | 2.838507516 |
| PC_3082p | PC | (22:6_22:6) | -0.35363695 | 1.602660936 |
| SM_1409n | SM | (d42:2) | -0.35443073 | 2.24771756 |
| PE_3475p | PE | (24:2_14:3) | -0.357552 | 1.903118089 |
| dMePE_1484n | dMePE | (22:2) | -0.36257008 | 1.358508889 |
| LPA_217n | LPA | (18:1) | -0.36878191 | 1.56096808 |
| TG_4691p | TG | (29:0_11:4_18:3) | -0.37533738 | 1.438695786 |
| dMePE_1968n | dMePE | (20:3_20:4) | -0.37791804 | 1.14494882 |
| FA_176n | FA | (22:6) | -0.38023307 | 1.577315259 |
| PC_2534p | PC | (20:4_13:0) | -0.39067295 | 1.234928192 |
| PE_3359p | PE | (36:4) | -0.39067295 | 1.234928192 |
| PA_508n | PA | (10:1e_8:0) | -0.39707095 | 1.681770067 |
| PC_659n | PC | (18:0_22:6) | -0.40647549 | 1.729221003 |
| CmE_206p | CmE | (12:0) | -0.40690962 | 2.111682505 |
| PE_3481p | PE | (18:0_20:5) | -0.41932486 | 3.176935332 |
| dMePE_1841n | dMePE | (18:0_20:2) | -0.43785821 | 0.933544845 |
| PC_2882p | PC | (28:0_10:4) | -0.44332654 | 1.302724218 |
| PC_3034p | PC | (40:7) | -0.44332654 | 1.302724218 |
| MePC_2273p | MePC | (39:4e) | -0.45391039 | 2.337212301 |
| LPC_1127p | LPC | (22:6) | -0.45655677 | 1.868572247 |
| Hex2Cer_856p | Hex2Cer | (d45:0+O) | -0.45779807 | 2.765715928 |
| Hex2Cer_865p | Hex2Cer | (t45:0) | -0.45779807 | 2.765715928 |
| PC_2486p | PC | (10:1e_14:4) | -0.45943162 | 1.227866285 |
| PC_616n | PC | (17:0_20:4) | -0.46406307 | 2.796222525 |
| PIP_1327n | PIP | (37:4e) | -0.46496272 | 1.49745207 |
| MePC_1906p | MePC | (21:3e) | -0.48351724 | 1.484784815 |
| PE_3181p | PE | (10:1e_10:2) | -0.50165985 | 0.938928709 |
| CerP_101n | CerP | (d44:3) | -0.50182127 | 0.972452958 |
| SPHP_1430n | SPHP | (m22:1) | -0.50753753 | 0.966835573 |
| cPA_1438n | cPA | (16:0) | -0.50951027 | 1.656362486 |
| LPC_1132p | LPC | (22:6) | -0.51164582 | 2.218687934 |
| LPC_246n | LPC | (20:4) | -0.54167094 | 0.991632342 |
| LPC_244n | LPC | (20:3) | -0.54949782 | 1.356721741 |
| TG_4705p | TG | (20:1_18:1_22:6) | -0.55581616 | 1.685019348 |
| LPC_1128p | LPC | (22:6) | -0.55908387 | 3.033171985 |
| LPC_1078p | LPC | (20:4) | -0.56910855 | 1.885260294 |
| TG_4256p | TG | (46:2) | -0.57994443 | 2.57447416 |
| PA_562n | PA | (20:4_22:6) | -0.58862882 | 1.52989753 |
| PE_3603p | PE | (40:8) | -0.60741735 | 2.464270378 |
| PI_1275n | PI | (18:3_18:2) | -0.62583478 | 0.997603458 |
| PE_768n | PE | (16:0_18:3) | -0.63584367 | 2.215473296 |
| PC_3044p | PC | (40:8) | -0.63911827 | 1.002321928 |
| dMePE_1775n | dMePE | (16:0_20:4) | -0.65298092 | 0.95707941 |
| LPE_1142p | LPE | (16:0) | -0.67442413 | 2.239102174 |
| TG_4105p | TG | (12:1e_6:0_18:3) | -0.68846708 | 1.701295748 |
| DLCL_142n | DLCL | (40:8) | -0.6918777 | 1.143717329 |
| PC_2819p | PC | (22:0_14:4) | -0.70191865 | 1.459273549 |
| LPC_1095p | LPC | (20:5) | -0.70209513 | 1.149506601 |
| LPC_1081p | LPC | (20:4) | -0.70744651 | 1.277613988 |
| TG_4044p | TG | (10:0_12:2_12:2) | -0.71271805 | 1.511137368 |
| dMePE_1974n | dMePE | (18:2_22:6) | -0.71848617 | 1.065486063 |
| PE_3614p | PE | (41:5e) | -0.73426644 | 2.482140097 |
| PIP_1328n | PIP | (43:3e) | -0.73696559 | 1.176891536 |
| PE_3487p | PE | (38:6) | -0.73887518 | 1.256189349 |
| PC_2299p | PC | (15:0) | -0.74908188 | 1.559025885 |
| dMePE_1778n | dMePE | (18:1_18:3) | -0.75032398 | 2.185257877 |
| LPE_254n | LPE | (16:0) | -0.7635598 | 1.088000897 |
| StE_3920p | StE | (16:0) | -0.76733924 | 3.827204131 |
| PE_3372p | PE | (16:0p_20:4) | -0.77230797 | 1.188977913 |
| DG_233p | DG | (16:0e) | -0.77536911 | 0.735875293 |
| MG_1199p | MG | (16:0) | -0.77536911 | 0.735875293 |
| MG_1776p | MG | (29:1) | -0.78068697 | 1.171795264 |
| PE_764n | PE | (16:1_18:2) | -0.78297076 | 1.387065998 |
| dMePE_1698n | dMePE | (17:1_18:2) | -0.78622071 | 1.668503963 |
| LPA_222n | LPA | (20:3) | -0.80145432 | 1.092645305 |
| dMePE_1978n | dMePE | (20:4_22:6) | -0.80851886 | 2.182697755 |
| LdMePE_443n | LdMePE | (20:4) | -0.82086685 | 1.576108326 |
| dMePE_1590n | dMePE | (16:0_18:1) | -0.8326056 | 0.841170297 |
| DG_748p | DG | (54:4) | -0.83471539 | 1.17083938 |
| LdMePE_453n | LdMePE | (22:4) | -0.83561318 | 1.771463169 |
| SM_1389n | SM | (d34:1) | -0.83927969 | 2.206309382 |
| PE_1067n | PE | (18:1_22:6) | -0.84582907 | 1.378988373 |
| PE_3100p | PE | (16:1e) | -0.85913746 | 1.349066581 |
| PC_3045p | PC | (40:8) | -0.88613204 | 0.935463527 |
| dMePE_1897n | dMePE | (16:0_22:6) | -0.92386893 | 1.714337725 |
| MePC_2243p | MePC | (37:6) | -0.9248125 | 1.351300172 |
| Cer_65p | Cer | (d18:0_16:0) | -0.95269429 | 1.714795646 |
| DG_311p | DG | (19:2e) | -0.98859524 | 0.822596719 |
| PE_3286p | PE | (16:1e_18:1) | -1.00580562 | 1.208568828 |
| MG_1733p | MG | (29:1) | -1.06342538 | 0.915584651 |
| LPE_280n | LPE | (17:0) | -1.06413034 | 1.19567151 |
| PE_3282p | PE | (22:0_12:2) | -1.07885542 | 1.594513786 |
| TG_4225p | TG | (15:0_14:0_16:1) | -1.08615664 | 2.159630319 |
| PE_3599p | PE | (40:7) | -1.10825289 | 0.943522679 |
| Cer_113p | Cer | (d18:1_24:1) | -1.11973924 | 1.789807522 |
| LPE_340n | LPE | (20:4) | -1.12151782 | 1.510455906 |
| PC_2935p | PC | (18:3_20:3) | -1.13358315 | 1.019090011 |
| SM_3814p | SM | (d36:1) | -1.13404797 | 2.965365937 |
| PE_3463p | PE | (18:0_20:4) | -1.19714647 | 0.743373361 |
| PC_573n | PC | (18:0_16:0) | -1.20163386 | 1.976754087 |
| TG_4712p | TG | (20:3_20:5_22:6) | -1.20903404 | 1.739802403 |
| PC_2686p | PC | (34:2e) | -1.2107671 | 1.363757171 |
| MePC_2259p | MePC | (38:9) | -1.22616911 | 1.201686538 |
| PC_3049p | PC | (18:4_22:4) | -1.23266076 | 2.816607788 |
| LdMePE_446n | LdMePE | (20:4) | -1.23562825 | 0.763388212 |
| LdMePE_377n | LdMePE | (16:0) | -1.24792751 | 1.561257113 |
| PC_2569p | PC | (20:0_14:1) | -1.25457283 | 1.526480587 |
| MePC_1996p | MePC | (33:1) | -1.31577587 | 1.579708951 |
| PC_2384p | PC | (8:1e_10:1) | -1.32192809 | 2.510672886 |
| PE_830n | PE | (36:2) | -1.3360492 | 0.929928487 |
| PE_685n | PE | (18:1e) | -1.35693454 | 1.541742036 |
| DG_475p | DG | (17:1_16:1) | -1.43295941 | 1.129588993 |
| PE_939n | PE | (18:0_20:4) | -1.43538614 | 1.474650601 |
| PE_950n | PE | (18:0e_20:4) | -1.44745898 | 1.412670233 |
| PA_506n | PA | (8:0e_10:0) | -1.45490196 | 1.059100768 |
| PC_2744p | PC | (36:2) | -1.45943162 | 1.280222073 |
| MG_1730p | MG | (29:1) | -1.51727569 | 0.915690563 |
| PE_3407p | PE | (16:0e_21:1) | -1.5187288 | 4.020422757 |
| PE_3621p | PE | (20:4_22:6) | -1.52356196 | 2.254267406 |
| Cer_21n | Cer | (d40:1) | -1.53373718 | 1.221083757 |
| DG_675p | DG | (19:0_18:2) | -1.53533173 | 2.644996111 |
| MePC_2272p | MePC | (39:4) | -1.56187889 | 2.450627692 |
| PC_2554p | PC | (20:1_14:0) | -1.56187889 | 1.477133465 |
| ChE_195p | ChE | (20:4) | -1.5849625 | 1.830768427 |
| LPC_1000p | LPC | (18:2) | -1.61143471 | 2.665049656 |
| PE_710n | PE | (20:2e) | -1.61325155 | 1.248581419 |
| MePC_2233p | MePC | (37:4e) | -1.65711229 | 1.287339851 |
| LdMePE_379n | LdMePE | (16:0) | -1.68314289 | 0.501251711 |
| PC_2388p | PC | (8:1e_10:1) | -1.81557543 | 2.420110134 |
| PE_3095p | PE | (16:0e) | -1.9068906 | 0.993061628 |
| PE_753n | PE | (16:0_18:2) | -1.94753258 | 2.259636761 |
| LPE_344n | LPE | (22:4) | -2.45285896 | 2.333376926 |
| DG_262p | DG | (16:1e) | -2.80735492 | 0.988940517 |
| PEt_1128n | PEt | (36:0e) | -2.91253716 | 2.634644645 |
